# Cisplatin neurotoxicity targets specific subpopulations and K^+^ channels in tyrosine-hydroxylase positive dorsal root ganglia neurons

**DOI:** 10.1101/2022.01.12.476091

**Authors:** Carrie J. Finno, Yingying Chen, Seojin Park, Jeong H. Lee, Cristina M. Perez-Flores, Ebenezer N. Yamoah

## Abstract

Among the features of cisplatin chemotherapy-induced peripheral neuropathy are chronic pain and innocuous mechanical hypersensitivity. The complete etiology of the latter remains unknown. Here, we show that cisplatin targets a heterogeneous population of tyrosine hydroxylase-positive (TH^+^) primary afferent dorsal root ganglion neurons (DRGNs) within the primary afferent dorsal root ganglia in mice, determined using single-cell transcriptome and electrophysiological analyses. TH^+^ DRGNs regulate innocuous mechanical sensation through C-low threshold mechanoreceptors. A differential assessment of wild-type and vitamin E deficient TH+ DRGNs revealed heterogeneity and specific functional phenotypes. The TH+ DRGNs comprise; fast-adapting eliciting one action potential (AP; 1-AP), moderately-adapting (≥ 2-APs), in responses to square-pulse current injection, and spontaneously firing (SF). Cisplatin increased the input resistance and AP frequency but reduced the temporal coding feature of 1-AP and ≥ 2-APs neurons. By contrast, cisplatin has no measurable effect on the SF neurons. Vitamin E reduced the cisplatin-mediated increased excitability, but did not improve the TH^+^ neuron temporal coding properties. Cisplatin mediates its effect by targeting outward K^+^ current, likely carried by through K2P18.1 *(Kcnk18)*, discovered through the differential transcriptome studies and heterologous expression. Studies show a potential new cellular target for chemotherapy-induced peripheral neuropathy and implicate the possible neuroprotective effects of vitamin E in cisplatin chemotherapy.

## Introduction

Cisplatin is one of the most efficacious chemotherapeutic drugs to treat several solid and blood cancers (Tsang et al., 2009). Still, it is wrought with peripheral neuropathic side effects that limit its chemotherapeutic value. The anti-cancer effects of cisplatin and related platinum compounds stem from the induction of intra- and inter-strand DNA cross-linker and denaturation of nuclear and mitochondrial DNA. The ensuing effects are necrotic and apoptotic cell death of cancer cells (Eastman and Barry, 1987; Dzagnidze et al., 2007). Puzzlingly, the underlying mechanism for cisplatin-mediated neurotoxicity remains unclear. Symptoms include painful paresthesia in the extremities and thermal and tactile allodynia or hyperalgesia (Hu et al., 2019b), with dorsal root ganglion neurons (DRGNs) serving as the unquestionable target. The diversity of neuropathic repercussions may stem from DRGNs’ heterogeneity in properties and functions and varying susceptibility to cisplatin toxicity. Studies have shown that cisplatin-mediated degeneration of large-myelinated DRGNs underlies patients’ peripheral neuropathy (Cavaletti et al., 1992; Krarup-Hansen et al., 1999; McDonald et al., 2005; Krarup-Hansen et al., 2007), but the etiology for exaggerated mechanosensitivity is unknown (Starobova and Vetter, 2017). The knowledge gap was one of the motivations for the current studies.

DRGNs are heterogeneous in size, with distinct neurochemistry, and can be grouped into as many as thirteen clusters based on their gene expression, subserving multiple sensory modalities (Usoskin et al., 2015; Li et al., 2016). Among the classes of DRGNs are tyrosine-hydroxylase positive (TH^+^) primary afferents, implicated for mechanical hypersensitivity in animal models of cisplatin chemotherapy-induced peripheral neuropathy (CIPN) (Draxler et al., 2014). The TH^+^ afferents represent unmyelinated C-low threshold mechanoreceptors (C-LTMRs). C-LTMRs regulate innocuous mechanical sensation across species and label as TH^+^ in mice, but not in humans (Brumovsky, 2016). However, recent comparative transcriptomic studies clearly demonstrate that TH^+^ DRGNs in mice correspond to C-LTMRs in non-human primates (Kupari et al., 2021) and humans (Tavares-Ferreira et al., 2021). C-LTMRs have been implicated in animal models of cisplatin chemotherapy-induced peripheral neuropathy (CIPN) (Li et al., 2011; Draxler et al., 2014). They are perceived to respond to innocuous mechanical stimulation, cooling sensations, and chemical-mediated pain hypersensitivity (Malmberg et al., 1997). Additionally, reports have suggested functional heterogeneity within murine TH^+^ DRGNs (Delfini et al., 2013; Gaillard et al., 2014), and unsupervised classification of DRGNs in mice has identified two molecular subsets of TH^+^ neurons; TH1 and TH2. The TH2 neurons were most altered with vitamin E deficiency (Finno et al., 2019). The importance of vitamin E in TH+ DRGNs in CIPN is emerging, underpinned by the findings that decreased plasma vitamin E levels increased susceptibility to peripheral neuropathy (Bove et al., 2001). The stark resemblance between the clinical and neuropathologic features of CIPN (Muller et al., 1983; Traber et al., 1987) and patients with peripheral neuropathy due to familial ataxia with vitamin E deficiency (AVED) (Gotoda et al., 1995; Yokota et al., 1996) raises the possibility that the two sensory deficits share a common underlying mechanism.

We tested the hypothesis that the heterogeneity in molecular features of TH^+^ DRGNs yields functional diversity, and the shared characteristics of CIPN and vitamin E deficiency can be used to identify therapeutic targets of the two sensory deficits. We show that TH^+^ DRGNs consist of at least three functionally distinct neuronal subtypes, two adapting and one spontaneously active neuronal class, with differential responses to the effects of cisplatin and vitamin E. We demonstrated that the neuroprotective effects of vitamin E on CIPN might be mediated through effects on a two-pore domain subfamily channel, K_2P_18.1 *(Kcnk18*), thus providing a potential therapeutic target for the treatment of sensory deficits associated with CIPN and vitamin E deficiency.

## Materials and Methods

### Single-cell RNA sequencing

In a previous study, single-cell RNA sequencing was performed on DRG cell culture from wild-type C57BL6/J mice and tocopherol transfer-alpha protein null mice (*Ttpa*^-/-^) to investigate the effect of vitamin E deficiency on DRGN gene expression (Finno et al., 2019). Mice were 5-6 months of age and consisted of 1 male and 1 female mouse from three experimental groups; *Ttpa*^+/+^ fed a regular diet (WT), *Ttpa*^-/-^ mice fed a vitamin E deficient diet (DEF), and *Ttpa*^-/-^ mice fed a highly vitamin E supplemented diet (SUPP). DRGNs were collected from C1 (cervical) level to L6 (lumbar), with an average of 3,614 ± 470 DRGN from two replicate mice per group. Barcoded 3’ single-cell libraries were prepared from single-cell solutions using the Chromium Single Cell 3’ Library and Gel bead kit vs2 (10X Genomics, Pleasanton, CA). Libraries were pooled and sequenced on an Illumina HiSeq4000 with pair-end 100 bp reads.

A total of 382 million reads was generated (17,862 ± 2,625 reads/cell). Cellranger v.2.0.1 and bcl2fastq v.2.17.1.14 commands mkfastq and count were used to generate fastq files per sample, align to mm10, filter, and perform barcode and UMI counting. Approximately 65% of reads mapped to the murine transcriptome, with an average of 1,788 ± 283 genes detected per cell, with no difference between experimental groups (Finno et al., 2019). Analyses were conducted in R, version 3.4.4 (Team, 2018). Normalization, clustering, and calculation of TSNE (van der Maaten and Hinton, 2008) coordinates were performed using Seurat, version 2.3.0 (Satija et al., 2018).

Unsupervised single-cell transcriptome profiling identified 14 initial subpopulations in all mice (Finno et al., 2019), using gene profiles as previously reported (Usoskin et al., 2015; Li et al., 2016). These clusters were classified as PEP1, PEP2, NP1, NP2, NP2-2, NP3, NF1, NF2, NF3, NF4-5, TH1, TH2, and “Unassigned” for the unassigned cluster (Finno et al., 2019). The identification of two distinct TH subgroups had not been previously reported. Despite highly overlapping transcriptional profiles between TH1 and TH2, these subtypes were too unrelated to merge (Finno et al., 2019).

### TH1 vs. TH2 DRGN single-cell RNA sequencing analysis

Two analyses were performed to investigate further the distinct profiles of TH1 vs. TH2 DRGNs in this sample set. The first analysis determined the number of genes unique to a single cluster when TH1 and TH2 subtypes were compared against all other DRGN subpopulations. This analysis was performed in Seurat using the FindAllMarkers option (Satija et al., 2018). The second analysis was to compare the only TH1 vs. TH2 transcriptional profiles, using a logfc. threshold of 0.25. For both analyses, subsequent pathway analyses were performed using Panther Pathway overrepresentation analysis (http://pantherdb.org/).

### Single-molecule fluorescence in situ hybridization (smFISH) with RNAscope

DRGNs were dissected and fixed in 4% PFA in DEPC-treated PBS for 2 hr at 4°C to preserve RNA Samples were washed with DEPC-treated PBS three times. DRG samples were sequentially dehydrated in 10%, 20%, and 30% sucrose solution at 4°C for 1 hr, 2 hr, and overnight, respectively, then embedded in OCT for cryosection. Samples were cryo-sectioned to a thickness of 10 μm, placed onto Superfrost slides, and stored at -80 °C until processed. Probe hybridization was performed according to the manufacturer’s instructions (Advanced Cell Diagnostics, ACD). Sections were immersed in pre-chilled 4% PFA for 15 min at 4°C. Sections were then dehydrated at RT in 50%, 70%, and twice in 100% ethanol for 5 min each and allowed to dry for 1-2 min. Fixation and dehydration were followed by protease digestion, using protease 4 for 30 min at RT Sections were then incubated at 40°C with the following solutions: 1) target probe in hybridization buffer A for 3 hrs; 2) preamplifier in hybridization buffer B for 30 mins; 3) amplifier in hybridization buffer B at 40°C for 15 mins; and 4) label probe in hybridization buffer C for 15 mins. After each hybridization step, slides were washed with washing buffer three times at RT. For fluorescent detection, the label probe was conjugated to Alexa Fluor 594. Probes, positive and blank negative controls were obtained from ACD. Sequences of the target probes, preamplifier, amplifier, and label probe are proprietary. Detailed information about the probe sequences can be obtained by signing a nondisclosure agreement provided by the manufacturer. Incubation in DAPI solution for 15s at RT was performed to label cell nuclei. Slides were then mounted in Fluoromount-G and sealed under a coverslip. Images were captured with a Nikon A1 and Olympus FV1000 confocal microscope. Dots in each fluorescent positive cell were counted and scored as described.

### Cell culture

Dorsal root ganglion neurons were isolated from the mouse TH-EGFP-positive (TH-EGFP+) transgenic mice, using a combination of enzymatic and mechanical procedures. Adult male and female (6-8 weeks old) TH-EGFP+ mice were euthanized, and the DRG were removed in a solution containing Minimum Essential Medium with HBSS (Invitrogen), 0.2 g/L kynurenic acid, 10 mM MgCl_2_, 2% fetal bovine serum (FBS; v/v), and 6 g/L glucose. The central DRG tissue was dissected and digested in an enzyme mixture containing 1 mg/ml collagenase type I and 1 mg/ml DNase at 37°C for 15 min. After a series of gentle triturations and centrifugation in 0.45 M sucrose, the cell pellets were reconstituted in 900 μl of culture medium (Neurobasal-A, supplemented with 2% B27 (v/v), 0.5 mM L-glutamine, and 100 U/ml penicillin; Invitrogen), and filtered through a 40 μm cell strainer for cell culture and electrophysiological experiments. For adequate voltage-clamp and satisfactory electrophysiological experiments, we cultured DRGNs for ∼24 hr. All electrophysiological experiments were performed at room temperature (RT; 21–22°C).

### Electrophysiology

Patch-clamp experiments were performed in standard whole-cell using an Axopatch 200B amplifier (Axon Instruments). Patch pipettes had a resistance of 2–3 MΩ when filled with an internal solution consisting of (in mM): 146 KCl, 10 HEPES, 5 EGTA, 1 CaCl_2_, 1 MgCl_2_, and 4 MgATP and 0.4 Na_2_GTP (pH 7.3 with KOH). The external solution contained (in mM): 145 NaCl, 6 KCl, 10 HEPES, 1 MgCl_2_, 2 CaCl_2_, and 10 glucose (pH 7.3 with NaOH). All experiments were done at RT Currents were sampled at 50 or 20 kHz and filtered at 5 or 2 kHz. Voltages were not corrected for a liquid junction potential.

### Current and voltage-clamp experiments

Whole-cell membrane potential recordings were performed using an Axopatch 200B amplifier (Molecular Devices, Sunnyvale, CA). Membrane potentials were amplified, bandpass filtered (2-10 kHz), and digitized at 5-50 kHz using an analog-to-digital converter (Digidata 1200, Molecular Devices) as described earlier (Levic et al., 2007; Rodriguez-Contreras et al., 2008). Electrodes (2-3 MΩ) were pulled from borosilicate glass pipettes, and the tips were fire-polished. The normal extracellular/bath solution consisted of (in mM) 130 NaCl, 5 KCl, 1 MgCl_2_, 2 CaCl_2_, 10 D-glucose, and 10 4-(2-hydroxyethyl)-1-piperazineethanesulfonic acid (HEPES), pH 7.3. The normal internal/pipette/ solution contained (in mM) 132 KCl, 1 MgCl_2_, 0.01 CaCl_2_, 2 ethylene glycol-bis(β-aminoethyl ether)-N,N,N′,N′-tetraacetic acid (EGTA) 5 ATP-K_2_, and 10 HEPES, pH 7.3. The seal resistance was typically 5-10 GΩ. Capacitance and series resistance compensation (>90%) were made, and traces were filtered at 2 kHz using an 8-pole Bessel filter and sampled at 5 kHz. The liquid junction potentials (LJP) were measured (2.1 ± 1.2 mV; n = 87) and corrected as described previously (Rodriguez-Contreras and Yamoah, 2001). Data analyses were performed using the pClamp and Origin software (MicroCal Inc., Northampton, MA). Where appropriate, pooled data are presented as mean <u>⩲</u> SD.

Whole-cell voltage-clamp recordings were conducted at RT, using an Axopatch 200B amplifier and filtered at 2 kHz through a low-pass Bessel filter. Data were digitized at 0.5-1.0 kHz using a Digi-data analog-to-digital converter. DRGNs, after 1-2 days in culture, was held in a bath solution (in mM; 4 KCl, 2 MgCl_2_, 0.1 CaCl_2_, 10 HEPES, 10 D-glucose, 75 NaCl, 50 N-methyl glucamine, 20 tetraethylammonium chloride (TEACl), and 5 4-Aminopyridine (4-AP)). The bath solution composition curbed K^+^ currents. We suppressed Ca^2+^ and Cl^-^ currents by using 0.05 mM bath Ca^2+^ and replacing 90 mM bath Cl^-^ and pipette solution with aspartate. Replacing Cl^-^ with aspartate increased the LJP by 9.5 ± 1.8 mV (n = 36), which was corrected. Borosilicate glass pipettes were pulled using a Sutter P-97 Flaming-Brown micropipette puller (Sutter Instruments) and fire-polished for an optimal pipette resistance (2-3Ω). Pipettes were backfilled with an internal solution (in mM; 2.5 EGTA, 95 NMGCl, 5 NaCl, 10 CsCl, 2 MgCl_2_, 0.1 CaCl_2_, 10 HEPES, 5 ATP-bis(tris), and 5 glutathione). Data were considered for analyses if the seal resistance was greater than 2GΩ and access resistance was less than 7 MΩ.

After achieving a gigaohm seal, gentle suction was applied to form a whole-cell configuration. A holding potential of -110 mV was applied to all protocols to prime Na^+^ currents for activation. Activation was analyzed with depolarizing voltage steps from -70 to 60 mV, using a ΔV of 2.5 mV. Inactivation was studied using a standard protocol and range of pre-pulse from -110 to 60 mV, using a ΔV of 5 mV. All protocols employed p/n (n = 7) leak subtraction at the holding potential.

### Statistical analyses

Where appropriate, pooled data are presented as mean ± SD. Significant differences between groups were tested using t-test and ANOVA, where applicable. The null hypothesis was rejected when the two-tailed *p*-value < 0.05 is indicated with *, < 0.01 with **, < with ***. The number of mice and neurons are reported as *n*.

### Study Approval

Animals were housed and cared for under the University of California Davis (UCD) and University of Reno (UNR) standing committee on animal use and care (IACUC) as well as the Guide for the Care and Use of Laboratory animals (8^th^ edition, 2011). All procedures performed were also approved by the University (UCD and UNR.) IACUC.

## Results

### Transcriptomic profiling of murine TH^+^ DRGN subpopulations

A previous single-cell RNA-sequencing study to evaluate the effect of vitamin E on DRGN gene expression revealed distinct TH^+^ neurons (Finno et al., 2019). Despite highly overlapping transcriptional profiles within the TH^+^ DRGN group, at least two subtypes were not closely related enough to merge, resulting in additional distinction into TH1 and TH2 subgroups. In particular, profound alterations with vitamin E deficiency were identified in the TH2 subpopulation (Finno et al., 2019). We hypothesize that distinct molecular identity may yield diverse functional phenotypes amongst the TH^+^ DRGNs, that albeit, likely confer varied pharmacology. We determined the number of genes unique to a single cluster when TH1 and TH2 groups were compared against other DRGN subpopulations. Within the TH1 subpopulation, 102 uniquely expressed genes were identified. However, 254 unique genes were identified in the TH2 subpopulation (**Table S1**). The top five transcripts defining each DRGN subpopulation are represented in **Fig. 1A**. Upregulation of voltage-gated Ca^2+^ and K^+^ channels have been detected with vitamin E deficiency, specifically in the TH2 subpopulation, including Ca_v_2.3 and the Ca^2+-^activated K^+^ channels, Kcnmb1 and Kcnmb2 (Finno et al., 2019). The remaining K^+^ channels significantly upregulated with vitamin E deficiency in TH^+^ DRGNs were the K^+^ two-pore (K_2P_) domain subfamily channels (*Kcnk3, Kcnk12*, and *Kcnk18*). Of these, *Kcnk18* was differentially expressed across five DRGN subpopulations (**Fig. 1B**), with more significant upregulation in TH2 neurons (p=4.74 × 10^-5^) than TH1 (p=0.01). This channel was thus prioritized for further investigation.

**Figure 1.**
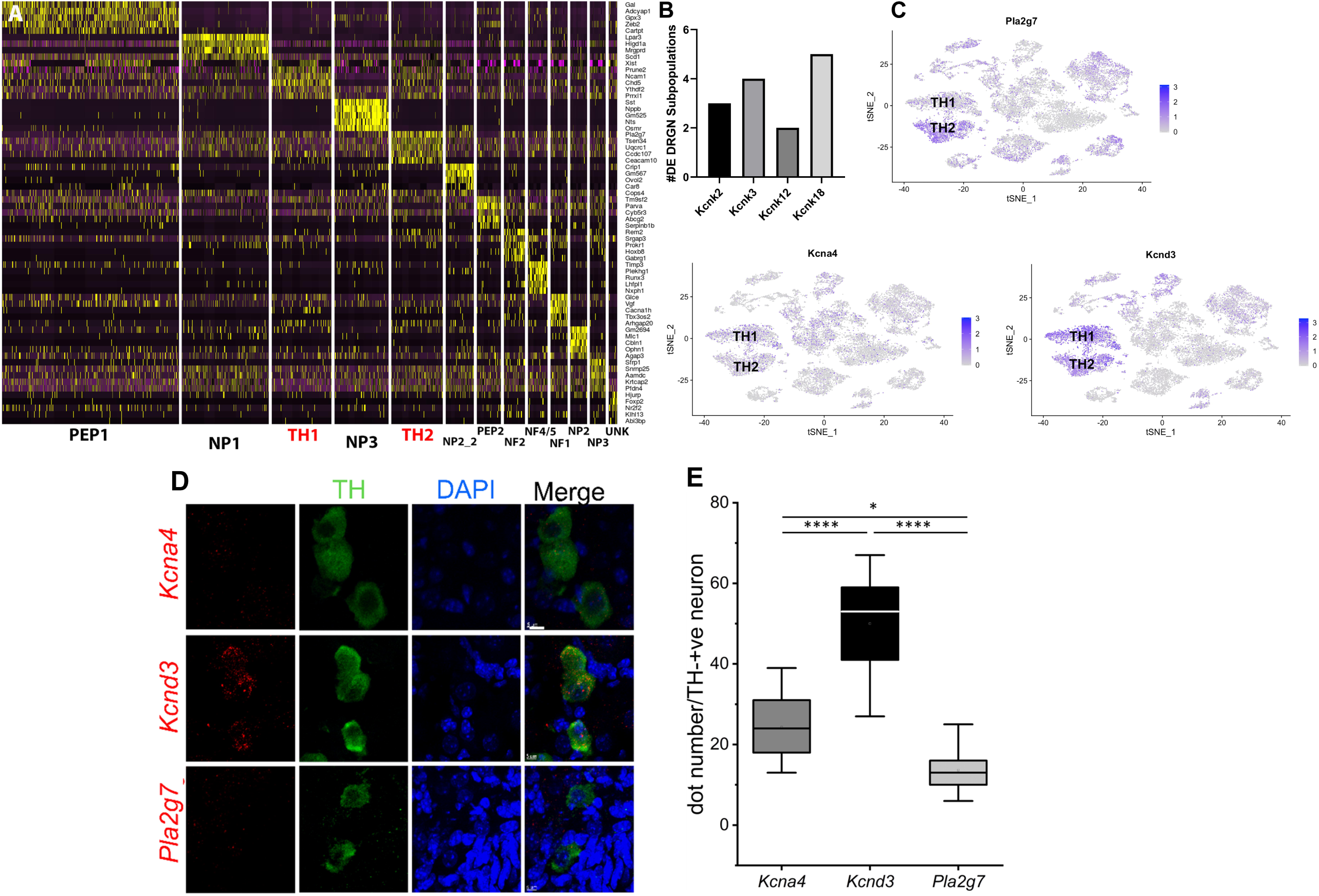
TH1 and TH2 are distinct subpopulations of TH^+^ DRGNs of adult mice. **A**. Heat map of the top five unique genes defined each DRGN subpopulation in a previously published study (Finno et al., 2019). TH^+^ DRGNs (TH1 and TH2; red) are distinct subpopulations based on transcriptional profiles. **B**. Differential transcript expression of K^+^ two-pore domain subfamily K channels (*Kcnk2, Kcnk3, Kcnk12*, and *Kcnk18*) across the DRGN subpopulations, with *Kcnk18*, differentially expressed across five subpopulations (PEP1, PEP2, TH1, TH2, UNK). **C**. TSNE plots demonstrating overall expression of *Pla2g*7 (TH2>TH1), *Kcna4* (TH1>TH2), and *Kcnd3* (TH1>TH2) from single-cell RNA-sequencing (Finno et al., 2019). **D**. Single-molecule fluorescence in situ hybridization (smFISH) with RNAscope from TH-EGFP+ transgenic mice, demonstrating expression of *Kcna4, Kcnd3*, and *Pla2g7* in TH^+^ DRGNs. Green: TH^+^ DRGN, Blue: DAPI nuclear stain, Red: *Kcna4* (first row), *Kcnd3* (second row), and *Pla2g7* (third row) mRNA. Scale bars represent 5 μm. **E**. Quantification of mRNA using RNAscope for *Kcna4, Kcnd3, and Pla2g7*. Within TH^+^ DRGNs, *Kcnd3* was the most highly expressed. Mean + SD, N = 11 counts per experimental group, one-way ANOVA, or Kruskal-Wallis. Compared to GAPDH (not shown) one-way ANOVA, *p< 0.05, **p< 0.01, ***p< 0.001 ****p<0.0001. DE=differentially expressed, NF=neurofilament, NP=non-peptidergic, PEP=peptidergic, TH=tyrosine hydroxylase, UN=unknown cluster, *n*=2 mice per group with ∼3,600 cells/mouse profiled.

The top TH2 unique transcript, *phospholipase A2 group 7* (*Pla2g7)*, was overrepresented in the TH2 subpopulation when evaluating an expression using the TSNE plot (**Fig. 1C**). For the TH1 subpopulation, the top transcripts included peripherin (*Prph*) and ribosomal proteins, which were not very specific when evaluating the TSNE plot (**Fig. S1**). Of the differentially expressed K^+^ channels, K_v_1.4 (*Kcna4*) significantly defined the TH1 subpopulation (p=0.0001; **Fig 1C**). Pathway analysis for the unique genes in the TH1 subpopulation did not identify any statistically significant pathways (P_FDR_<0.05). In contrast, a 10.07-fold enrichment of the ubiquitin-proteasome pathway (P_FDR_ = 0.007) was placed in the TH2 subpopulation.

The second analysis compared the only TH1 vs. TH2 transcriptional profiles. A total of 1052 differentially expressed transcripts were identified, with 310 higher in TH1 and 742 higher in TH2 (**Table S2**). Of the differentially expressed K^+^ channels, K_v_4.3 (*Kcnd3*) was significantly increased in the TH1 vs. TH2 subpopulation (p=6.17 × 10^-21^; **Fig 1C**). The 310 transcripts that were higher in TH1 DRGNs corresponded to 18 upregulated pathways, including hedgehog signalling, the three metabotropic glutamate receptor pathway, and many G-protein-related receptor signalling pathways (**Fig. S2A**). The 742 transcripts that were higher in TH2 DRGNs were associated with two upregulated pathways, Parkinson’s disease (P_FDR_ = 1.02 × 10^-7^) and the ubiquitin-proteasome pathway (P_FDR_ = 0.003) (**Fig. S2B**). To validate these top differentially expressed transcripts (*Pla2g7, Kcna4*, and *Kcnd3*) in TH^+^ DRGNs, RNA-scope was performed (**Fig. 1D**). All transcripts were expressed in TH^+^ DRGNs, with *Kcnd3* expressed highest (**Fig. 1E**). These results support the distinction of at least two TH+ subpopulations of DRGNs in adult mice and prioritized three K^+^ channels, K_2P_18.1 (*Kcnk18*), K_v_1.4 (*Kcna4*), and K_v_4.3 (*Kcnd3*), for further experiments.

### Electrophysiological recordings of TH^+^ DRGNs

To test the hypothesis that the molecular heterogeneity in TH^+^ DRGNs yield functional diversity, we examined their response properties using TH-EGFP-positive (TH-EGFP+) transgenic mice. Data were obtained from isolated TH-EGFP+ DRGNs from adult (6-8-week-old) male and female mice. Three functional subtypes of TH^+^ DRGNs were identified (**Fig. 2**). The first subtype included fast-adapting TH^+^ DRNGs that elicited one action potential (1-AP) upon current injection (**Fig. 2A**). The second included moderately adapting TH^+^ DRGNs that produced greater than two APs (≥ 2-APs) upon current injection (**Fig. 2B**). The third subtype was spontaneously firing (SF) TH^+^ DRGNs (**Fig. 2C**). The three functionally distinct TH^+^ DRGNs showed stark differences in membrane input resistances and AP properties, sufficient to consider them separate (see **Fig. 2** legend). To further define the three classes of TH^+^ DRGNs, we switched from current-to voltage-clamp and measured the underlying whole-cell K^+^ current in the TH^+^ DRGN subtypes (**Fig. 2D-F**). Held at -90 and 40 mV and stepped from -100 mV to 40 mV using 10-mV increments, the yielding outward currents varied across the TH^+^ DRGNs (**Fig. 2G-H**). Assessment of inward current was not feasible under the experimental configurations.

**Figure 2.**
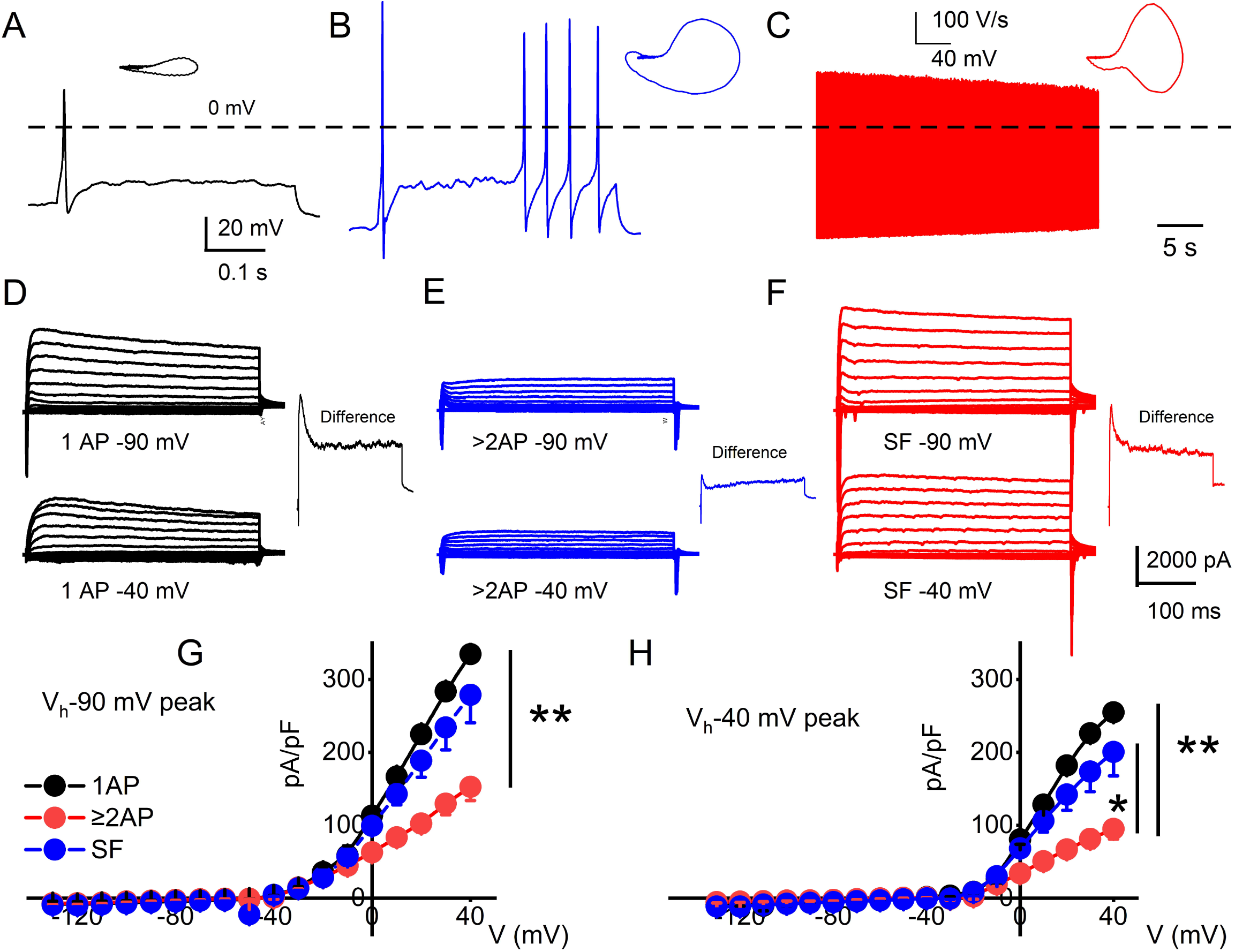
Functional differences in response properties of TH^+^ DRGNs. **A-C**. Representative AP traces were recorded from TH^+^ DRGNs. **A**. Fast-adapting TH^+^ DRGNs elicit 1-AP upon current injection (0.2 nA for the example shown). **B**. Moderately-adapting TH^+^ DRGNs, eliciting ≥ 2-APs (0.2 nA). **C**. Spontaneously firing (SF) TH^+^ DRGNs. **D-F**. Whole-cell K^+^ currents in 1-AP (**D**), ≥ 2-APs (**E**), and SF (**F**) TH^+^ DRGNs. Voltage-clamp recordings of TH^+^ DRGNs following current-clamp assessment. TH^+^ DRGNs were held at -90, and -40 mV stepped from -100 to 40 mV using 10-mV increments. The insets show a difference current trace using 40-mV stepped potential between neurons held at -90 and -40 mV. **G-H**. Summary data for the current-voltage (I/V) relationship showing differences in the current densities between the three classes of TH^+^ DRGNs. TH^+^ DRGNs with 1-AP (in black symbols), ≥ 2-APs (in blue symbols), and SF (in red symbols). (* p < 0.05; ** p < 0.01; n = 15 neurons from 4 mice).

We used oscillatory current injections at 0.4, 5, and 10 Hz as a proxy for the frequency regime of natural stimuli to evaluate the response properties of identified TH^+^ DRGNs (**Fig. 3A-B**). In response to sinusoidal current, the spike frequencies were enhanced as stimulus frequency increased in the fast-adapting 1-AP-neurons, reaching a saturating response at >10 Hz (**Fig. 3C**). For moderately-adapting ≥ 2-APs-neurons, the spike frequency declined, attaining an asymptotic level at >20 Hz (**Fig. 3C**). The vector strength measures the degree of synchronization between the stimulus and response. A value of 1 refers to perfect phase synchrony, and 0 refers to a random relation between stimulus and response (Goldberg and Brown, 1969). The computed vector strength for the 1-AP-was consistently greater than ≥ 2-APs-neurons (cf., a caveat in analyses, **Fig. 3** legend). Vector strength reduced as the stimulation frequency increased above 20 Hz for the two classes of neurons. However, invariably, the 1-AP-neurons had higher vector strength, indicating enhanced temporal coding (**Fig. 3D**). Results support distinct functional roles for three subtypes of TH^+^ DRGNs; 1-AP, ≥ 2-APs, and SF.

**Figure 3.**
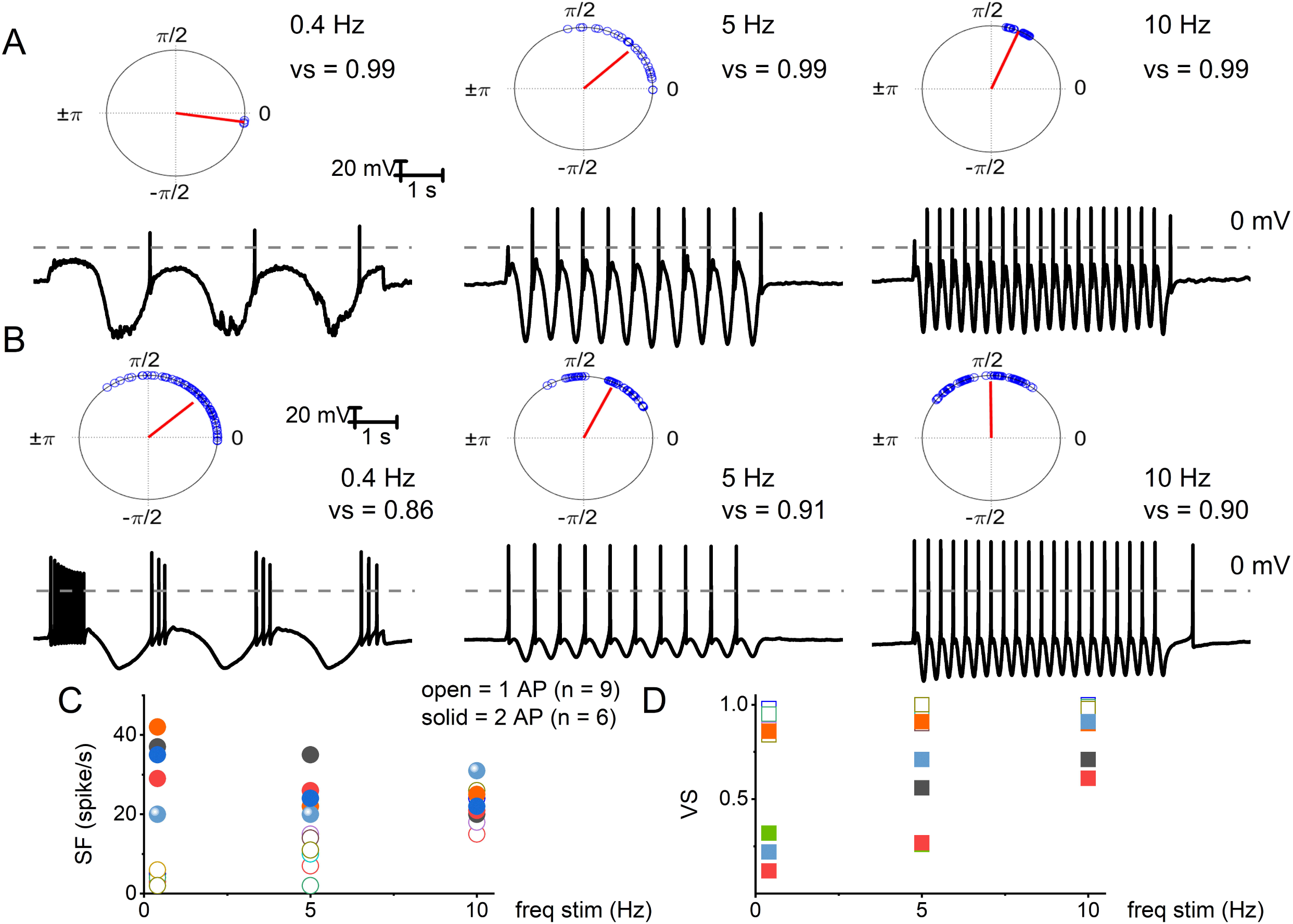
Electrical stimulation of TH^+^ DRGNs with oscillatory current injections. **A**. Upper panel: polar plots for a TH^+^ DRGNs with a 1-AP-elicited square pulse. Below are representative membrane responses of TH^+^ DRGNs to oscillatory current injection (0.2 nA) at 0.4, 5, and 10 Hz. **B**. Row of similar polar plots derived from TH^+^ DRGNs responding with ≥ 2-APs after square pulse injection (0.2 nA) to 0.4, 5, and 10 Hz oscillatory currents. **C-D**. Summary data of the two sets of TH^+^ DRGNs, showing the relationship between oscillatory current injection (0.4, 5, and 10 Hz, 0.2 nA) and spike frequency (solid and open symbols represent data from neurons with 1 AP, and ≥ 2 AP response features). **D**. Data were obtained from the same neurons as in **C**, with computed vector strength (VS). 1-AP TH^+^ DRGNs consistently showed higher VS, indicating better temporal coding. Each symbol represents a different TH^+^ DRGNs (n = 9 for 1-AP, n = 6 for ≥ 2-APs TH^+^ DRGNs).

### Cisplatin and vitamin E-mediated alterations of TH^+^ DRGNs and potential K^+^ channel target

The TH^+^ DRGNs are mechanically sensitive and are subject to profound transcriptional modifications in response to vitamin E deficiency (Finno et al., 2019). We surmised that cisplatin might target TH^+^ DRGNs stemming from the drug’s tactile hyperalgesia side effects (Hu et al., 2019b). As exemplified in **Fig. 4A**, cisplatin had minimal impact on the membrane input resistance of SF TH^+^ DRGNs. By contrast, cisplatin-induced a time-dependent membrane input resistance increase in the 1-AP- and ≥ 2-AP neurons (**Fig. 4A**). The effects of cisplatin can be seen in the expected increase in spike frequency and membrane excitability in TH^+^ DRGNs, except for the SF neurons (**Fig. 4B**). Whole-cell currents from 1-AP and ≥ 2-APs TH^+^ DRGNs showed cisplatin-mediated outward current reduction (**Fig. 4C**). For outward currents elicited from -70 mV to 0 mV, cisplatin reduced the current by ∼13% for 1-AP TH^+^ DRGNs and ∼30% for ≥ 2-APs TH^+^ DRGNs (**Fig. 4C**). When a sinusoidal current was injected into a 1-AP TH^+^ DRGN, spike frequency increased (p<0.001), and the vector strength decreased (p<0.001) after application cisplatin (**Fig. 4D**). The differential effects of cisplatin on the TH^+^ DRGN subtypes substantiated the assertion that the neurons may be functionally and pharmacologically different despite a common distinct transcriptional biomarker.

**Figure 4.**
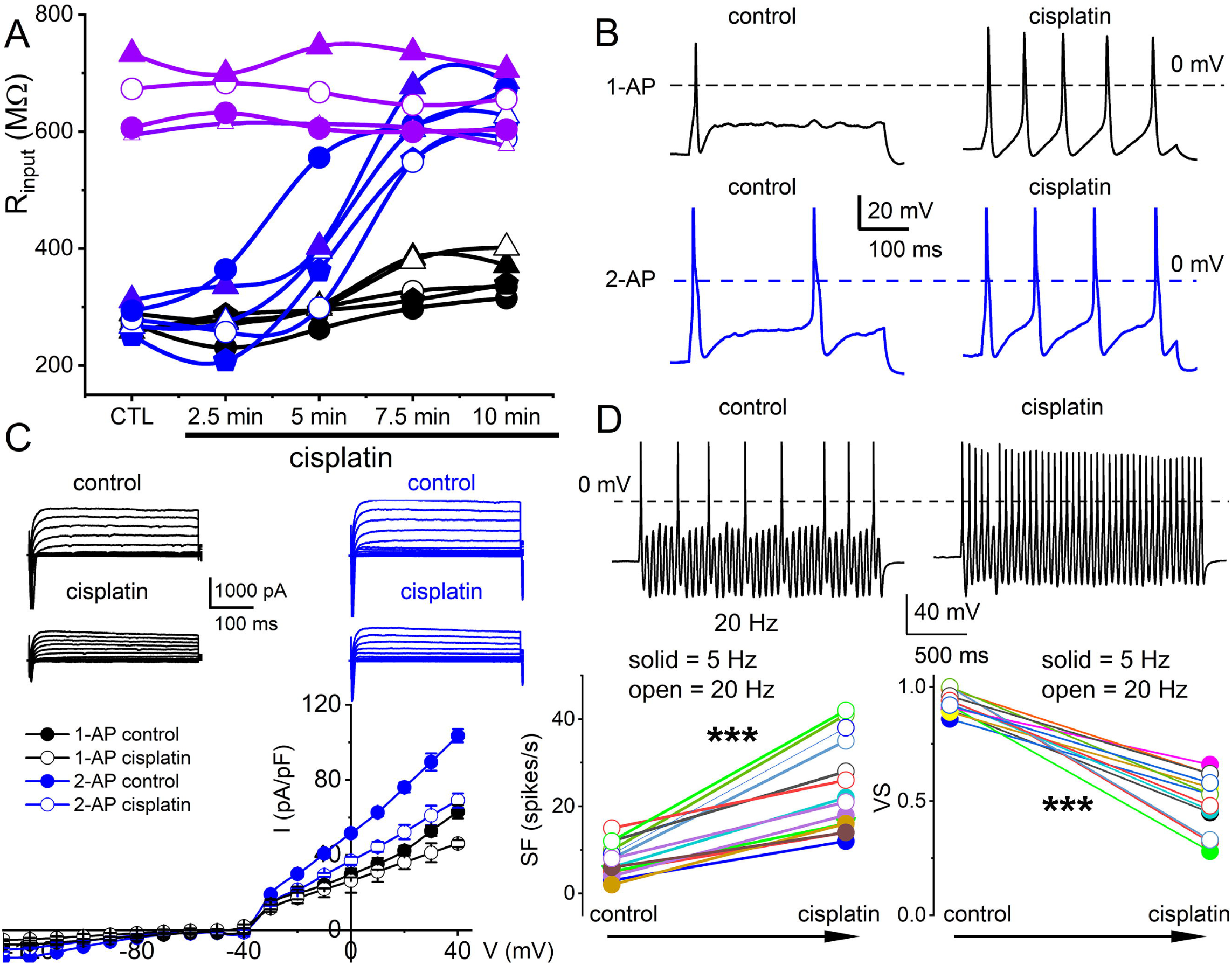
Cisplatin differentially increased membrane excitability of TH^+^ DRGNs but reduced their temporal coding properties. **A**. Time-dependent alterations of the input resistance of TH^+^ DRGNs after application of 2 μM cisplatin. Black and blue symbols represent TH^+^ DRGNs that elicit 1-AP (n = 5) and ≥ 2-APs (n = 5) upon current injection. In purple are spontaneously firing neurons (n = 4). **B**. Exemplary effects of cisplatin on rectangular-shaped current-injected APs on two distinct TH^+^ DRGNs (1-AP (upper, shown in black traces, and ≥ 2-APs (lower, shown in blue traces, panels). **C**. Whole-cell currents from 1-AP (black) and ≥ 2-APs TH^+^ DRGNs (blue) show cisplatin-mediated outward currents reduction. The lower panel shows the corresponding current density (pA/pF) and voltage relation (n = 9). For outward currents elicited from -70 mV to 0 mV, cisplatin reduced the current by ∼13% for 1-AP TH^+^ DRGNs and ∼30% for ≥ 2-APs TH^+^ DRGNs. **D**. Representative response properties of 1-AP TH^+^ DRGNs generated by simultaneous sinusoidal current injection (0.2 nA, 20 Hz). The amplitude criterion of a valid AP was 0 mV overshoot (dashed line). The lower panels summarize spike frequencies and vector strength (VS) changes before and after cisplatin (1 μM) application. Data were collected using 5 Hz (n = 8) and 20 Hz (n = 7) sinusoidal current. (*** p < 0.001).

Because vitamin E has specific marked effects on TH^+^ DRGNs (Finno et al., 2019), we examined vitamin E-mediated alterations of cisplatin-induced excitability, focusing on 1-AP-neurons. The addition of cisplatin significantly increased the input resistance of 1-AP TH^+^ DRGNs (p = 3.75 × 10^-4^), and the subsequent application of vitamin E reduced the input resistance to baseline levels (p = 2.32 × 10^-2^, **Fig. 5A**). The application of vitamin E reduced the cisplatin-induced increased excitability of 1-AP TH^+^ DRGNs (**Fig. 5B-D**) but did not improve the vector strength (**Fig. 5E**). Results suggest that vitamin E reduces the cisplatin-mediated increase in TH^+^ DRGN excitability but does not improve the temporal coding properties.

**Figure 5.**
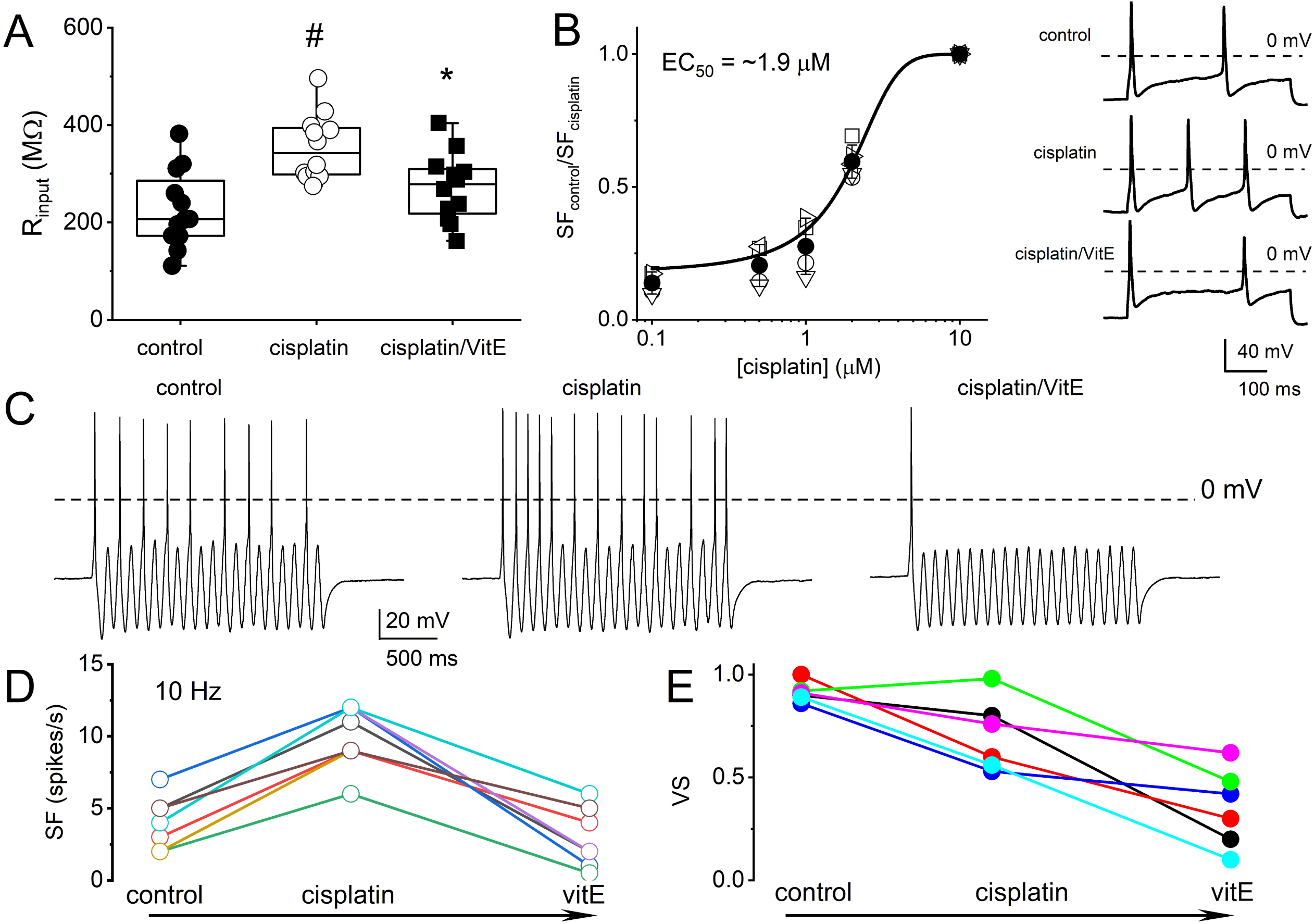
Vitamin E reduces cisplatin-mediated increased excitability of TH^+^ DRGNs but does not improve the coding properties. **A**. Summary of the effects of cisplatin on the input resistance of TH^+^ DRGNs after application of 1.5 μM cisplatin and the subsequent effects of the application of vitamin E (vitE, 100 μM) and 1.5 μM cisplatin. Differences between control and the treatment were tested using one-way ANOVA. #p = 3.75 × 10^-4^ and *p = 2.32 × 10^-2^ (n = 12). **B**. Dose-response relation on cisplatin-mediated increased spike frequency and applied cisplatin concentrations. A Hill coefficient of 2 and EC50 of 1.9 ± 0.3 μM (n = 4) was estimated. The inset on the right shows the effects of 0.5 μM cisplatin. **C**. Effects of cisplatin (0.5 μM) and vitE (100 μM) on the response properties of TH^+^ DRGNs generated by injecting sinusoidal current (0.2 nA, 10 Hz). **D**. The relation between spike-frequency in control, cisplatin and after cisplatin/vitE application (n = 6). Data were assessed from TH^+^ DRGNs that generated 1-AP in response to a square pulse. **E**. The corresponding computed VS, cisplatin, and vitE reduce the VS (n = 6).

To determine the properties of the underlying current responsible for cisplatin-mediated effects on 1-AP TH+ DRGNs, we examine the resting membrane potential in control and following cisplatin application. For 29 DRGNs examined, cisplatin produced 6 <u>⩲</u>2 mV depolarizing shift in the resting membrane potential (**Fig. 6A-B**). The rheobase current was reduced as summarized in **Fig. 6B**, suggesting that the underlying current may be partially operating at rest. Analyses of the threshold of all-or-none AP in control and upon application of cisplatin revealed reduced activation threshold in cisplatin (**Fig. 6C-E**). We predicted that the cisplatin-mediated effect likely is active at rest. We switched to voltage-clamped mode and evaluated the effects of cisplatin on the holding current at -70 mV holding potential. Consistent with the expectation, cisplatin reduced the holding current, and applying vitamin E reduced the cisplatin effects on the holding current and the ramp-voltage protocol-induced current, substantiating the prediction that a baseline “leak” current may be blocked by cisplatin (**Fig. 6F-G**).

**Figure 6.**
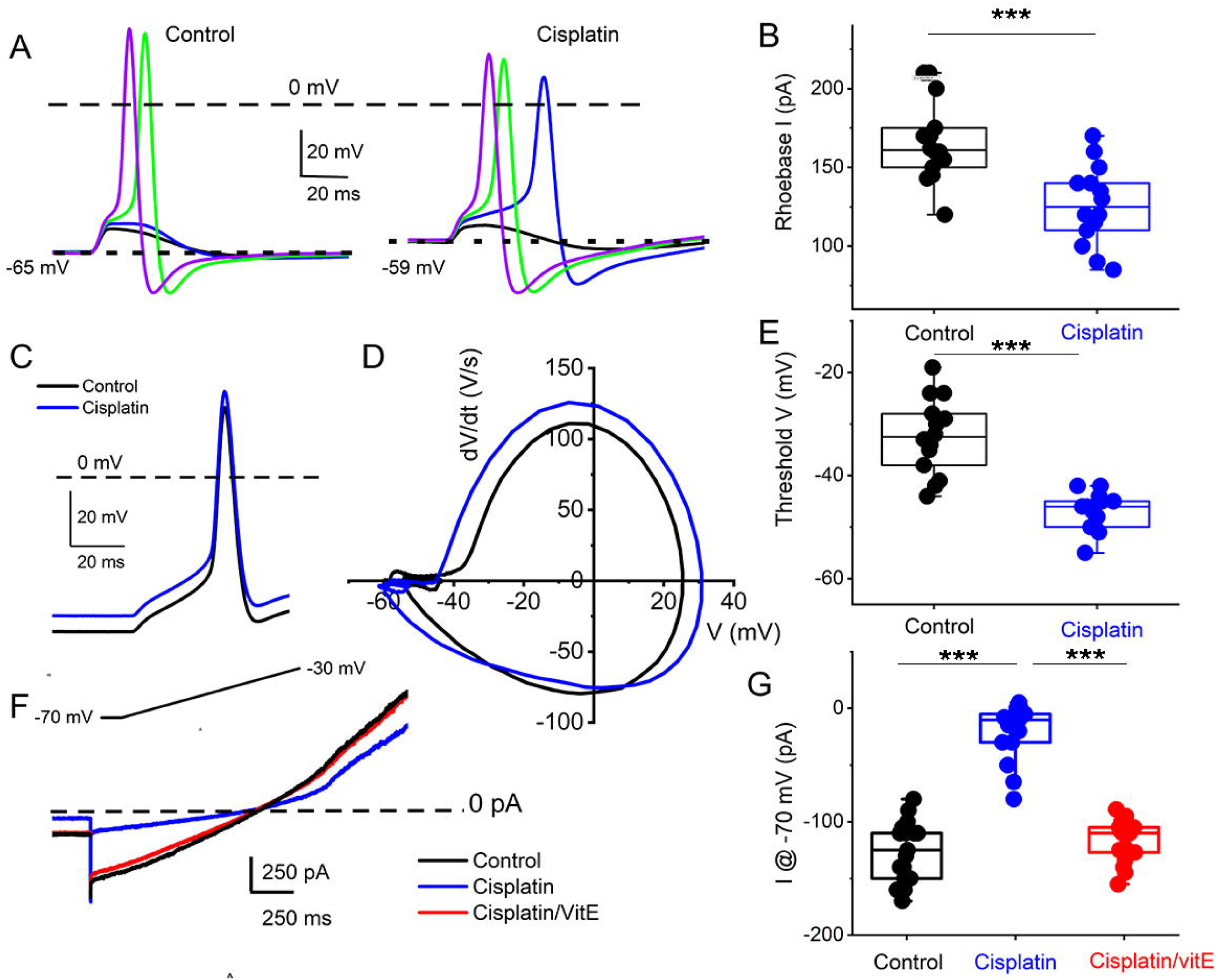
Cisplatin reduced induced membrane potential depolarization and reduced action potential threshold by suppressing a resting current. **A**. Left panel. Membrane depolarizations and action potentials were generated using a 5-ms current injection of different amplitudes. The strategy was used to determine the action potential threshold before and after applying 1 μM cisplatin (right panel). The dashed line indicates the 0-mV level, and the dotted line represents the resting membrane potentials. Note that cisplatin mediated ∼5-8-mV membrane depolarization. **B**. Average and raw data of the effective rheobase evaluated from **A** plotted for control and after application of cisplatin (1 μM). *p* values for statistical comparison are shown, and statistical significance is indicated with an asterisk (***p<0.001, n = 14 neurons from 3 mice). **C**. Exemplary action potentials were generated using 150 pA current injection for control (in black) and after cisplatin application (in blue). **D**. Phase plot (dV/dt) of the action potentials in **C** comparing controls (in black) and cisplatin effect (in blue). The membrane voltage threshold is determined from the phase plots for control and after cisplatin application. Cisplatin reduced the threshold voltage significantly (***p<0.001) as summarized in **E** (n = 13 from 3 mice). **F**. Current traces showing generated with a voltage-ramp from -70 mV holding voltage to -30 mV for control (black) and in the presence of cisplatin (1 μM, blue) followed by application of solution containing 1 μM cisplatin and 100 μM vitE (red). Cisplatin reduced the holding current, which was reversed with vitE application. Summary of the measured holding current at -70 mV holding voltage in control, after cisplatin and vitE (***p<0.001, n = 16 from 4 mice).

K^+^ channel transcripts altered in TH^+^ DRGNs in vitamin E deficient mice are *Kcnd3, Kcna4, Kcnk18* (Finno et al., 2019), which encode for Kv4.3, Kv1.4, and K_2P_18.1 channels, respectively. We expressed human K_v_1.4 and K_2P_18.1 plasmids in the human embryonic kidney (HEK) 293 cell line to identify potential K^+^ channels responsible for altering the outward current cisplatin-mediated effects. The cisplatin-sensitive current’s kinetic was non-inactivating and active at the resting membrane potential (**Fig. 5B, Fig. 6**). We excluded the Kv4.3 channel from the list because the current shows inactivation kinetics, and it is primarily a cardiac ion channel (Dixon et al., 1996; Wu et al., 2010). HEK 293 cells transfected with K_2P_18.1, and Kv1.4 plasmids yielded outward currents compared to non-transfected cells. Transfected cells were assessed with and without cisplatin before and after pre-incubation with vitamin E. While cisplatin reduced the current density in K_2P_18.1 and K_v_1.4 transfected HEK 293 cells (**Fig. 7**), pre-incubation with vitamin E only abolished the effects of cisplatin on the K_2P_18.1 current (K_2P_18.1; *p* <0.001 and K_v_1.4; p = 0.94 **Fig. 7C and F**, respectively). The neuroprotective effects of vitamin E on cisplatin-induced reduction of outward currents may be mediated through the K_2P_18.1 channel.

**Figure 7.**
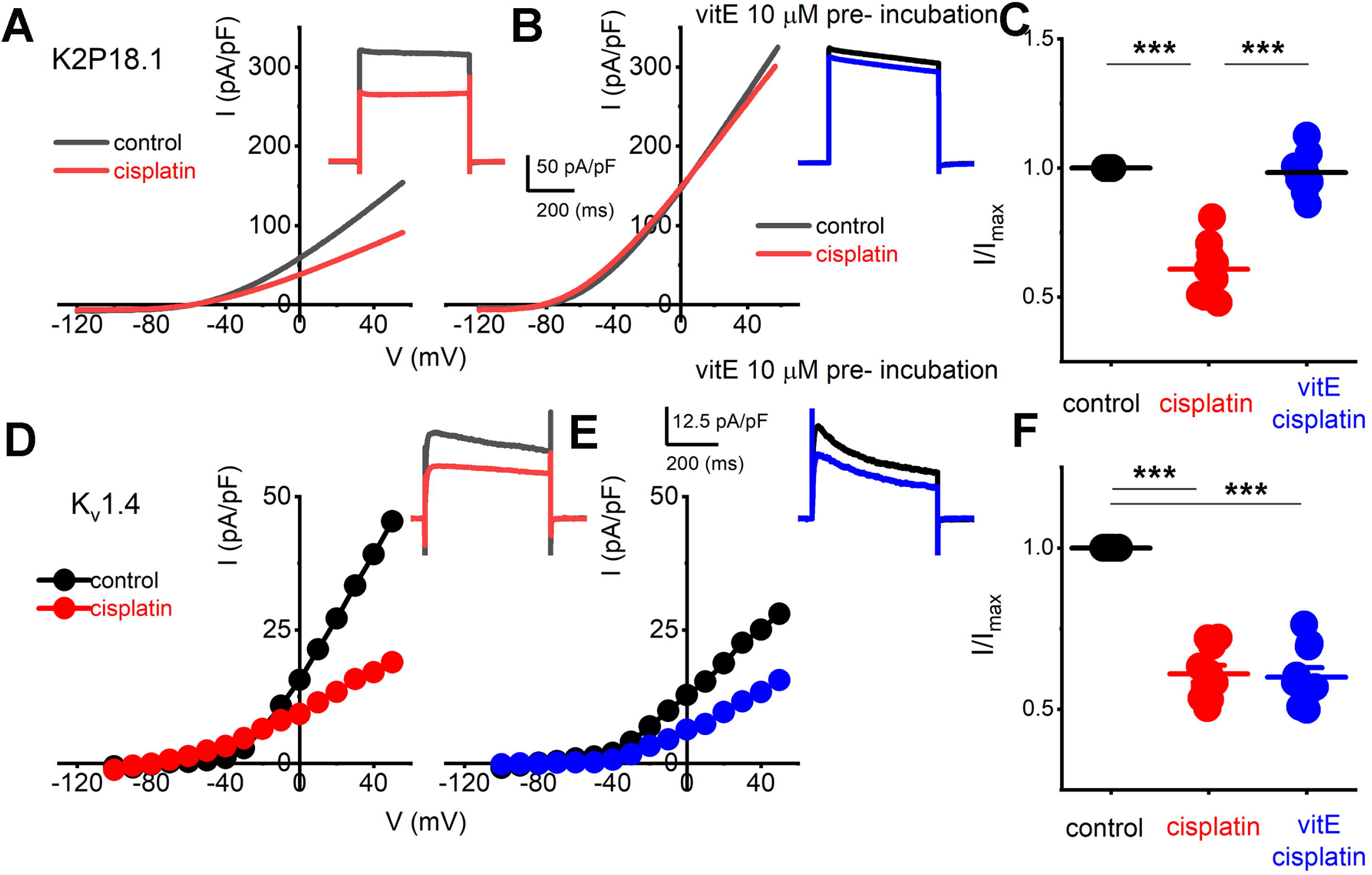
Effects of vitE pre-incubation on cisplatin-mediated alterations on expressed K_2P_18.1 and K_v_1.4 in HEK 293 cells. Outward K^+^ currents from HEK 293 cells transfected with the mouse K_2P_18.1 and K_v_1.4 plasmids. **A**. K_2P_18.1 channel expression; the recordings were obtained from a holding potential of -80 mV and stepped to -100 mV with a ramp to 60 mV within 500 ms (in black) or a square pulse ranging from -70 to 60 mV (ΔV = 10 mV). The insets show square-pulse elicited current. The red- square and ramp traces depict the effects of cisplatin application (10 μm). Cisplatin (10 μM) blocked ∼40% of the current. **B**. In contrast, pre-incubation (∼2 hrs) in 10-μM vitE sufficed to abolish the effects of cisplatin on K_2P_18.1 current (blue). I-V relationship in response to a ramp stimulus. **C**. Summary data of effects of cisplatin and after vitE pre-incubation obtained from n = 14 cells. **D-F**. Outward K^+^ currents elicited after K_v_1.4 transfection in HEK 293 cells. Cisplatin produced ∼45% reduction of the current density (**E**). However, pre-incubation of vitE did not reverse the cisplatin-mediated reduction of the K_v_1.4-mediated outward current (**F**; n = 12 cells). (*** p < 0.001)

## Discussion

The etiology and targeted DRGNs underlying exaggerated mechanosensitivity following cisplatin chemotherapy remain a puzzle, since targeted degeneration of only large-myelinated DRGNs may not suffice to account for the mechanism underlying CIPN (Starobova and Vetter, 2017; Zajaczkowska et al., 2019). Previous reports show that, within the mechanosensitive subset of TH^+^ DRGNs, two transcriptionally distinct classes, TH1 and TH2, emerged (Finno et al., 2019). The heterogeneity of TH^+^ DRGNs was revealed further by the functional classification into three distinct neurons, namely fast (1-AP), moderately (> 2-AP) adapting, and spontaneously firing (SF) neuronal subtypes. The differential effects of cisplatin in increasing the membrane excitability of the fast and moderately adapting TH^+^ DRNGs, while the response properties of the SF neurons are impervious to the drug, raise the certainty that the TH^+^ DRGNs are not homogeneous. Current analyses show that vitamin E may curb cisplatin’s effects on mechanosensitive TH^+^ DRGN excitability. However, vitamin E may not reverse the loss of mechanical encoding properties of the neurons. A candidate K^+^ channel through which cisplatin may confer neuroexcitatory actions and the apparent protective effects of vitamin E is the two-pore K^+^ channel, K_2P_18.1.

Identifying two distinct TH^+^ subtypes through single-cell transcriptomic profiling is unique to the current dataset (Finno et al., 2019). Previous studies clustered the unmyelinated TH^+^ DRGNs into one population (Usoskin et al., 2015; Li et al., 2016). This earlier research identified the TH^+^ expression in the small neuronal C5-1 and C6-1 subpopulations (Li et al., 2016). These neurons were also *Mrgprd*^+^, which firmly defined the non-peptidergic subpopulation (Finno et al., 2019). Differences in the sampling ages (6-10-weeks (Usoskin et al., 2015; Li et al., 2016) versus 30-weeks (Finno et al., 2019) may invoke potential age-related transcriptional regulation. However, the current functional studies performed on TH-EGFP+ transgenic were from 6-8-week-old mice age, refuting aging as the likely explanation for the seemingly contrasting results from previous reports. In the mouse, a distinct population of TH^+^ DRGNs, constituting 10-14% of TH^+^ DRGNs and located primarily in the lumbar DRG, innervate a portion of colorectal and bladder neurons (Brumovsky et al., 2012). This subset of neurons is positive for CGRP, unlike the non-visceral TH^+^ C-LTMR subpopulation (Brumovsky et al., 2012). While not included in the main transcriptional TH1 and TH2 subsets, these may be the SF neuronal subset that innervates the smooth muscles.

The three functional TH^+^ DRGN subtypes had distinct coding properties and are likely to have distinct roles in sensation. The fast-adapting 1-AP TH^+^ DRGNs had the lowest overall amplitude, input resistance, and the best temporal coding (i.e., high vector strength). These 1-AP DRGNs are time-coding neurons and would likely result in a fast-acting response to touch. Moderately-adapting ≥ 2-APs TH^+^ DRGNs fired ≥ 2-APs, with overall amplitudes, input resistance, and resting membrane potentials similar to the SF TH^+^ DRGNs, but lower current densities than both 1-AP and SF subtypes. Temporal coding of ≥ 2-APs increased at higher stimulation frequencies but remained lower than 1-AP TH+ DRGNs. These ≥ 2-APs neurons would likely result in a moderate response to touch. Lastly, the SF TH+ DRGNs would likely produce a slow reaction to the mechanical sensation as these are rate-coding neurons. Although we have functionally defined these three neuronal subtypes, *in vivo* animal models with specific neuronal sets deleted are required to link the molecular subtypes (i.e., TH1 vs. TH2) to these functional subtypes (1-AP vs. ≥ 2-APs vs. SF).

The platinum derivate chemotherapeutic agents, cisplatin and oxaliplatin, produce painful peripheral neuropathies as dose-limiting side effects. Cisplatin damages all types of myelinated fibers (Boehmerle et al., 2014). In particular, large-diameter myelinated DRGNs (Cavaletti et al., 1992; Krarup-Hansen et al., 1999; McDonald et al., 2005; Krarup-Hansen et al., 2007) are more susceptible. A recent study using *Vglut3*^-/-^ mice, which lack TH^+^ DRGNs and thus C-LTMRs, demonstrated that Vglut3 cells are necessary to fully express mechanical hypersensitivity in oxaliplatin-induced neuropathy (12). We have shown that cisplatin differentially affected TH^+^ DRGNs, with no effect on SF subtypes but a reduction in outward current in both 1-AP and ≥ 2-APs TH^+^ DRGN subtypes. Cisplatin increased spike frequency and significantly reduced vector strength, suggesting stimulus-response temporal coding in fast-adapting 1-AP TH^+^ DRGNs declines.

The putative mechanisms of the neurotoxicity associated with cisplatin chemotherapy include binding to mitochondrial DNA, leading to altered mitochondrial function and release of reactive oxygen species, changes in axon morphology, leading to sensory-motor axon degeneration, and/or chelation of extracellular calcium, resulting in altered calcium homeostasis (Starobova and Vetter, 2017). As a potent antioxidant, the neuroprotective effects of vitamin E against CIPN have been previously documented but were focused primarily on the effects on large-diameter myelinated DRGNs (Cavaletti et al., 1992; Leonetti et al., 2003; Pace et al., 2003). *In vitro* pre-incubation with vitamin E blocked the cisplatin-induced neuron excitability in fast-acting 1-AP TH^+^ DRGNs, without restoring vector strength. With the discovery that vitamin E provides a protective effect against cisplatin-induced neurotoxicity within TH^+^ DRGNs, other agents that target these DRGN subtypes could be investigated to alleviate the neurotoxic effects of CIPN.

We provide evidence that the neuroprotective effects of vitamin E on cisplatin-induced CIPN may be modulated through K_2P_18.1 *(Kcnk18*). Knck18 encodes TWIK-related K^+^ channel (TRESK), a K_2P_ channel with a prominent role in pain pathways. Pharmacologic inhibition of TRESK induces spontaneous pain behavior (Tulleuda et al., 2011). In humans, a dominant-negative frameshift mutation in *KCNK18* segregates in patients with migraines (Lafreniere et al., 2010), and a causal role for this loss of function mutation was recently established (Pettingill et al., 2019). TRESK heteromerizes with two distantly related K_2P_ channels, TREK1 and TREK2 (Royal et al., 2019). Decreased expression of TREK1 was identified after treatment with a related platinum-based chemotherapeutic, oxaliplatin, in murine DRGNs (Descoeur et al., 2011). Moreover, in our previous studies using a vitamin E deficiency mouse model, *Kcnk18* was significantly upregulated in multiple DRGN subpopulations, including peptidergic and TH^+^ DRGNs (Finno et al., 2019). Thus, TRESK, and its interaction with TREK1, may be potential targets across DRGN subpopulations of cisplatin neurotoxicity. The neuroprotective role of vitamin E could be in maintaining the baseline excitability of this channel.

Despite early results that supported a protective role for vitamin E in the presentation of CIPN (Pace et al., 2003; Argyriou et al., 2005; Argyriou et al., 2006; Pace et al., 2010), a large-scale randomized clinical trial in 2011 refuted these results, concluding that vitamin E was ineffective in preventing CIPN (Kottschade et al., 2011). However, the 2011 clinical trial consisted of CIPN induced primarily by taxanes, whereas the earlier clinical trials specifically investigated the use of vitamin E in preventing cisplatin-induced CIPN. Additionally, the trials administered vitamin E at the onset of chemotherapy, rather than initiating supplementation before chemotherapy, similar to the design used in most animal studies (Leonetti et al., 2003) and analogous to pre-incubation with vitamin E in the *in vitro* experiments. A meta-analysis demonstrated that, while vitamin E supplementation did not decrease the overall incidence of CIPN, vitamin E significantly prevented cisplatin-associated neurotoxicity (Huang et al., 2016). Thus, while the current literature suggests vitamin E provides a minimal clinical improvement in CIPN (Hu et al., 2019a), early and targeted vitamin E supplementation to prevent cisplatin-associated peripheral neuropathy requires investigation.

We have identified three functionally distinct TH^+^ DRGNs with differential responses to cisplatin. The addition of vitamin E reduced the cisplatin-mediated increased excitability of 1-AP TH^+^ DRGNs but did not improve the temporal coding properties. The neuroprotective effects of vitamin E on CIPN may be modulated through K_2P_18.1 *(Kcnk18*) across DRGN subpopulations. The study provides a potential therapeutic target in patients receiving cisplatin chemotherapy.

## Author contributions

CJF and ENY were responsible for the conceptualization, funding acquisition, methodology, investigation, formal analysis, resources, and manuscript writing. YC, SP, JL, and CP contributed to the study and analysis of results. All authors have reviewed the final manuscript.

## Abbreviations

AP: action potential
AVED: ataxia with vitamin E deficiency
CIPN: cisplatin-induced peripheral neuropathy
C-LTMRs: C-low threshold mechanoreceptors
DRGN: dorsal root ganglia neurons
IACUC: institutional animal care and use committee
scRNA-seq: single-cell RNA-sequencing
TH: tyrosine hydroxylase
Ttpa: tocopherol transfer protein (alpha)
SF: spontaneously firing
smFISH: single molecule fluorescence in-situ hybridization
UCD: University of California Davis
UCR: University of Reno

## Acknowledgments

The National Institutes of Health (NIH) supported this work to CJF (L40 TR001136). ENY was supported by (NIH: AG051443, DC015135, AG060504, and DC016099). The authors have declared that no conflict of interest exists.

