## Supplemental File for "Cisplatin neurotoxicity targets specific subpopulations and K^+^ channels in tyrosine-hydroxylase positive dorsal root ganglia neurons"

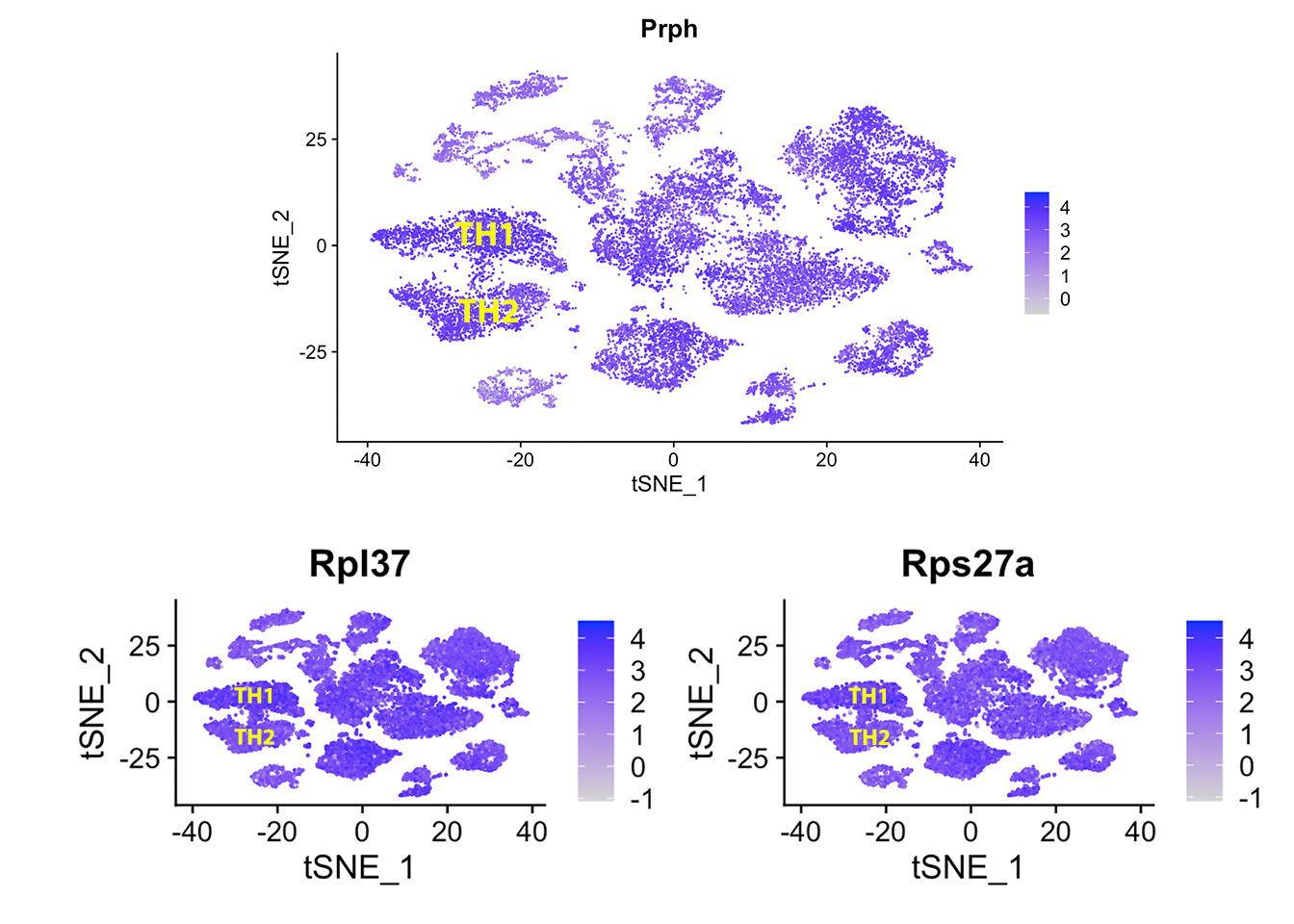

**Figure S1. TH1 specific transcripts.** TSNE plots of the top differentially expressed TH1 transcripts, *Prph, Rpl37,* and *Rps27a*. While significantly upregulated in TH1 DRGNs, these transcripts were highly expressed in all DRGN subpopulations.

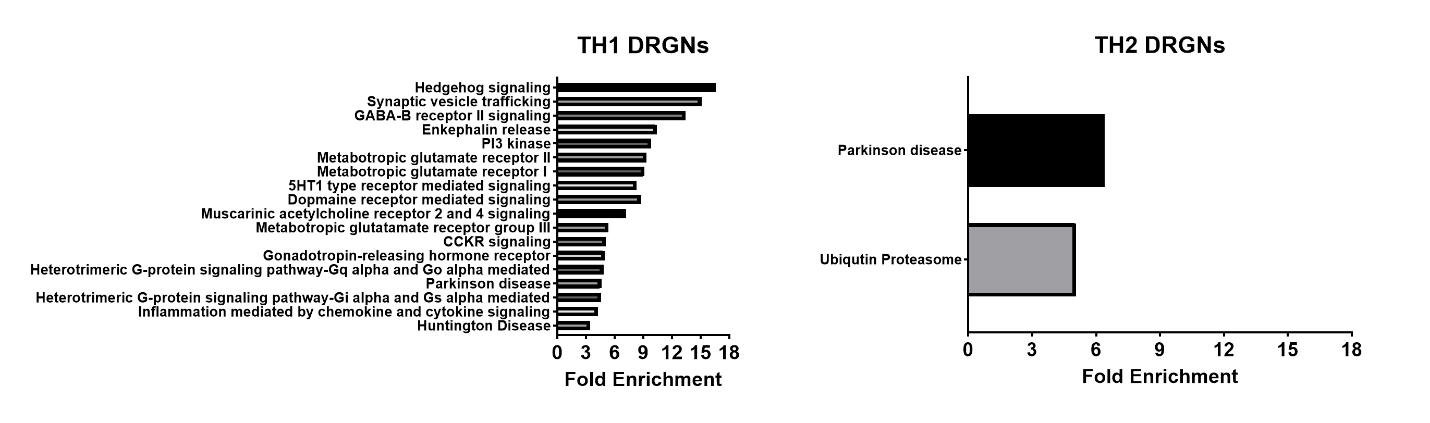

**Figure S2. TH1 and TH2 transcriptomic profiles constitute distinct biologic pathways.** Within TH1 DRGNs (**left**), 18 pathways were statistically enriched, including the three metabotropic glutamate receptor pathways and many G-protein-related receptor signaling pathways. Within TH2 DRGNs (**right**), had Parkinson's disease, and the ubiquitin-proteasome pathways were enriched.

| **Table S1: Genes unique to TH1 and TH2 when compared to all other dorsal root ganglia neuronal subpopulations** | | | | | | |
| --- | --- | --- | --- | --- | --- | --- |
| gene | p_val | avg_logFC | pct.1 | pct.2 | p_val_adj | Cluster |
| Prph | 1.00E-305 | 0.432017 | 1 | 0.986 | 1.14E-301 | TH1 |
| Rpl37 | 4.07E-281 | 0.35792 | 0.998 | 0.996 | 4.66E-277 | TH1 |
| Fau | 5.74E-281 | 0.39051 | 0.996 | 0.99 | 6.57E-277 | TH1 |
| Prune2 | 5.36E-261 | 0.894773 | 0.993 | 0.969 | 6.13E-257 | TH1 |
| Rps27a | 2.39E-259 | 0.373122 | 0.996 | 0.991 | 2.73E-255 | TH1 |
| Rplp0 | 2.42E-212 | 0.478595 | 0.96 | 0.908 | 2.77E-208 | TH1 |
| Klf7 | 8.29E-179 | 0.35463 | 0.986 | 0.963 | 9.49E-175 | TH1 |
| Rpl6 | 2.92E-162 | 0.341402 | 0.982 | 0.968 | 3.34E-158 | TH1 |
| Rpl26 | 8.09E-158 | 0.285129 | 0.995 | 0.988 | 9.25E-154 | TH1 |
| Ncam1 | 7.86E-156 | 0.668321 | 0.832 | 0.705 | 8.99E-152 | TH1 |
| Pfdn5 | 9.64E-148 | 0.385181 | 0.956 | 0.93 | 1.10E-143 | TH1 |
| Rpl35a | 1.05E-134 | 0.321054 | 0.985 | 0.977 | 1.20E-130 | TH1 |
| Rps7 | 1.75E-133 | 0.331036 | 0.972 | 0.949 | 2.00E-129 | TH1 |
| Ptms1 | 2.84E-115 | 0.362365 | 0.953 | 0.933 | 3.25E-111 | TH1 |
| Rpl11 | 2.43E-111 | 0.269275 | 0.99 | 0.975 | 2.77E-107 | TH1 |
| Chd51 | 1.38E-102 | 0.692368 | 0.702 | 0.615 | 1.57E-98 | TH1 |
| Scn9a | 7.30E-100 | 0.421536 | 0.909 | 0.863 | 8.36E-96 | TH1 |
| Rpl7 | 7.16E-90 | 0.331971 | 0.955 | 0.929 | 8.19E-86 | TH1 |
| Eef1g1 | 3.52E-86 | 0.276172 | 0.967 | 0.957 | 4.02E-82 | TH1 |
| Cox7a2l | 1.68E-85 | 0.302427 | 0.98 | 0.972 | 1.93E-81 | TH1 |
| Rps26 | 2.80E-85 | 0.281773 | 0.954 | 0.937 | 3.20E-81 | TH1 |
| Zcrb1 | 6.82E-84 | 0.394996 | 0.84 | 0.789 | 7.81E-80 | TH1 |
| Use1 | 1.02E-78 | 0.38411 | 0.903 | 0.873 | 1.17E-74 | TH1 |
| Serp2 | 1.21E-77 | 0.340231 | 0.931 | 0.887 | 1.39E-73 | TH1 |
| Rpl4 | 1.71E-77 | 0.299348 | 0.964 | 0.935 | 1.95E-73 | TH1 |
| Hnrnpa11 | 2.80E-77 | 0.318551 | 0.893 | 0.871 | 3.21E-73 | TH1 |
| Rbfox1 | 3.03E-76 | 0.399667 | 0.871 | 0.831 | 3.46E-72 | TH1 |
| Ythdf2 | 3.65E-75 | 0.532303 | 0.744 | 0.675 | 4.17E-71 | TH1 |
| Slc9a61 | 1.13E-69 | 0.365619 | 0.835 | 0.788 | 1.29E-65 | TH1 |
| Prrxl1 | 7.62E-65 | 0.547931 | 0.836 | 0.821 | 8.72E-61 | TH1 |
| Eif5b | 2.07E-62 | 0.42484 | 0.794 | 0.785 | 2.37E-58 | TH1 |
| Spop1 | 6.97E-60 | 0.418882 | 0.826 | 0.787 | 7.98E-56 | TH1 |
| Rpl7a | 1.24E-50 | 0.303963 | 0.892 | 0.88 | 1.42E-46 | TH1 |
| Eif3h | 2.50E-50 | 0.331914 | 0.861 | 0.84 | 2.86E-46 | TH1 |
| Rp9 | 5.71E-43 | 0.357921 | 0.826 | 0.809 | 6.53E-39 | TH1 |
| Elavl3 | 4.66E-31 | 0.301481 | 0.755 | 0.747 | 5.33E-27 | TH1 |
| Bach2 | 1.16E-30 | 0.465818 | 0.538 | 0.486 | 1.33E-26 | TH1 |
| Rtf1 | 3.02E-28 | 0.338792 | 0.756 | 0.769 | 3.45E-24 | TH1 |
| Lrrc8d | 7.14E-27 | 0.334482 | 0.66 | 0.647 | 8.17E-23 | TH1 |
| Ppm1l | 1.27E-25 | 0.432289 | 0.588 | 0.555 | 1.45E-21 | TH1 |
| Lats2 | 2.89E-24 | 0.397697 | 0.578 | 0.55 | 3.31E-20 | TH1 |
| Zfhx4 | 1.15E-21 | 0.416961 | 0.68 | 0.633 | 1.31E-17 | TH1 |
| Krit1 | 6.26E-21 | 0.309801 | 0.63 | 0.615 | 7.16E-17 | TH1 |
| Nudc | 1.19E-20 | 0.31148 | 0.791 | 0.789 | 1.36E-16 | TH1 |
| Pik3ca | 1.37E-18 | 0.335337 | 0.641 | 0.631 | 1.56E-14 | TH1 |
| Peli2 | 1.88E-18 | 0.422903 | 0.459 | 0.418 | 2.16E-14 | TH1 |
| Drap1 | 2.36E-18 | 0.263698 | 0.804 | 0.809 | 2.70E-14 | TH1 |
| Map3k1 | 5.42E-18 | 0.430007 | 0.548 | 0.511 | 6.20E-14 | TH1 |
| Creld1 | 1.25E-17 | 0.294119 | 0.766 | 0.753 | 1.43E-13 | TH1 |
| Dgkk | 2.29E-17 | 0.481887 | 0.603 | 0.571 | 2.62E-13 | TH1 |
| Prcp | 4.62E-17 | 0.450608 | 0.5 | 0.468 | 5.29E-13 | TH1 |
| Gm9843 | 4.64E-16 | 0.259909 | 0.803 | 0.802 | 5.31E-12 | TH1 |
| Zbtb7a | 5.09E-16 | 0.282799 | 0.746 | 0.786 | 5.82E-12 | TH1 |
| Zfp703 | 5.05E-15 | 0.460958 | 0.644 | 0.611 | 5.78E-11 | TH1 |
| Tcerg1l | 2.67E-14 | 0.392012 | 0.636 | 0.611 | 3.06E-10 | TH1 |
| Gemin7 | 1.72E-13 | 0.272603 | 0.774 | 0.775 | 1.97E-09 | TH1 |
| Slc25a1 | 1.90E-13 | 0.288763 | 0.721 | 0.749 | 2.17E-09 | TH1 |
| Csnk1g3 | 2.01E-13 | 0.337779 | 0.594 | 0.608 | 2.30E-09 | TH1 |
| Rab11fip2 | 5.79E-13 | 0.340158 | 0.685 | 0.72 | 6.63E-09 | TH1 |
| Lrrc75b | 7.11E-13 | 0.39315 | 0.669 | 0.655 | 8.14E-09 | TH1 |
| Acbd5 | 9.32E-13 | 0.261746 | 0.673 | 0.694 | 1.07E-08 | TH1 |
| Tln2 | 1.86E-11 | 0.278681 | 0.806 | 0.839 | 2.13E-07 | TH1 |
| Arglu1 | 4.07E-11 | 0.282116 | 0.66 | 0.673 | 4.66E-07 | TH1 |
| Suds3 | 5.31E-11 | 0.271127 | 0.758 | 0.758 | 6.08E-07 | TH1 |
| 2510009E07Rik | 1.19E-10 | 0.290971 | 0.615 | 0.621 | 1.36E-06 | TH1 |
| Zmiz1 | 2.39E-10 | 0.340676 | 0.543 | 0.536 | 2.74E-06 | TH1 |
| Med13l | 2.64E-10 | 0.371156 | 0.513 | 0.499 | 3.02E-06 | TH1 |
| AI504432 | 3.42E-10 | 0.373487 | 0.583 | 0.572 | 3.91E-06 | TH1 |
| Nxph3 | 6.37E-10 | 0.448102 | 0.553 | 0.564 | 7.29E-06 | TH1 |
| Deaf1 | 5.30E-09 | 0.358735 | 0.742 | 0.77 | 6.06E-05 | TH1 |
| Coq2 | 8.61E-09 | 0.291979 | 0.734 | 0.739 | 9.85E-05 | TH1 |
| Kcna4 | 1.27E-08 | 0.344269 | 0.637 | 0.63 | 0.000146 | TH1 |
| St6galnac5 | 2.33E-08 | 0.379305 | 0.617 | 0.599 | 0.000267 | TH1 |
| Fdx1l | 3.14E-08 | 0.27637 | 0.742 | 0.751 | 0.000359 | TH1 |
| Cul4a | 5.61E-08 | 0.307175 | 0.546 | 0.542 | 0.000642 | TH1 |
| Uhrf1bp1l | 3.70E-07 | 0.315491 | 0.567 | 0.576 | 0.004234 | TH1 |
| Marf1 | 4.36E-07 | 0.308372 | 0.798 | 0.788 | 0.004985 | TH1 |
| Foxp1 | 4.86E-07 | 0.351098 | 0.54 | 0.53 | 0.005561 | TH1 |
| Twf1 | 9.47E-07 | 0.298991 | 0.534 | 0.544 | 0.010837 | TH1 |
| Maz | 1.25E-06 | 0.312685 | 0.645 | 0.682 | 0.014301 | TH1 |
| Nhs | 6.75E-06 | 0.342398 | 0.492 | 0.487 | 0.077218 | TH1 |
| Fam135a | 8.67E-06 | 0.31793 | 0.466 | 0.471 | 0.099203 | TH1 |
| Rrbp1 | 9.68E-06 | 0.328744 | 0.512 | 0.519 | 0.110723 | TH1 |
| Dpysl5 | 1.12E-05 | 0.31073 | 0.524 | 0.519 | 0.127808 | TH1 |
| Lrfn5 | 1.16E-05 | 0.366255 | 0.663 | 0.662 | 0.132568 | TH1 |
| Rnf141 | 1.36E-05 | 0.274328 | 0.533 | 0.543 | 0.155519 | TH1 |
| Arhgap31 | 1.38E-05 | 0.365106 | 0.425 | 0.418 | 0.158228 | TH1 |
| Srrm1 | 2.11E-05 | 0.254776 | 0.721 | 0.729 | 0.241407 | TH1 |
| Clock | 2.95E-05 | 0.306726 | 0.499 | 0.514 | 0.33781 | TH1 |
| Adamts9 | 4.44E-05 | 0.348983 | 0.585 | 0.575 | 0.508373 | TH1 |
| Fst | 6.67E-05 | 0.358749 | 0.583 | 0.574 | 0.76264 | TH1 |
| Slc12a7 | 0.000158 | 0.263691 | 0.371 | 0.403 | 1 | TH1 |
| Gclc | 0.000197 | 0.313882 | 0.498 | 0.503 | 1 | TH1 |
| Tacc1 | 0.000722 | 0.381927 | 0.575 | 0.619 | 1 | TH1 |
| AW551984 | 0.001047 | 0.351744 | 0.576 | 0.572 | 1 | TH1 |
| Myadm | 0.001139 | 0.282384 | 0.479 | 0.494 | 1 | TH1 |
| Celf3 | 0.001831 | 0.356467 | 0.599 | 0.589 | 1 | TH1 |
| Mrpl9 | 0.002068 | 0.364188 | 0.57 | 0.609 | 1 | TH1 |
| Tmem2 | 0.002427 | 0.267663 | 0.513 | 0.513 | 1 | TH1 |
| Tmf1 | 0.002893 | 0.264799 | 0.702 | 0.684 | 1 | TH1 |
| Dusp6 | 0.003195 | 0.290423 | 0.44 | 0.457 | 1 | TH1 |
| Mesdc1 | 0.004767 | 0.281967 | 0.765 | 0.809 | 1 | TH1 |
| Pla2g71 | 0 | 0.822942 | 0.919 | 0.673 | 0 | TH2 |
| Tsen34 | 0 | 0.678656 | 0.96 | 0.785 | 0 | TH2 |
| Uqcrc11 | 1.12E-284 | 0.660663 | 0.946 | 0.738 | 1.29E-280 | TH2 |
| Pmvk1 | 2.75E-281 | 0.647425 | 0.959 | 0.781 | 3.15E-277 | TH2 |
| Ccdc107 | 6.41E-275 | 0.672978 | 0.902 | 0.695 | 7.33E-271 | TH2 |
| Tceal81 | 1.04E-270 | 0.636131 | 0.943 | 0.745 | 1.19E-266 | TH2 |
| Pdcd101 | 6.28E-248 | 0.620968 | 0.939 | 0.757 | 7.19E-244 | TH2 |
| Akr1b3 | 1.40E-231 | 0.647818 | 0.826 | 0.615 | 1.61E-227 | TH2 |
| Ccdc109b | 3.88E-221 | 0.581838 | 0.871 | 0.679 | 4.44E-217 | TH2 |
| Hsp90ab1 | 1.04E-219 | 0.322049 | 1 | 0.995 | 1.20E-215 | TH2 |
| Higd2a1 | 2.27E-212 | 0.529271 | 0.965 | 0.819 | 2.60E-208 | TH2 |
| Med21 | 5.85E-212 | 0.54358 | 0.905 | 0.711 | 6.69E-208 | TH2 |
| Ceacam103 | 5.24E-203 | 1.272883 | 0.833 | 0.725 | 5.99E-199 | TH2 |
| Bad1 | 6.82E-203 | 0.532035 | 0.877 | 0.689 | 7.80E-199 | TH2 |
| Snrpd31 | 2.66E-189 | 0.533457 | 0.884 | 0.704 | 3.05E-185 | TH2 |
| Hcfc1r1 | 2.95E-189 | 0.462486 | 0.978 | 0.873 | 3.37E-185 | TH2 |
| Psma12 | 2.43E-188 | 0.503568 | 0.922 | 0.812 | 2.77E-184 | TH2 |
| Ndufv11 | 1.96E-187 | 0.521311 | 0.903 | 0.722 | 2.24E-183 | TH2 |
| Cd631 | 7.72E-187 | 0.520567 | 0.894 | 0.697 | 8.83E-183 | TH2 |
| Eif3g | 8.57E-187 | 0.532185 | 0.868 | 0.685 | 9.80E-183 | TH2 |
| Aig11 | 3.43E-186 | 0.535101 | 0.858 | 0.679 | 3.93E-182 | TH2 |
| Dnajc15 | 1.49E-184 | 0.540769 | 0.857 | 0.665 | 1.71E-180 | TH2 |
| Spag7 | 7.64E-182 | 0.498019 | 0.903 | 0.724 | 8.74E-178 | TH2 |
| Mllt11 | 2.17E-180 | 0.390493 | 0.992 | 0.948 | 2.48E-176 | TH2 |
| Timm17a | 3.85E-179 | 0.490779 | 0.942 | 0.782 | 4.41E-175 | TH2 |
| Apoa1bp | 1.76E-178 | 0.505149 | 0.88 | 0.698 | 2.01E-174 | TH2 |
| Sdhd1 | 2.99E-177 | 0.528534 | 0.876 | 0.694 | 3.42E-173 | TH2 |
| Avpi1 | 2.99E-174 | 0.516184 | 0.811 | 0.635 | 3.43E-170 | TH2 |
| Ndufa81 | 1.18E-172 | 0.435433 | 0.968 | 0.836 | 1.35E-168 | TH2 |
| Coa6 | 2.70E-171 | 0.501395 | 0.838 | 0.659 | 3.09E-167 | TH2 |
| Gadd45gip1 | 2.29E-169 | 0.487436 | 0.894 | 0.723 | 2.62E-165 | TH2 |
| Chmp51 | 3.43E-169 | 0.448694 | 0.968 | 0.834 | 3.92E-165 | TH2 |
| 1110001J03Rik | 4.98E-167 | 0.474427 | 0.85 | 0.665 | 5.69E-163 | TH2 |
| Fam173a | 4.55E-166 | 0.476374 | 0.899 | 0.735 | 5.20E-162 | TH2 |
| Mrpl28 | 6.27E-163 | 0.465884 | 0.893 | 0.717 | 7.17E-159 | TH2 |
| Hexb | 1.58E-162 | 0.493125 | 0.868 | 0.768 | 1.81E-158 | TH2 |
| Mcts11 | 5.02E-159 | 0.44022 | 0.885 | 0.695 | 5.74E-155 | TH2 |
| Med31 | 1.63E-154 | 0.453685 | 0.845 | 0.672 | 1.86E-150 | TH2 |
| 2-Mar | 1.13E-149 | 0.475578 | 0.808 | 0.639 | 1.29E-145 | TH2 |
| Rheb1 | 3.14E-147 | 0.374684 | 0.989 | 0.904 | 3.59E-143 | TH2 |
| Dhrs71 | 3.91E-147 | 0.515795 | 0.804 | 0.624 | 4.47E-143 | TH2 |
| Psmc2 | 2.13E-145 | 0.462279 | 0.879 | 0.794 | 2.44E-141 | TH2 |
| Ethe1 | 5.07E-143 | 0.471073 | 0.804 | 0.635 | 5.80E-139 | TH2 |
| Chchd1 | 5.25E-143 | 0.431218 | 0.9 | 0.736 | 6.00E-139 | TH2 |
| Mrpl211 | 4.22E-140 | 0.429845 | 0.861 | 0.692 | 4.83E-136 | TH2 |
| Ccl1 | 2.28E-136 | 0.539313 | 0.737 | 0.568 | 2.61E-132 | TH2 |
| Tex2641 | 6.72E-135 | 0.448812 | 0.847 | 0.68 | 7.69E-131 | TH2 |
| Emc21 | 8.06E-134 | 0.410535 | 0.844 | 0.676 | 9.22E-130 | TH2 |
| Bcas2 | 1.59E-133 | 0.392146 | 0.917 | 0.76 | 1.82E-129 | TH2 |
| Sec62 | 2.21E-132 | 0.376183 | 0.964 | 0.836 | 2.52E-128 | TH2 |
| Ccdc167 | 3.66E-132 | 0.47277 | 0.791 | 0.633 | 4.18E-128 | TH2 |
| Psmb4 | 9.33E-132 | 0.395216 | 0.928 | 0.785 | 1.07E-127 | TH2 |
| Tmem60 | 1.27E-131 | 0.413895 | 0.894 | 0.735 | 1.45E-127 | TH2 |
| Tceb1 | 1.59E-131 | 0.379556 | 0.981 | 0.876 | 1.82E-127 | TH2 |
| Ndufa101 | 5.42E-131 | 0.410077 | 0.922 | 0.78 | 6.20E-127 | TH2 |
| Mrpl46 | 1.82E-129 | 0.439035 | 0.806 | 0.646 | 2.08E-125 | TH2 |
| Pja1 | 3.11E-128 | 0.408725 | 0.856 | 0.693 | 3.56E-124 | TH2 |
| Uqcc2 | 3.82E-124 | 0.352499 | 0.97 | 0.832 | 4.37E-120 | TH2 |
| Mrpl12 | 3.84E-124 | 0.386947 | 0.845 | 0.7 | 4.39E-120 | TH2 |
| Timm101 | 7.87E-124 | 0.400831 | 0.853 | 0.701 | 9.00E-120 | TH2 |
| Dohh | 1.77E-123 | 0.413898 | 0.87 | 0.72 | 2.03E-119 | TH2 |
| Snapc5 | 7.86E-123 | 0.395547 | 0.864 | 0.716 | 8.99E-119 | TH2 |
| Ppp1r35 | 4.58E-122 | 0.426272 | 0.833 | 0.674 | 5.24E-118 | TH2 |
| Polr2k1 | 1.88E-119 | 0.380521 | 0.968 | 0.838 | 2.15E-115 | TH2 |
| Fars2 | 2.80E-117 | 0.390104 | 0.751 | 0.602 | 3.21E-113 | TH2 |
| Mrpl14 | 8.54E-117 | 0.360141 | 0.904 | 0.752 | 9.77E-113 | TH2 |
| 0610011F06Rik | 1.21E-116 | 0.39621 | 0.796 | 0.637 | 1.38E-112 | TH2 |
| Psmd6 | 9.93E-116 | 0.39034 | 0.843 | 0.684 | 1.14E-111 | TH2 |
| Tsen15 | 1.29E-114 | 0.400875 | 0.796 | 0.643 | 1.48E-110 | TH2 |
| Rbm8a | 1.58E-113 | 0.346309 | 0.874 | 0.719 | 1.80E-109 | TH2 |
| Aamp | 7.70E-113 | 0.387455 | 0.819 | 0.668 | 8.81E-109 | TH2 |
| Pfdn1 | 8.09E-113 | 0.349132 | 0.944 | 0.806 | 9.26E-109 | TH2 |
| Mrps26 | 6.15E-112 | 0.37046 | 0.858 | 0.724 | 7.03E-108 | TH2 |
| Ostc | 3.88E-111 | 0.370464 | 0.872 | 0.725 | 4.44E-107 | TH2 |
| Suclg1 | 9.61E-111 | 0.374389 | 0.811 | 0.659 | 1.10E-106 | TH2 |
| Snrpd1 | 1.16E-110 | 0.379745 | 0.842 | 0.694 | 1.32E-106 | TH2 |
| Polr2e1 | 4.47E-110 | 0.402443 | 0.807 | 0.657 | 5.11E-106 | TH2 |
| Rpp211 | 2.90E-109 | 0.361639 | 0.855 | 0.704 | 3.32E-105 | TH2 |
| Txnl1 | 8.56E-109 | 0.355249 | 0.901 | 0.759 | 9.79E-105 | TH2 |
| Mzt2 | 1.58E-108 | 0.415 | 0.767 | 0.622 | 1.80E-104 | TH2 |
| Fam132a | 2.30E-108 | 0.372928 | 0.773 | 0.626 | 2.63E-104 | TH2 |
| Tm2d2 | 3.35E-108 | 0.362386 | 0.843 | 0.694 | 3.83E-104 | TH2 |
| Naa201 | 3.43E-107 | 0.422746 | 0.804 | 0.651 | 3.93E-103 | TH2 |
| Ict1 | 6.34E-107 | 0.341456 | 0.844 | 0.692 | 7.25E-103 | TH2 |
| 1810037I17Rik | 6.95E-107 | 0.330785 | 0.951 | 0.815 | 7.95E-103 | TH2 |
| Hmgb31 | 8.08E-107 | 0.400037 | 0.756 | 0.613 | 9.24E-103 | TH2 |
| Ten1 | 8.41E-107 | 0.342054 | 0.827 | 0.682 | 9.62E-103 | TH2 |
| Acot9 | 1.37E-106 | 0.367484 | 0.785 | 0.634 | 1.56E-102 | TH2 |
| Snx31 | 2.75E-105 | 0.353129 | 0.945 | 0.821 | 3.15E-101 | TH2 |
| 2610001J05Rik | 3.63E-105 | 0.348978 | 0.926 | 0.792 | 4.16E-101 | TH2 |
| Eny2 | 6.32E-105 | 0.358398 | 0.861 | 0.717 | 7.23E-101 | TH2 |
| Pcmt1 | 9.82E-105 | 0.337873 | 0.961 | 0.846 | 1.12E-100 | TH2 |
| Unc50 | 5.44E-104 | 0.389778 | 0.825 | 0.676 | 6.22E-100 | TH2 |
| Cirbp | 7.40E-104 | 0.328442 | 0.921 | 0.782 | 8.46E-100 | TH2 |
| Hes6 | 1.15E-103 | 0.404341 | 0.783 | 0.629 | 1.32E-99 | TH2 |
| Plpp1 | 2.03E-103 | 0.369936 | 0.754 | 0.611 | 2.32E-99 | TH2 |
| Phf5a1 | 3.63E-103 | 0.325686 | 0.834 | 0.691 | 4.15E-99 | TH2 |
| Eif6 | 9.02E-103 | 0.356822 | 0.86 | 0.711 | 1.03E-98 | TH2 |
| Apoo | 1.62E-102 | 0.381067 | 0.835 | 0.774 | 1.85E-98 | TH2 |
| Mrpl43 | 1.74E-102 | 0.35836 | 0.828 | 0.693 | 1.99E-98 | TH2 |
| 2010111I01Rik | 9.12E-102 | 0.307846 | 0.834 | 0.687 | 1.04E-97 | TH2 |
| Gnpda2 | 1.32E-101 | 0.321774 | 0.853 | 0.702 | 1.51E-97 | TH2 |
| Slc35g2 | 2.13E-101 | 0.367231 | 0.734 | 0.586 | 2.44E-97 | TH2 |
| Ssna1 | 4.47E-100 | 0.318648 | 0.867 | 0.719 | 5.12E-96 | TH2 |
| Snrpc | 7.52E-100 | 0.37206 | 0.788 | 0.644 | 8.61E-96 | TH2 |
| Sugt1 | 1.19E-99 | 0.318137 | 0.841 | 0.701 | 1.36E-95 | TH2 |
| Lsm5 | 3.79E-99 | 0.363355 | 0.784 | 0.574 | 4.34E-95 | TH2 |
| Mtch2 | 4.79E-99 | 0.332364 | 0.892 | 0.747 | 5.48E-95 | TH2 |
| Ppa1 | 1.92E-98 | 0.340284 | 0.811 | 0.66 | 2.20E-94 | TH2 |
| Plpp2 | 2.50E-98 | 0.313822 | 0.733 | 0.599 | 2.86E-94 | TH2 |
| Mrps21 | 4.14E-98 | 0.326154 | 0.901 | 0.713 | 4.74E-94 | TH2 |
| Yipf4 | 3.33E-97 | 0.329099 | 0.93 | 0.796 | 3.81E-93 | TH2 |
| Med29 | 3.69E-97 | 0.350933 | 0.793 | 0.651 | 4.22E-93 | TH2 |
| Cacybp | 5.03E-97 | 0.334627 | 0.908 | 0.773 | 5.75E-93 | TH2 |
| Clic1 | 1.04E-96 | 0.323636 | 0.915 | 0.774 | 1.19E-92 | TH2 |
| Prmt1 | 4.92E-96 | 0.365509 | 0.802 | 0.652 | 5.63E-92 | TH2 |
| Isoc11 | 5.92E-96 | 0.419419 | 0.756 | 0.615 | 6.78E-92 | TH2 |
| Pin41 | 1.01E-94 | 0.314374 | 0.847 | 0.648 | 1.15E-90 | TH2 |
| Alkbh7 | 9.51E-94 | 0.383281 | 0.755 | 0.62 | 1.09E-89 | TH2 |
| Mrps18a | 1.69E-92 | 0.324427 | 0.884 | 0.76 | 1.93E-88 | TH2 |
| Timm22 | 2.94E-92 | 0.300203 | 0.803 | 0.667 | 3.36E-88 | TH2 |
| Atg1011 | 8.45E-92 | 0.357845 | 0.806 | 0.662 | 9.67E-88 | TH2 |
| Carkd | 8.88E-92 | 0.354089 | 0.758 | 0.62 | 1.02E-87 | TH2 |
| Tm2d3 | 2.36E-91 | 0.373077 | 0.771 | 0.64 | 2.70E-87 | TH2 |
| 1810058I24Rik | 5.37E-91 | 0.306389 | 0.953 | 0.861 | 6.14E-87 | TH2 |
| Flot11 | 6.58E-91 | 0.306239 | 0.919 | 0.782 | 7.53E-87 | TH2 |
| Tmem1471 | 7.40E-91 | 0.308395 | 0.915 | 0.778 | 8.47E-87 | TH2 |
| Sssca1 | 7.68E-91 | 0.355255 | 0.792 | 0.653 | 8.79E-87 | TH2 |
| Ech1 | 7.35E-90 | 0.327563 | 0.773 | 0.638 | 8.41E-86 | TH2 |
| Psmd4 | 1.02E-89 | 0.30828 | 0.886 | 0.755 | 1.16E-85 | TH2 |
| Serpinf1 | 1.41E-89 | 0.420308 | 0.749 | 0.592 | 1.61E-85 | TH2 |
| Lhfpl5 | 1.38E-88 | 0.428293 | 0.753 | 0.606 | 1.58E-84 | TH2 |
| Pts | 2.06E-88 | 0.336548 | 0.807 | 0.675 | 2.36E-84 | TH2 |
| Sec61b | 2.08E-88 | 0.30524 | 0.911 | 0.773 | 2.38E-84 | TH2 |
| Sdf2 | 6.38E-88 | 0.319877 | 0.827 | 0.684 | 7.30E-84 | TH2 |
| Mrpl551 | 6.81E-88 | 0.287244 | 0.871 | 0.731 | 7.79E-84 | TH2 |
| Smarcb1 | 7.20E-88 | 0.308239 | 0.748 | 0.609 | 8.24E-84 | TH2 |
| Timm10b | 1.25E-87 | 0.307394 | 0.861 | 0.724 | 1.43E-83 | TH2 |
| Adrm1 | 2.22E-86 | 0.323347 | 0.801 | 0.659 | 2.53E-82 | TH2 |
| D10Jhu81e | 3.94E-86 | 0.34422 | 0.767 | 0.634 | 4.51E-82 | TH2 |
| Jagn1 | 2.89E-85 | 0.355677 | 0.8 | 0.66 | 3.31E-81 | TH2 |
| Hspbp1 | 2.07E-84 | 0.348751 | 0.781 | 0.648 | 2.37E-80 | TH2 |
| Prmt2 | 3.96E-84 | 0.279074 | 0.836 | 0.702 | 4.53E-80 | TH2 |
| Eci1 | 2.28E-83 | 0.392211 | 0.759 | 0.613 | 2.61E-79 | TH2 |
| Ccdc124 | 2.45E-83 | 0.303276 | 0.921 | 0.799 | 2.80E-79 | TH2 |
| Tmem55b | 1.52E-82 | 0.284856 | 0.903 | 0.773 | 1.74E-78 | TH2 |
| Sumo2 | 2.69E-82 | 0.267193 | 0.988 | 0.923 | 3.07E-78 | TH2 |
| Acyp21 | 6.04E-81 | 0.292745 | 0.844 | 0.711 | 6.91E-77 | TH2 |
| Cct71 | 1.29E-80 | 0.288977 | 0.859 | 0.726 | 1.47E-76 | TH2 |
| Fam96b | 2.81E-80 | 0.274508 | 0.921 | 0.793 | 3.21E-76 | TH2 |
| Gm6710 | 6.23E-80 | 0.312142 | 0.743 | 0.614 | 7.13E-76 | TH2 |
| Snrnp27 | 1.46E-79 | 0.276389 | 0.881 | 0.704 | 1.67E-75 | TH2 |
| Uqcrc2 | 6.92E-79 | 0.292201 | 0.801 | 0.668 | 7.91E-75 | TH2 |
| Slc25a11 | 7.48E-79 | 0.288884 | 0.798 | 0.68 | 8.55E-75 | TH2 |
| Psme1 | 1.27E-78 | 0.306202 | 0.74 | 0.615 | 1.46E-74 | TH2 |
| Mrps11 | 3.38E-78 | 0.363207 | 0.767 | 0.637 | 3.86E-74 | TH2 |
| Yipf1 | 4.54E-78 | 0.319357 | 0.755 | 0.627 | 5.20E-74 | TH2 |
| Sec13 | 1.77E-77 | 0.306449 | 0.762 | 0.636 | 2.02E-73 | TH2 |
| Frg1 | 3.70E-77 | 0.299596 | 0.781 | 0.649 | 4.24E-73 | TH2 |
| Nxf1 | 4.34E-77 | 0.338014 | 0.801 | 0.683 | 4.97E-73 | TH2 |
| Pdcd5 | 5.79E-77 | 0.268492 | 0.945 | 0.796 | 6.63E-73 | TH2 |
| Etfa | 8.09E-77 | 0.313241 | 0.824 | 0.699 | 9.26E-73 | TH2 |
| Psmd142 | 8.54E-77 | 0.277305 | 0.847 | 0.722 | 9.77E-73 | TH2 |
| Rraga | 1.51E-76 | 0.281151 | 0.884 | 0.763 | 1.73E-72 | TH2 |
| Sra1 | 3.29E-76 | 0.280286 | 0.83 | 0.702 | 3.76E-72 | TH2 |
| Mrps36 | 3.53E-76 | 0.263949 | 0.892 | 0.752 | 4.04E-72 | TH2 |
| Slc35b1 | 4.62E-76 | 0.311844 | 0.766 | 0.642 | 5.28E-72 | TH2 |
| Cdc5l | 4.76E-76 | 0.311974 | 0.745 | 0.621 | 5.45E-72 | TH2 |
| Ndufaf3 | 1.19E-75 | 0.26442 | 0.819 | 0.689 | 1.37E-71 | TH2 |
| Rpl36al | 1.59E-75 | 0.261367 | 0.952 | 0.85 | 1.82E-71 | TH2 |
| Mrps16 | 2.45E-75 | 0.274945 | 0.806 | 0.685 | 2.80E-71 | TH2 |
| Znhit3 | 5.46E-75 | 0.30969 | 0.768 | 0.639 | 6.24E-71 | TH2 |
| Tsg101 | 1.29E-74 | 0.26271 | 0.884 | 0.758 | 1.48E-70 | TH2 |
| Magoh | 1.60E-74 | 0.32344 | 0.82 | 0.771 | 1.84E-70 | TH2 |
| Idh3g | 1.90E-74 | 0.25541 | 0.754 | 0.632 | 2.17E-70 | TH2 |
| 2310036O22Rik | 2.19E-74 | 0.275193 | 0.929 | 0.819 | 2.51E-70 | TH2 |
| Psmc6 | 5.60E-74 | 0.281085 | 0.824 | 0.699 | 6.40E-70 | TH2 |
| Lsm1 | 1.10E-73 | 0.280061 | 0.83 | 0.7 | 1.26E-69 | TH2 |
| Tbc1d19 | 5.94E-73 | 0.291362 | 0.747 | 0.62 | 6.79E-69 | TH2 |
| Psme2 | 1.17E-72 | 0.25836 | 0.836 | 0.704 | 1.34E-68 | TH2 |
| Mea1 | 1.46E-72 | 0.25828 | 0.88 | 0.762 | 1.67E-68 | TH2 |
| Coro1a | 3.81E-72 | 0.29604 | 0.763 | 0.652 | 4.36E-68 | TH2 |
| Fkbp2 | 4.31E-72 | 0.261424 | 0.964 | 0.864 | 4.94E-68 | TH2 |
| Txndc17 | 6.10E-72 | 0.268749 | 0.932 | 0.836 | 6.98E-68 | TH2 |
| Med30 | 1.42E-71 | 0.280627 | 0.813 | 0.684 | 1.62E-67 | TH2 |
| Atg3 | 2.63E-71 | 0.30999 | 0.845 | 0.716 | 3.01E-67 | TH2 |
| Psmd12 | 6.80E-71 | 0.293043 | 0.806 | 0.698 | 7.78E-67 | TH2 |
| Bud31 | 1.12E-70 | 0.294413 | 0.774 | 0.651 | 1.28E-66 | TH2 |
| Rbm42 | 2.17E-70 | 0.30489 | 0.737 | 0.616 | 2.48E-66 | TH2 |
| Isca2 | 2.59E-70 | 0.282872 | 0.831 | 0.715 | 2.97E-66 | TH2 |
| Pafah1b3 | 2.67E-70 | 0.28348 | 0.835 | 0.701 | 3.06E-66 | TH2 |
| Tmem14a | 2.73E-70 | 0.274115 | 0.836 | 0.712 | 3.12E-66 | TH2 |
| Mrpl4 | 3.42E-70 | 0.273963 | 0.756 | 0.628 | 3.92E-66 | TH2 |
| Popdc3 | 3.89E-70 | 0.250303 | 0.744 | 0.628 | 4.45E-66 | TH2 |
| Cbr1 | 4.10E-70 | 0.270443 | 0.784 | 0.659 | 4.69E-66 | TH2 |
| Psmd8 | 6.14E-70 | 0.277627 | 0.816 | 0.694 | 7.02E-66 | TH2 |
| Nipsnap3b | 9.09E-70 | 0.264272 | 0.848 | 0.716 | 1.04E-65 | TH2 |
| Sar1b | 2.56E-69 | 0.296559 | 0.794 | 0.662 | 2.93E-65 | TH2 |
| Vbp1 | 3.07E-69 | 0.25826 | 0.952 | 0.845 | 3.52E-65 | TH2 |
| Ddost | 6.34E-69 | 0.297437 | 0.746 | 0.626 | 7.26E-65 | TH2 |
| Gpx2 | 2.03E-68 | 0.334909 | 0.687 | 0.567 | 2.32E-64 | TH2 |
| Bola1 | 2.47E-68 | 0.279016 | 0.787 | 0.668 | 2.83E-64 | TH2 |
| Tmem53 | 2.71E-68 | 0.287624 | 0.706 | 0.586 | 3.10E-64 | TH2 |
| Uqcc3 | 6.05E-68 | 0.250813 | 0.9 | 0.785 | 6.92E-64 | TH2 |
| Aimp2 | 7.65E-68 | 0.269532 | 0.757 | 0.633 | 8.76E-64 | TH2 |
| 2210013O21Rik | 1.29E-67 | 0.259734 | 0.954 | 0.853 | 1.48E-63 | TH2 |
| Tsn | 1.51E-67 | 0.261959 | 0.887 | 0.773 | 1.73E-63 | TH2 |
| Sf3b61 | 4.22E-67 | 0.283008 | 0.823 | 0.696 | 4.82E-63 | TH2 |
| Txndc15 | 5.26E-67 | 0.276838 | 0.775 | 0.657 | 6.02E-63 | TH2 |
| Dapk3 | 2.48E-66 | 0.256211 | 0.742 | 0.631 | 2.84E-62 | TH2 |
| 2810428I15Rik | 4.04E-66 | 0.250959 | 0.963 | 0.876 | 4.63E-62 | TH2 |
| Eri3 | 3.76E-65 | 0.252361 | 0.808 | 0.69 | 4.31E-61 | TH2 |
| Lsm7 | 3.91E-65 | 0.262171 | 0.688 | 0.511 | 4.47E-61 | TH2 |
| Pepd | 1.21E-64 | 0.296512 | 0.71 | 0.597 | 1.38E-60 | TH2 |
| Hacd1 | 5.10E-64 | 0.28803 | 0.766 | 0.65 | 5.84E-60 | TH2 |
| Cyb561d2 | 1.11E-63 | 0.256299 | 0.759 | 0.642 | 1.27E-59 | TH2 |
| Nsfl1c | 1.72E-63 | 0.25477 | 0.81 | 0.692 | 1.97E-59 | TH2 |
| Mvb12a1 | 2.30E-63 | 0.260401 | 0.78 | 0.67 | 2.63E-59 | TH2 |
| Lyrm4 | 7.18E-63 | 0.265531 | 0.777 | 0.667 | 8.22E-59 | TH2 |
| Ola1 | 8.08E-63 | 0.252246 | 0.837 | 0.725 | 9.24E-59 | TH2 |
| Slc51a | 2.38E-62 | 0.34596 | 0.674 | 0.557 | 2.72E-58 | TH2 |
| Mrpl15 | 4.35E-61 | 0.299855 | 0.737 | 0.628 | 4.98E-57 | TH2 |
| Tomm40 | 1.20E-60 | 0.272529 | 0.743 | 0.632 | 1.38E-56 | TH2 |
| Tssc4 | 2.81E-60 | 0.257845 | 0.709 | 0.596 | 3.22E-56 | TH2 |
| Med19 | 6.82E-60 | 0.260578 | 0.766 | 0.655 | 7.80E-56 | TH2 |
| Yif1a | 1.95E-59 | 0.28756 | 0.773 | 0.654 | 2.23E-55 | TH2 |
| Ercc1 | 2.14E-59 | 0.274573 | 0.712 | 0.598 | 2.44E-55 | TH2 |
| Tmco1 | 3.07E-59 | 0.263806 | 0.823 | 0.71 | 3.52E-55 | TH2 |
| Blvra | 5.70E-58 | 0.286699 | 0.743 | 0.617 | 6.52E-54 | TH2 |
| Psmg3 | 4.92E-57 | 0.264669 | 0.732 | 0.624 | 5.62E-53 | TH2 |
| Mrpl22 | 8.96E-57 | 0.310539 | 0.825 | 0.767 | 1.02E-52 | TH2 |
| Sac3d1 | 9.05E-56 | 0.254731 | 0.753 | 0.645 | 1.04E-51 | TH2 |
| Ssr2 | 9.33E-56 | 0.269323 | 0.763 | 0.651 | 1.07E-51 | TH2 |
| Commd9 | 1.25E-55 | 0.276262 | 0.739 | 0.625 | 1.43E-51 | TH2 |
| Tbrg1 | 1.39E-55 | 0.270968 | 0.75 | 0.632 | 1.59E-51 | TH2 |
| Clpp | 2.28E-55 | 0.256331 | 0.806 | 0.69 | 2.61E-51 | TH2 |
| Ubl7 | 2.62E-55 | 0.254613 | 0.731 | 0.627 | 3.00E-51 | TH2 |
| Lsm3 | 1.36E-54 | 0.259927 | 0.728 | 0.622 | 1.56E-50 | TH2 |
| Creb3 | 1.88E-54 | 0.256375 | 0.716 | 0.61 | 2.15E-50 | TH2 |
| Emd | 1.55E-53 | 0.284974 | 0.744 | 0.631 | 1.77E-49 | TH2 |
| Gpr137b | 4.50E-53 | 0.258567 | 0.699 | 0.594 | 5.15E-49 | TH2 |
| 1700021F05Rik | 4.55E-52 | 0.25281 | 0.725 | 0.623 | 5.20E-48 | TH2 |
| Prdx4 | 1.62E-51 | 0.264646 | 0.72 | 0.617 | 1.85E-47 | TH2 |
| Slc50a1 | 2.90E-50 | 0.254163 | 0.731 | 0.627 | 3.32E-46 | TH2 |
| Tsfm | 3.60E-49 | 0.254521 | 0.726 | 0.625 | 4.12E-45 | TH2 |
| Fcf1 | 3.95E-49 | 0.26341 | 0.722 | 0.617 | 4.52E-45 | TH2 |
| Fads3 | 7.57E-48 | 0.254595 | 0.703 | 0.596 | 8.66E-44 | TH2 |
| Wdr18 | 1.82E-47 | 0.273394 | 0.73 | 0.625 | 2.08E-43 | TH2 |
| Mtx1 | 3.83E-44 | 0.251241 | 0.695 | 0.599 | 4.38E-40 | TH2 |
| Depdc7 | 5.48E-44 | 0.280059 | 0.675 | 0.562 | 6.26E-40 | TH2 |
| Rce1 | 8.73E-43 | 0.267869 | 0.706 | 0.607 | 9.99E-39 | TH2 |
| Fbp2 | 1.53E-38 | 0.345265 | 0.651 | 0.561 | 1.75E-34 | TH2 |
| Tmem159 | 4.40E-28 | 0.280827 | 0.779 | 0.74 | 5.03E-24 | TH2 |
| Pkp3 | 1.69E-25 | 0.306107 | 0.646 | 0.564 | 1.93E-21 | TH2 |

| **Table S2: Analysis of TH1 vs. TH2 single-cell gene expression; a positive log2FC indicates higher expression in TH1 dorsal root ganglia neurons** | | | | | |
| --- | --- | --- | --- | --- | --- |
| Gene | p_val | avg_logFC | pct.1 | pct.2 | p_val_adj |
| Rps19 | 0 | 0.573924475 | 0.999 | 0.998 | 0 |
| Rpl41 | 0 | 0.500153608 | 1 | 1 | 0 |
| Fth1 | 0 | 0.480047554 | 1 | 1 | 0 |
| Stmn3 | 0 | -0.628129813 | 0.98 | 0.998 | 0 |
| Chchd2 | 0 | -0.645446233 | 0.983 | 1 | 0 |
| Atp5l | 0 | -0.707568055 | 0.968 | 1 | 0 |
| Uqcr11 | 0 | -0.715531187 | 0.956 | 0.997 | 0 |
| Cfl1 | 0 | -0.758991236 | 0.937 | 0.998 | 0 |
| Bex2 | 0 | -0.776061875 | 0.939 | 0.998 | 0 |
| Atp6v1f | 0 | -0.783961889 | 0.926 | 0.996 | 0 |
| Tuba1a | 0 | -0.804071377 | 0.986 | 0.997 | 0 |
| Cox7a2 | 0 | -0.810162394 | 0.959 | 0.999 | 0 |
| Uqcr10 | 0 | -0.818703796 | 0.893 | 0.998 | 0 |
| Rbx1 | 0 | -0.828397218 | 0.911 | 0.997 | 0 |
| Uchl1 | 0 | -0.842812479 | 0.983 | 1 | 0 |
| Stmn1 | 0 | -0.850013445 | 0.983 | 0.999 | 0 |
| Ndufs5 | 0 | -0.850533877 | 0.88 | 0.992 | 0 |
| Ppia | 0 | -0.851446722 | 0.998 | 1 | 0 |
| Gm7271 | 0 | -0.851760937 | 0.911 | 0.993 | 0 |
| Myl6 | 0 | -0.880583026 | 0.819 | 0.987 | 0 |
| Dctn3 | 0 | -0.881083861 | 0.877 | 0.993 | 0 |
| Atp5g1 | 0 | -0.887217269 | 0.961 | 0.999 | 0 |
| Cox6b1 | 0 | -0.888243259 | 0.951 | 0.999 | 0 |
| Cox5a | 0 | -0.888967397 | 0.928 | 0.999 | 0 |
| Dynll1 | 0 | -0.91890572 | 0.974 | 0.999 | 0 |
| Aldoa | 0 | -0.919039663 | 0.951 | 0.997 | 0 |
| Sncg | 0 | -0.929784482 | 1 | 1 | 0 |
| Psma7 | 0 | -0.929916183 | 0.854 | 0.993 | 0 |
| Atpif1 | 0 | -0.930811908 | 0.981 | 1 | 0 |
| Atp5j2 | 0 | -0.933565747 | 0.905 | 0.996 | 0 |
| Calm2 | 0 | -0.939539939 | 0.999 | 1 | 0 |
| Rabac1 | 0 | -0.968995559 | 0.871 | 0.995 | 0 |
| Ndufs7 | 0 | -0.969818157 | 0.737 | 0.979 | 0 |
| Calm1 | 0 | -0.972127815 | 0.991 | 1 | 0 |
| Anxa2 | 0 | -0.973573146 | 0.891 | 0.994 | 0 |
| Cox6a1 | 0 | -0.979258683 | 0.94 | 0.995 | 0 |
| Psmb3 | 0 | -0.997290641 | 0.755 | 0.986 | 0 |
| Ndufb4 | 0 | -1.006673613 | 0.825 | 0.994 | 0 |
| Spcs1 | 0 | -1.019860575 | 0.71 | 0.984 | 0 |
| Psmb6 | 0 | -1.022806581 | 0.791 | 0.992 | 0 |
| Uqcrb | 0 | -1.022844111 | 0.875 | 0.995 | 0 |
| Ndufb6 | 0 | -1.02968983 | 0.773 | 0.993 | 0 |
| Dad1 | 0 | -1.045304528 | 0.746 | 0.99 | 0 |
| Gabarapl2 | 0 | -1.073328466 | 0.921 | 0.999 | 0 |
| Mdh1 | 0 | -1.074910118 | 0.731 | 0.975 | 0 |
| Ndufab1 | 0 | -1.076710609 | 0.77 | 0.99 | 0 |
| Cd9 | 0 | -1.086745405 | 0.799 | 0.985 | 0 |
| Ndufb2 | 0 | -1.098936589 | 0.811 | 0.994 | 0 |
| Nop10 | 0 | -1.117311821 | 0.825 | 0.996 | 0 |
| Psmb2 | 0 | -1.133926144 | 0.811 | 0.982 | 0 |
| Ctsl | 0 | -1.144569222 | 0.747 | 0.968 | 0 |
| Ndufa11 | 0 | -1.166295896 | 0.865 | 0.997 | 0 |
| Atp6v0b | 0 | -1.168170019 | 0.875 | 0.999 | 0 |
| Tecr | 0 | -1.20159641 | 0.749 | 0.989 | 0 |
| Prdx1 | 0 | -1.201597075 | 0.857 | 0.996 | 0 |
| Hspb1 | 0 | -1.265847724 | 0.805 | 0.993 | 0 |
| Lxn | 0 | -1.295287906 | 0.858 | 0.996 | 0 |
| S100a10 | 0 | -1.30008047 | 0.952 | 0.999 | 0 |
| Txn1 | 0 | -1.309485344 | 0.946 | 0.999 | 0 |
| Mgst3 | 0 | -1.363437598 | 0.713 | 0.978 | 0 |
| Ndufa4 | 0 | -1.526808889 | 0.97 | 1 | 0 |
| Ubb | 0 | -1.748505444 | 0.955 | 0.999 | 0 |
| Nme1 | 6.09E-305 | -0.848252775 | 0.832 | 0.992 | 6.96E-301 |
| Anxa5 | 6.77E-305 | -0.674732636 | 0.939 | 0.997 | 7.74E-301 |
| Sh3bgrl3 | 1.40E-302 | -0.621340664 | 0.979 | 0.999 | 1.60E-298 |
| Rarres1 | 5.25E-302 | -0.768501813 | 0.851 | 0.997 | 6.00E-298 |
| Uqcrq | 6.00E-295 | -0.684621168 | 0.955 | 0.999 | 6.87E-291 |
| Atp5j | 3.93E-293 | -0.628924598 | 0.947 | 0.997 | 4.50E-289 |
| Tubb2a | 5.06E-290 | -1.021879193 | 0.726 | 0.965 | 5.79E-286 |
| Ldha | 1.66E-289 | -0.874313348 | 0.803 | 0.987 | 1.90E-285 |
| Cisd1 | 8.63E-289 | -0.750305725 | 0.89 | 0.994 | 9.88E-285 |
| Pfn1 | 5.78E-288 | -0.720053021 | 0.92 | 0.997 | 6.61E-284 |
| Tppp3 | 5.89E-287 | -0.501673766 | 1 | 1 | 6.74E-283 |
| Cst3 | 1.43E-286 | -0.742037173 | 0.802 | 0.996 | 1.63E-282 |
| Sdhb | 3.37E-282 | -0.914544992 | 0.73 | 0.972 | 3.85E-278 |
| Fez1 | 2.47E-280 | -0.788407731 | 0.821 | 0.977 | 2.82E-276 |
| Gapdh | 1.11E-277 | -0.768808495 | 0.897 | 0.99 | 1.27E-273 |
| Rbms3 | 1.20E-276 | 0.52418634 | 1 | 0.998 | 1.37E-272 |
| Ndufc2 | 1.69E-276 | -0.868815394 | 0.724 | 0.973 | 1.93E-272 |
| Cox6c | 7.07E-275 | -0.507238385 | 0.986 | 1 | 8.09E-271 |
| Vps29 | 9.86E-274 | -0.889290711 | 0.69 | 0.972 | 1.13E-269 |
| Hsp90aa1 | 7.15E-272 | -0.622061861 | 0.935 | 0.995 | 8.18E-268 |
| Prdx2 | 7.30E-271 | -0.781522751 | 0.829 | 0.988 | 8.36E-267 |
| Rpl13a | 8.67E-270 | 0.592934364 | 0.992 | 0.989 | 9.92E-266 |
| Rheb | 3.89E-267 | -0.791711126 | 0.822 | 0.989 | 4.45E-263 |
| Psmb5 | 4.75E-265 | -0.922869766 | 0.69 | 0.954 | 5.44E-261 |
| Cox7b | 4.17E-264 | -0.812038349 | 0.815 | 0.983 | 4.77E-260 |
| Calcb | 1.20E-263 | -1.053634564 | 0.84 | 0.976 | 1.37E-259 |
| Rpl3 | 2.52E-263 | 0.598967971 | 0.996 | 0.992 | 2.88E-259 |
| Rpl37 | 9.12E-263 | 0.503190783 | 0.998 | 0.998 | 1.04E-258 |
| Fkbp3 | 9.71E-263 | -0.883119657 | 0.692 | 0.967 | 1.11E-258 |
| Cox8a | 3.99E-260 | -0.437932428 | 0.998 | 1 | 4.57E-256 |
| Atp5g3 | 1.08E-259 | -0.75642579 | 0.853 | 0.988 | 1.23E-255 |
| Mif | 7.87E-258 | -0.828092796 | 0.724 | 0.977 | 9.00E-254 |
| Swi5 | 8.66E-258 | -0.791911906 | 0.791 | 0.981 | 9.91E-254 |
| Polr2g | 1.87E-257 | -0.920003868 | 0.64 | 0.942 | 2.14E-253 |
| Atp5o | 1.05E-255 | -0.740358897 | 0.846 | 0.986 | 1.20E-251 |
| Ndufb8 | 1.16E-253 | -0.582593621 | 0.961 | 0.997 | 1.32E-249 |
| Ywhaz | 1.20E-251 | 0.587508931 | 0.999 | 0.993 | 1.37E-247 |
| Tonsl | 4.00E-251 | -0.891304074 | 0.685 | 0.953 | 4.57E-247 |
| Nedd8 | 3.99E-250 | -0.613057262 | 0.95 | 0.997 | 4.57E-246 |
| Ftl1 | 9.26E-249 | 0.410550803 | 1 | 1 | 1.06E-244 |
| Zfhx3 | 5.07E-248 | 0.622487499 | 0.998 | 0.994 | 5.80E-244 |
| Tceb2 | 2.28E-246 | -0.568285879 | 0.971 | 0.996 | 2.61E-242 |
| Ndufa7 | 2.34E-246 | -0.760529755 | 0.842 | 0.984 | 2.68E-242 |
| Atp6v1e1 | 2.43E-244 | -0.83433308 | 0.688 | 0.956 | 2.78E-240 |
| Tagln3 | 9.27E-241 | -0.802836543 | 0.789 | 0.975 | 1.06E-236 |
| Prdx5 | 4.23E-240 | -0.701114646 | 0.872 | 0.989 | 4.84E-236 |
| Rpl13 | 6.42E-237 | 0.541710068 | 0.995 | 0.99 | 7.35E-233 |
| Rps23 | 2.73E-236 | 0.523146204 | 0.995 | 0.994 | 3.12E-232 |
| Rps5 | 5.70E-236 | 0.523246912 | 0.996 | 0.992 | 6.52E-232 |
| Usmg5 | 1.68E-234 | -0.613960616 | 0.906 | 0.994 | 1.92E-230 |
| Itm2b | 3.27E-233 | -0.688509003 | 0.835 | 0.991 | 3.75E-229 |
| Rps18 | 4.40E-233 | 0.545530621 | 0.995 | 0.989 | 5.03E-229 |
| Eif4a2 | 3.90E-231 | 0.514030476 | 0.999 | 0.995 | 4.46E-227 |
| Uqcrc1 | 2.27E-229 | -0.787669899 | 0.655 | 0.946 | 2.60E-225 |
| Rpl10 | 3.01E-229 | 0.527714778 | 0.995 | 0.992 | 3.45E-225 |
| Atp5f1 | 2.44E-227 | -0.738679814 | 0.77 | 0.975 | 2.79E-223 |
| Cstb | 3.44E-224 | -0.805743391 | 0.675 | 0.948 | 3.94E-220 |
| Hn1 | 2.16E-222 | -0.77685851 | 0.715 | 0.964 | 2.47E-218 |
| BC031181 | 1.84E-221 | -0.747434916 | 0.742 | 0.971 | 2.11E-217 |
| Map1lc3a | 8.64E-220 | -0.489421091 | 0.975 | 0.996 | 9.88E-216 |
| Pla2g7 | 1.96E-219 | -0.767369644 | 0.62 | 0.919 | 2.24E-215 |
| Atp5b | 1.31E-218 | -0.71611954 | 0.835 | 0.979 | 1.50E-214 |
| Psma2 | 1.33E-217 | -0.776409604 | 0.729 | 0.96 | 1.52E-213 |
| Ndufc1 | 1.15E-216 | -0.700569065 | 0.74 | 0.981 | 1.31E-212 |
| Mdh2 | 3.41E-216 | -0.745160894 | 0.749 | 0.969 | 3.90E-212 |
| Tubb4b | 3.49E-216 | -0.785056545 | 0.675 | 0.939 | 3.99E-212 |
| Ssu72 | 1.86E-215 | -0.780967277 | 0.664 | 0.939 | 2.12E-211 |
| Psma6 | 1.88E-215 | -0.798293311 | 0.746 | 0.918 | 2.15E-211 |
| Cox5b | 5.53E-215 | -0.488852582 | 0.966 | 0.997 | 6.33E-211 |
| Emc4 | 1.50E-214 | -0.778576243 | 0.63 | 0.924 | 1.71E-210 |
| Kif21a | 2.59E-214 | 0.481058154 | 0.999 | 0.998 | 2.96E-210 |
| Rps27 | 2.98E-214 | 0.548768677 | 0.995 | 0.991 | 3.41E-210 |
| Mpc1 | 5.98E-214 | -0.679574984 | 0.83 | 0.985 | 6.84E-210 |
| Ifi27 | 1.09E-212 | -0.870120825 | 0.65 | 0.927 | 1.25E-208 |
| Hagh | 1.14E-212 | -0.786947523 | 0.618 | 0.918 | 1.31E-208 |
| Pomp | 2.40E-212 | -0.666899015 | 0.839 | 0.989 | 2.75E-208 |
| Fabp5 | 6.45E-211 | -0.731543128 | 0.8 | 0.975 | 7.38E-207 |
| Rab3a | 4.17E-210 | -0.660556863 | 0.867 | 0.986 | 4.77E-206 |
| Cyb5a | 8.51E-210 | -0.76931581 | 0.713 | 0.958 | 9.74E-206 |
| Cd81 | 2.27E-208 | -0.765869151 | 0.584 | 0.944 | 2.59E-204 |
| Psmb1 | 1.46E-207 | -0.649424567 | 0.852 | 0.984 | 1.67E-203 |
| Ndufa5 | 5.99E-206 | -0.635309139 | 0.813 | 0.985 | 6.86E-202 |
| Dynlrb1 | 7.64E-206 | -0.534923446 | 0.947 | 0.997 | 8.74E-202 |
| Gstm5 | 1.59E-201 | -0.73401913 | 0.609 | 0.917 | 1.82E-197 |
| Snrpd3 | 2.50E-201 | -0.726602389 | 0.586 | 0.884 | 2.85E-197 |
| Ctsb | 2.56E-201 | -0.644491223 | 0.8 | 0.969 | 2.93E-197 |
| Higd2a | 4.05E-200 | -0.712835267 | 0.733 | 0.965 | 4.64E-196 |
| Psma1 | 1.21E-197 | -0.702096913 | 0.76 | 0.922 | 1.38E-193 |
| Crip2 | 2.29E-197 | -0.743508236 | 0.776 | 0.97 | 2.62E-193 |
| Hprt | 1.46E-196 | -0.68871774 | 0.75 | 0.971 | 1.68E-192 |
| Cdc42 | 1.77E-196 | -0.574087391 | 0.86 | 0.988 | 2.03E-192 |
| Ndufa12 | 5.45E-196 | -0.681769189 | 0.7 | 0.961 | 6.24E-192 |
| Emc9 | 4.52E-194 | -0.692752195 | 0.676 | 0.939 | 5.18E-190 |
| Rpl32 | 1.77E-193 | 0.52530831 | 0.99 | 0.985 | 2.03E-189 |
| Rps27a | 8.06E-193 | 0.451202564 | 0.996 | 0.997 | 9.22E-189 |
| Rpl27a | 1.15E-192 | 0.519054161 | 0.991 | 0.989 | 1.31E-188 |
| Rpl37a | 5.32E-192 | 0.449730348 | 0.995 | 0.996 | 6.09E-188 |
| Tshz2 | 9.62E-192 | 0.634938268 | 0.977 | 0.956 | 1.10E-187 |
| Bsg | 1.33E-191 | -0.674537806 | 0.735 | 0.967 | 1.53E-187 |
| Bcap31 | 1.35E-191 | -0.74878279 | 0.63 | 0.909 | 1.55E-187 |
| Tmem50a | 1.55E-191 | -0.656783519 | 0.785 | 0.973 | 1.77E-187 |
| Coa3 | 1.92E-191 | -0.645664144 | 0.817 | 0.978 | 2.20E-187 |
| Ndufs4 | 3.55E-191 | -0.685115826 | 0.63 | 0.919 | 4.06E-187 |
| Ndufa9 | 1.45E-188 | -0.696072424 | 0.573 | 0.872 | 1.66E-184 |
| Mcts1 | 1.63E-188 | -0.651425474 | 0.582 | 0.885 | 1.87E-184 |
| Tmem59 | 1.90E-188 | -0.671609466 | 0.768 | 0.968 | 2.18E-184 |
| Tmbim4 | 1.82E-186 | -0.683073092 | 0.608 | 0.899 | 2.08E-182 |
| Med21 | 2.54E-186 | -0.668089416 | 0.62 | 0.905 | 2.91E-182 |
| Cuta | 1.97E-184 | -0.675151438 | 0.63 | 0.901 | 2.25E-180 |
| Tbcb | 2.01E-184 | -0.658929088 | 0.774 | 0.961 | 2.29E-180 |
| Ubl5 | 5.00E-184 | -0.577228719 | 0.902 | 0.991 | 5.72E-180 |
| Ndufb11 | 8.78E-184 | -0.652973764 | 0.764 | 0.968 | 1.00E-179 |
| Mrpl41 | 4.27E-183 | -0.703934891 | 0.627 | 0.907 | 4.89E-179 |
| Snrpb | 4.55E-181 | -0.677997466 | 0.571 | 0.861 | 5.21E-177 |
| Ndufs8 | 2.08E-180 | -0.64787951 | 0.649 | 0.917 | 2.38E-176 |
| Ap2s1 | 8.47E-180 | -0.643794556 | 0.71 | 0.954 | 9.69E-176 |
| Fdps | 9.59E-179 | -0.698619719 | 0.628 | 0.922 | 1.10E-174 |
| Trappc2l | 1.08E-178 | -0.647721051 | 0.663 | 0.922 | 1.24E-174 |
| Camk2n1 | 1.74E-178 | 0.578205992 | 0.983 | 0.97 | 1.99E-174 |
| Taldo1 | 4.92E-178 | -0.68008113 | 0.577 | 0.869 | 5.63E-174 |
| Rpl9 | 6.61E-178 | 0.504165983 | 0.988 | 0.985 | 7.56E-174 |
| Psma4 | 4.01E-177 | -0.670018209 | 0.566 | 0.873 | 4.59E-173 |
| S100a13 | 4.53E-177 | -0.686448214 | 0.707 | 0.944 | 5.18E-173 |
| Ppp1r11 | 1.56E-176 | -0.696223378 | 0.68 | 0.935 | 1.79E-172 |
| Cycs | 1.75E-174 | -0.578755767 | 0.866 | 0.99 | 2.00E-170 |
| Selm | 2.56E-174 | -0.658191667 | 0.761 | 0.959 | 2.93E-170 |
| Gtf2h5 | 9.11E-174 | -0.643045057 | 0.741 | 0.953 | 1.04E-169 |
| Pdcd10 | 4.26E-172 | -0.646065291 | 0.68 | 0.939 | 4.88E-168 |
| Mpc2 | 1.01E-170 | -0.665555905 | 0.639 | 0.902 | 1.16E-166 |
| Tac1 | 4.96E-170 | -0.771898154 | 0.824 | 0.952 | 5.67E-166 |
| Psenen | 1.11E-169 | -0.659892142 | 0.668 | 0.93 | 1.27E-165 |
| Mrpl27 | 2.26E-169 | -0.63716197 | 0.642 | 0.911 | 2.59E-165 |
| Commd4 | 1.18E-166 | -0.59691247 | 0.748 | 0.887 | 1.35E-162 |
| H2-D1 | 1.54E-166 | -0.637043632 | 0.619 | 0.903 | 1.76E-162 |
| Rps8 | 2.10E-166 | 0.473620502 | 0.987 | 0.985 | 2.41E-162 |
| H2afz | 7.16E-166 | -0.572971862 | 0.856 | 0.989 | 8.20E-162 |
| Srp14 | 9.38E-166 | -0.538249287 | 0.887 | 0.99 | 1.07E-161 |
| Myl12b | 1.02E-165 | -0.628301399 | 0.712 | 0.952 | 1.17E-161 |
| Hint1 | 1.94E-165 | -0.393092251 | 0.987 | 0.997 | 2.22E-161 |
| Psma3 | 2.21E-165 | -0.630417512 | 0.735 | 0.959 | 2.53E-161 |
| Pitpnc1 | 2.95E-165 | 0.512633396 | 0.987 | 0.988 | 3.37E-161 |
| Zdhhc2 | 3.34E-165 | 0.472965933 | 0.991 | 0.992 | 3.82E-161 |
| Cox14 | 3.49E-165 | -0.609338038 | 0.784 | 0.97 | 3.99E-161 |
| Ndufb7 | 1.22E-164 | -0.57073457 | 0.86 | 0.98 | 1.40E-160 |
| Tmem256 | 2.08E-163 | -0.573561471 | 0.749 | 0.973 | 2.38E-159 |
| Hist1h2bc | 2.52E-162 | -0.629832434 | 0.764 | 0.965 | 2.88E-158 |
| Dpysl2 | 4.29E-162 | 0.479776195 | 0.995 | 0.994 | 4.91E-158 |
| Mrpl20 | 4.59E-162 | -0.608460311 | 0.695 | 0.938 | 5.25E-158 |
| Rpl18a | 3.81E-161 | 0.41076277 | 0.995 | 0.996 | 4.35E-157 |
| Ostf1 | 5.49E-161 | -0.653757427 | 0.81 | 0.973 | 6.28E-157 |
| Napa | 4.13E-160 | -0.61227706 | 0.68 | 0.929 | 4.73E-156 |
| Ndufv3 | 5.81E-160 | -0.519310072 | 0.859 | 0.986 | 6.65E-156 |
| Mien1 | 7.14E-160 | -0.575727139 | 0.81 | 0.976 | 8.17E-156 |
| Ndufa1 | 9.00E-160 | -0.601028957 | 0.617 | 0.937 | 1.03E-155 |
| Tsen34 | 3.98E-159 | -0.603749856 | 0.752 | 0.96 | 4.55E-155 |
| Piezo2 | 4.01E-159 | 0.521818454 | 0.978 | 0.984 | 4.58E-155 |
| Mrps14 | 4.19E-157 | -0.639561386 | 0.624 | 0.896 | 4.80E-153 |
| Erh | 5.66E-155 | -0.609015519 | 0.685 | 0.929 | 6.48E-151 |
| 0610009B22Rik | 6.71E-155 | -0.642307111 | 0.768 | 0.911 | 7.67E-151 |
| Spag7 | 2.30E-154 | -0.587036757 | 0.643 | 0.903 | 2.63E-150 |
| Rpl26 | 2.69E-154 | 0.415707957 | 0.995 | 0.992 | 3.08E-150 |
| Rps14 | 4.36E-154 | 0.368442053 | 0.998 | 0.998 | 4.99E-150 |
| Ndufs3 | 4.41E-154 | -0.607207797 | 0.656 | 0.921 | 5.04E-150 |
| Atp6v1g1 | 7.07E-154 | -0.459710871 | 0.949 | 0.995 | 8.09E-150 |
| S100a6 | 2.72E-153 | -0.870074356 | 0.885 | 0.946 | 3.11E-149 |
| Pmm1 | 7.69E-153 | -0.569515417 | 0.83 | 0.978 | 8.80E-149 |
| Mrpl36 | 7.89E-153 | -0.620316697 | 0.567 | 0.845 | 9.03E-149 |
| Akr1a1 | 6.58E-152 | -0.543816657 | 0.845 | 0.977 | 7.53E-148 |
| Aurkaip1 | 7.85E-152 | -0.591812225 | 0.759 | 0.956 | 8.98E-148 |
| Pttg1 | 7.55E-151 | -0.660943014 | 0.756 | 0.899 | 8.63E-147 |
| Tpi1 | 1.36E-150 | -0.574008212 | 0.697 | 0.928 | 1.56E-146 |
| Eef1a1 | 2.22E-150 | 0.5080108 | 0.974 | 0.976 | 2.54E-146 |
| Mrps33 | 2.92E-150 | -0.56701738 | 0.82 | 0.977 | 3.34E-146 |
| Rps29 | 6.06E-150 | 0.421773054 | 0.998 | 0.997 | 6.94E-146 |
| Vdac3 | 1.15E-148 | -0.592691583 | 0.636 | 0.902 | 1.32E-144 |
| Prr13 | 2.02E-148 | -0.427082662 | 0.974 | 0.999 | 2.31E-144 |
| Chchd10 | 6.42E-148 | -0.5528754 | 0.743 | 0.959 | 7.35E-144 |
| Cope | 9.09E-148 | -0.631001801 | 0.619 | 0.884 | 1.04E-143 |
| BC048546 | 1.28E-147 | -0.5641217 | 0.836 | 0.98 | 1.46E-143 |
| Cystm1 | 2.97E-147 | -0.596614161 | 0.641 | 0.912 | 3.39E-143 |
| Psmc2 | 6.03E-147 | -0.583238913 | 0.745 | 0.879 | 6.90E-143 |
| Ndufa13 | 6.38E-147 | -0.426630123 | 0.958 | 0.998 | 7.29E-143 |
| Hsbp1 | 9.62E-147 | -0.560814175 | 0.782 | 0.97 | 1.10E-142 |
| Rps4x | 2.56E-146 | 0.496296719 | 0.981 | 0.976 | 2.93E-142 |
| Mrpl28 | 7.16E-146 | -0.559418195 | 0.635 | 0.893 | 8.19E-142 |
| Rpl11 | 1.28E-145 | 0.478160083 | 0.99 | 0.981 | 1.46E-141 |
| Edf1 | 4.42E-145 | -0.501681425 | 0.893 | 0.988 | 5.06E-141 |
| Gadd45gip1 | 2.27E-144 | -0.566079967 | 0.642 | 0.894 | 2.59E-140 |
| Psmc4 | 8.08E-144 | -0.593592281 | 0.571 | 0.841 | 9.24E-140 |
| Romo1 | 9.70E-144 | -0.516833608 | 0.779 | 0.964 | 1.11E-139 |
| Rpp21 | 1.04E-143 | -0.588267197 | 0.593 | 0.855 | 1.19E-139 |
| Rps6 | 1.14E-143 | 0.419974918 | 0.992 | 0.99 | 1.31E-139 |
| Nt5c | 1.22E-143 | -0.582157042 | 0.637 | 0.905 | 1.40E-139 |
| Fau | 2.75E-143 | 0.370444215 | 0.996 | 0.996 | 3.14E-139 |
| Med31 | 7.68E-143 | -0.521107266 | 0.585 | 0.845 | 8.79E-139 |
| Trappc4 | 7.72E-143 | -0.566293798 | 0.774 | 0.966 | 8.84E-139 |
| Rpl14 | 1.05E-142 | 0.453207042 | 0.991 | 0.99 | 1.20E-138 |
| Cyc1 | 1.62E-142 | -0.574215142 | 0.688 | 0.919 | 1.85E-138 |
| Ndufa8 | 2.13E-141 | -0.5250557 | 0.793 | 0.968 | 2.44E-137 |
| Ywhag | 3.17E-141 | 0.455590563 | 0.995 | 0.989 | 3.62E-137 |
| Nkiras1 | 3.96E-141 | -0.540992855 | 0.668 | 0.903 | 4.53E-137 |
| Mrpl55 | 3.99E-141 | -0.528306663 | 0.61 | 0.871 | 4.57E-137 |
| Acot7 | 2.22E-140 | -0.561801794 | 0.735 | 0.947 | 2.54E-136 |
| Rps16 | 7.75E-140 | 0.447617584 | 0.988 | 0.99 | 8.86E-136 |
| Fabp3 | 8.62E-140 | -0.551750325 | 0.589 | 0.851 | 9.87E-136 |
| Mtx2 | 1.19E-139 | -0.559131518 | 0.573 | 0.841 | 1.36E-135 |
| Chmp2a | 1.62E-139 | -0.565475576 | 0.63 | 0.882 | 1.86E-135 |
| Arl6ip5 | 1.77E-139 | -0.587661998 | 0.689 | 0.929 | 2.02E-135 |
| Chmp5 | 4.82E-139 | -0.546059246 | 0.77 | 0.968 | 5.52E-135 |
| Apoa1bp | 6.86E-139 | -0.53307718 | 0.619 | 0.88 | 7.85E-135 |
| Hcfc1r1 | 2.55E-138 | -0.524954809 | 0.847 | 0.978 | 2.92E-134 |
| Mrps12 | 4.44E-138 | -0.568238686 | 0.707 | 0.931 | 5.08E-134 |
| Minos1 | 8.28E-138 | -0.454320789 | 0.932 | 0.993 | 9.47E-134 |
| Sf3b5 | 8.44E-138 | -0.532531627 | 0.634 | 0.891 | 9.66E-134 |
| 2010107E04Rik | 2.84E-137 | -0.476421524 | 0.922 | 0.993 | 3.25E-133 |
| Mrps17 | 4.90E-137 | -0.519393663 | 0.589 | 0.849 | 5.61E-133 |
| Med28 | 6.26E-137 | -0.553535723 | 0.686 | 0.941 | 7.17E-133 |
| Lamtor5 | 7.28E-137 | -0.535537948 | 0.711 | 0.948 | 8.33E-133 |
| Atp5d | 1.28E-136 | -0.485689717 | 0.89 | 0.986 | 1.47E-132 |
| Eif3g | 1.90E-136 | -0.551700203 | 0.615 | 0.868 | 2.17E-132 |
| Pfdn6 | 2.27E-136 | -0.55481963 | 0.629 | 0.888 | 2.60E-132 |
| Tspan3 | 4.21E-136 | -0.534992601 | 0.622 | 0.886 | 4.82E-132 |
| Ank2 | 2.77E-135 | 0.475730338 | 0.985 | 0.983 | 3.17E-131 |
| Phf24 | 2.61E-134 | 0.473717451 | 0.991 | 0.992 | 2.99E-130 |
| Tmem205 | 2.65E-134 | -0.574250106 | 0.748 | 0.878 | 3.04E-130 |
| Rpl6 | 5.78E-134 | 0.45507317 | 0.982 | 0.979 | 6.61E-130 |
| Sf3b6 | 6.20E-134 | -0.561654086 | 0.561 | 0.823 | 7.09E-130 |
| Atg101 | 1.26E-133 | -0.565614917 | 0.543 | 0.806 | 1.45E-129 |
| Timm13 | 3.13E-133 | -0.477126812 | 0.885 | 0.987 | 3.58E-129 |
| Mrps34 | 4.00E-133 | -0.511972129 | 0.593 | 0.845 | 4.57E-129 |
| Ube2v2 | 4.01E-133 | -0.546694958 | 0.75 | 0.961 | 4.59E-129 |
| Eno1 | 4.58E-133 | -0.502512993 | 0.648 | 0.901 | 5.24E-129 |
| Ckb | 4.88E-133 | -0.523121461 | 0.669 | 0.91 | 5.58E-129 |
| Tmem59l | 1.25E-132 | -0.506346587 | 0.6 | 0.844 | 1.43E-128 |
| Timm17a | 7.06E-132 | -0.545697655 | 0.714 | 0.942 | 8.08E-128 |
| Mbnl2 | 2.19E-131 | 0.47371203 | 0.983 | 0.984 | 2.50E-127 |
| Mllt11 | 4.87E-131 | -0.452297822 | 0.924 | 0.992 | 5.57E-127 |
| Dbi | 5.01E-131 | -0.525483425 | 0.633 | 0.897 | 5.74E-127 |
| Sdhd | 5.51E-131 | -0.551133347 | 0.618 | 0.876 | 6.31E-127 |
| Sri | 1.10E-130 | -0.519762343 | 0.707 | 0.932 | 1.26E-126 |
| Acot13 | 1.84E-130 | -0.50994329 | 0.616 | 0.866 | 2.10E-126 |
| Ndufs6 | 1.95E-130 | -0.490572934 | 0.862 | 0.981 | 2.23E-126 |
| Supt4a | 9.29E-130 | -0.508624043 | 0.642 | 0.895 | 1.06E-125 |
| Mrps25 | 2.36E-129 | -0.495963764 | 0.603 | 0.856 | 2.70E-125 |
| Psmb4 | 2.45E-129 | -0.527297966 | 0.713 | 0.928 | 2.80E-125 |
| Mrps6 | 3.51E-129 | -0.460338946 | 0.582 | 0.839 | 4.01E-125 |
| Mrpl51 | 3.81E-129 | -0.472889017 | 0.642 | 0.891 | 4.36E-125 |
| Ppib | 9.12E-129 | -0.473819993 | 0.597 | 0.849 | 1.04E-124 |
| Polr2j | 1.18E-128 | -0.495420342 | 0.621 | 0.874 | 1.35E-124 |
| 1110001J03Rik | 1.81E-128 | -0.500380703 | 0.589 | 0.85 | 2.07E-124 |
| Ldhb | 3.21E-128 | -0.500296673 | 0.571 | 0.807 | 3.67E-124 |
| Mrpl21 | 1.83E-127 | -0.505535236 | 0.617 | 0.861 | 2.10E-123 |
| Rps20 | 4.40E-127 | 0.494444835 | 0.974 | 0.963 | 5.03E-123 |
| Pin1 | 7.70E-127 | -0.494518091 | 0.674 | 0.91 | 8.81E-123 |
| Yipf4 | 9.09E-127 | -0.50379414 | 0.697 | 0.93 | 1.04E-122 |
| Higd1a | 1.20E-126 | -0.525189228 | 0.782 | 0.962 | 1.38E-122 |
| Sncb | 2.63E-126 | -0.522210062 | 0.751 | 0.942 | 3.01E-122 |
| Cct7 | 2.71E-126 | -0.521146884 | 0.622 | 0.859 | 3.10E-122 |
| Dhrs7 | 2.94E-126 | -0.551602414 | 0.545 | 0.804 | 3.36E-122 |
| Park7 | 8.01E-126 | -0.509934065 | 0.774 | 0.96 | 9.17E-122 |
| Med29 | 1.28E-125 | -0.479902519 | 0.552 | 0.793 | 1.47E-121 |
| Flot1 | 1.54E-124 | -0.520658954 | 0.678 | 0.919 | 1.76E-120 |
| Apoo | 7.77E-124 | -0.492842809 | 0.736 | 0.835 | 8.89E-120 |
| Ethe1 | 9.57E-124 | -0.497645001 | 0.559 | 0.804 | 1.10E-119 |
| Dpy30 | 1.94E-123 | -0.509094118 | 0.673 | 0.921 | 2.22E-119 |
| Lgals1 | 3.68E-123 | -0.617582199 | 0.794 | 0.949 | 4.21E-119 |
| Tmem208 | 4.10E-123 | -0.483396019 | 0.605 | 0.853 | 4.69E-119 |
| Etfb | 6.88E-123 | -0.521291926 | 0.583 | 0.836 | 7.88E-119 |
| Meg3 | 9.00E-123 | 0.498080523 | 0.989 | 0.98 | 1.03E-118 |
| Shfm1 | 1.30E-122 | -0.401589403 | 0.92 | 0.993 | 1.49E-118 |
| Trappc1 | 1.08E-121 | -0.524385134 | 0.659 | 0.905 | 1.24E-117 |
| Pgp | 1.26E-121 | -0.513698148 | 0.723 | 0.931 | 1.44E-117 |
| Ptma | 1.56E-121 | 0.388841672 | 0.988 | 0.992 | 1.78E-117 |
| Cdkn2d | 2.22E-121 | -0.496736322 | 0.684 | 0.905 | 2.54E-117 |
| Vimp | 2.23E-121 | -0.53113008 | 0.595 | 0.847 | 2.55E-117 |
| 1110008F13Rik | 5.09E-121 | -0.463163507 | 0.605 | 0.857 | 5.82E-117 |
| Suclg1 | 8.67E-121 | -0.477408672 | 0.569 | 0.811 | 9.92E-117 |
| Naa38 | 1.09E-120 | -0.491926673 | 0.583 | 0.907 | 1.25E-116 |
| Coq7 | 1.30E-120 | -0.463104428 | 0.575 | 0.805 | 1.49E-116 |
| Ndufb3 | 3.22E-120 | -0.468975944 | 0.77 | 0.961 | 3.69E-116 |
| Psmb7 | 8.44E-120 | -0.482938145 | 0.712 | 0.931 | 9.65E-116 |
| Rps2 | 4.20E-119 | 0.466496411 | 0.985 | 0.982 | 4.80E-115 |
| Psmd14 | 6.21E-119 | -0.48345689 | 0.614 | 0.847 | 7.11E-115 |
| N6amt2 | 1.16E-118 | -0.468269099 | 0.738 | 0.839 | 1.33E-114 |
| Pebp1 | 1.31E-118 | -0.489087945 | 0.783 | 0.961 | 1.50E-114 |
| Timm10 | 2.14E-118 | -0.475243415 | 0.615 | 0.853 | 2.45E-114 |
| Zmat5 | 9.40E-118 | -0.515700247 | 0.586 | 0.83 | 1.08E-113 |
| Tmem14c | 9.55E-118 | -0.452288317 | 0.558 | 0.794 | 1.09E-113 |
| Rpl17 | 1.40E-117 | 0.511489074 | 0.975 | 0.965 | 1.60E-113 |
| Polr2k | 3.06E-117 | -0.519089879 | 0.778 | 0.968 | 3.50E-113 |
| Tmem60 | 3.07E-117 | -0.49409913 | 0.653 | 0.894 | 3.51E-113 |
| Acyp2 | 3.61E-117 | -0.491776075 | 0.608 | 0.844 | 4.13E-113 |
| Pop5 | 5.50E-117 | -0.47966582 | 0.708 | 0.92 | 6.29E-113 |
| Dnajc15 | 4.12E-116 | -0.4982954 | 0.608 | 0.857 | 4.71E-112 |
| Rpl36 | 1.45E-115 | 0.421537743 | 0.98 | 0.979 | 1.66E-111 |
| Cdkn1a | 2.91E-115 | -0.510972382 | 0.751 | 0.946 | 3.33E-111 |
| Psmd6 | 7.28E-115 | -0.489461074 | 0.596 | 0.843 | 8.33E-111 |
| Ten1 | 8.59E-115 | -0.407472679 | 0.593 | 0.827 | 9.83E-111 |
| Pdlim7 | 2.42E-114 | -0.445044329 | 0.656 | 0.886 | 2.77E-110 |
| Elavl2 | 2.48E-114 | 0.4620785 | 0.978 | 0.978 | 2.84E-110 |
| 0610011F06Rik | 2.84E-114 | -0.44303243 | 0.55 | 0.796 | 3.25E-110 |
| Timm22 | 1.69E-113 | -0.394639644 | 0.579 | 0.803 | 1.93E-109 |
| Mrpl34 | 1.72E-113 | -0.47439443 | 0.59 | 0.823 | 1.97E-109 |
| Eif6 | 6.54E-113 | -0.472035915 | 0.619 | 0.86 | 7.48E-109 |
| Nfu1 | 1.11E-112 | -0.492868478 | 0.621 | 0.864 | 1.28E-108 |
| Bola1 | 1.62E-112 | -0.463892267 | 0.559 | 0.787 | 1.86E-108 |
| Dhrs1 | 1.65E-112 | -0.469107604 | 0.612 | 0.851 | 1.89E-108 |
| Commd1 | 2.01E-112 | -0.418995809 | 0.578 | 0.818 | 2.30E-108 |
| Tmem160 | 3.10E-112 | -0.459288925 | 0.815 | 0.965 | 3.55E-108 |
| Arl2 | 3.45E-112 | -0.427504352 | 0.609 | 0.847 | 3.95E-108 |
| Atox1 | 4.69E-112 | -0.475389899 | 0.815 | 0.962 | 5.37E-108 |
| Snrpc | 5.11E-112 | -0.448009225 | 0.554 | 0.788 | 5.84E-108 |
| Pcmt1 | 9.03E-112 | -0.468513096 | 0.758 | 0.961 | 1.03E-107 |
| Atp6v1d | 1.38E-111 | -0.470982256 | 0.68 | 0.901 | 1.58E-107 |
| Rpl36a | 1.99E-111 | 0.487644893 | 0.958 | 0.961 | 2.28E-107 |
| Rsu1 | 2.95E-111 | -0.468411932 | 0.833 | 0.947 | 3.37E-107 |
| Zwint | 3.87E-111 | -0.445076965 | 0.885 | 0.98 | 4.43E-107 |
| Sssca1 | 3.97E-111 | -0.458759652 | 0.559 | 0.792 | 4.54E-107 |
| Naa20 | 4.43E-111 | -0.517005417 | 0.558 | 0.804 | 5.07E-107 |
| Acyp1 | 6.19E-111 | -0.485057394 | 0.55 | 0.79 | 7.08E-107 |
| Dohh | 1.44E-110 | -0.479279534 | 0.641 | 0.87 | 1.65E-106 |
| Acot9 | 2.39E-110 | -0.407726454 | 0.548 | 0.785 | 2.74E-106 |
| Ndufb9 | 3.33E-110 | -0.395516459 | 0.936 | 0.994 | 3.81E-106 |
| Tm2d2 | 1.61E-109 | -0.439012374 | 0.607 | 0.843 | 1.84E-105 |
| Lamtor4 | 1.79E-109 | -0.45580341 | 0.685 | 0.908 | 2.04E-105 |
| Anapc11 | 2.26E-109 | -0.468373344 | 0.822 | 0.97 | 2.59E-105 |
| Pde6d | 2.58E-109 | -0.478361605 | 0.545 | 0.827 | 2.95E-105 |
| Slc25a3 | 3.06E-109 | -0.420928847 | 0.91 | 0.981 | 3.50E-105 |
| Commd3 | 3.66E-109 | -0.44482648 | 0.581 | 0.832 | 4.18E-105 |
| Mrpl35 | 4.18E-109 | -0.458068564 | 0.572 | 0.809 | 4.78E-105 |
| Isl2 | 7.27E-109 | -0.459194449 | 0.816 | 0.964 | 8.31E-105 |
| Rpl35a | 8.40E-109 | 0.420046478 | 0.985 | 0.985 | 9.61E-105 |
| Ndufv1 | 2.68E-108 | -0.457660861 | 0.681 | 0.903 | 3.06E-104 |
| Emc7 | 3.90E-108 | -0.447855545 | 0.631 | 0.865 | 4.46E-104 |
| Rps3a1 | 5.60E-108 | 0.447311611 | 0.965 | 0.962 | 6.41E-104 |
| Fam173a | 8.88E-108 | -0.447802121 | 0.665 | 0.899 | 1.02E-103 |
| Scg5 | 1.27E-107 | -0.440155276 | 0.558 | 0.792 | 1.45E-103 |
| Cuedc2 | 1.27E-107 | -0.4728938 | 0.747 | 0.95 | 1.45E-103 |
| Avpi1 | 1.78E-107 | -0.440430316 | 0.6 | 0.811 | 2.04E-103 |
| Rpl8 | 6.84E-107 | 0.45370016 | 0.973 | 0.965 | 7.83E-103 |
| Pdzd11 | 6.91E-107 | -0.465407987 | 0.545 | 0.786 | 7.90E-103 |
| Grpel1 | 7.90E-107 | -0.435471508 | 0.565 | 0.793 | 9.04E-103 |
| Pou4f1 | 1.02E-106 | 0.397043151 | 0.99 | 0.99 | 1.17E-102 |
| Rps7 | 1.07E-106 | 0.437011151 | 0.972 | 0.969 | 1.22E-102 |
| Cacybp | 3.85E-106 | -0.448345747 | 0.68 | 0.908 | 4.41E-102 |
| Cbr1 | 6.91E-106 | -0.416717955 | 0.554 | 0.784 | 7.91E-102 |
| Eif3f | 1.22E-105 | 0.492864554 | 0.965 | 0.96 | 1.39E-101 |
| Arpc3 | 1.26E-105 | -0.479685095 | 0.684 | 0.9 | 1.44E-101 |
| Chchd1 | 1.33E-105 | -0.458622802 | 0.667 | 0.9 | 1.52E-101 |
| Tceb1 | 1.61E-105 | -0.443499289 | 0.837 | 0.981 | 1.85E-101 |
| Polr2e | 1.63E-105 | -0.435536352 | 0.574 | 0.807 | 1.87E-101 |
| Idh3g | 2.12E-105 | -0.360559942 | 0.54 | 0.754 | 2.42E-101 |
| Lgmn | 2.42E-105 | -0.422138184 | 0.478 | 0.818 | 2.77E-101 |
| Ttll7 | 2.47E-105 | 0.303610995 | 0.999 | 1 | 2.83E-101 |
| Eef1g | 2.82E-105 | 0.491522093 | 0.967 | 0.962 | 3.23E-101 |
| Rps15a | 3.55E-105 | 0.418502161 | 0.975 | 0.979 | 4.06E-101 |
| Snrpd1 | 4.65E-105 | -0.431894469 | 0.614 | 0.842 | 5.32E-101 |
| Tm2d3 | 5.24E-105 | -0.453672234 | 0.548 | 0.771 | 5.99E-101 |
| Sec13 | 7.48E-105 | -0.422477102 | 0.543 | 0.762 | 8.56E-101 |
| Klf7 | 4.54E-104 | 0.376607281 | 0.986 | 0.992 | 5.19E-100 |
| Snca | 7.70E-104 | -0.504651237 | 0.633 | 0.852 | 8.81E-100 |
| Aamp | 1.06E-103 | -0.423959019 | 0.588 | 0.819 | 1.21E-99 |
| Psma5 | 1.63E-103 | -0.474343297 | 0.7 | 0.911 | 1.87E-99 |
| Ubash3b | 2.00E-103 | 0.636900333 | 0.865 | 0.837 | 2.29E-99 |
| Ran | 2.51E-103 | -0.429148415 | 0.788 | 0.963 | 2.87E-99 |
| Rps11 | 3.85E-103 | 0.338353732 | 0.995 | 0.994 | 4.41E-99 |
| Atp6v0d1 | 5.76E-103 | -0.451476289 | 0.676 | 0.904 | 6.59E-99 |
| Ndufv2 | 5.91E-103 | -0.41832834 | 0.65 | 0.879 | 6.76E-99 |
| Txnl1 | 7.87E-103 | -0.431011436 | 0.676 | 0.901 | 9.00E-99 |
| Tmem258 | 1.21E-102 | -0.449582372 | 0.565 | 0.882 | 1.39E-98 |
| Pcbp2 | 1.22E-102 | 0.463714833 | 0.958 | 0.958 | 1.39E-98 |
| Emc2 | 2.12E-102 | -0.402699087 | 0.613 | 0.844 | 2.43E-98 |
| Tubb5 | 3.31E-102 | 0.282437646 | 0.998 | 0.997 | 3.79E-98 |
| Mrps21 | 5.43E-102 | -0.457876059 | 0.623 | 0.901 | 6.21E-98 |
| Rpl31 | 8.62E-102 | 0.422501828 | 0.974 | 0.973 | 9.87E-98 |
| Rpl34 | 2.53E-101 | 0.369621568 | 0.987 | 0.989 | 2.90E-97 |
| Yipf1 | 3.90E-101 | -0.402461758 | 0.533 | 0.755 | 4.46E-97 |
| Akr1b3 | 4.54E-101 | -0.437621092 | 0.601 | 0.826 | 5.19E-97 |
| Ppa1 | 5.87E-101 | -0.409494234 | 0.574 | 0.811 | 6.71E-97 |
| Laptm4a | 1.07E-100 | -0.433638965 | 0.67 | 0.931 | 1.23E-96 |
| Mtch2 | 1.22E-100 | -0.424479045 | 0.654 | 0.892 | 1.40E-96 |
| Ndufa2 | 1.44E-100 | -0.362059893 | 0.93 | 0.992 | 1.64E-96 |
| Manf | 1.67E-100 | -0.427276152 | 0.685 | 0.912 | 1.91E-96 |
| Slc25a11 | 3.53E-100 | -0.372891568 | 0.595 | 0.798 | 4.04E-96 |
| Tex264 | 7.92E-100 | -0.441246382 | 0.607 | 0.847 | 9.06E-96 |
| Atp5h | 1.07E-99 | -0.349134838 | 0.942 | 0.989 | 1.22E-95 |
| Oaz1 | 1.10E-99 | -0.261221496 | 0.995 | 1 | 1.26E-95 |
| Tmem14a | 2.27E-99 | -0.415304524 | 0.609 | 0.836 | 2.60E-95 |
| Rprm | 3.41E-99 | -0.460372982 | 0.756 | 0.945 | 3.90E-95 |
| l7Rn6 | 3.88E-99 | -0.414522277 | 0.6 | 0.832 | 4.44E-95 |
| Atp5k | 4.82E-99 | -0.378945283 | 0.927 | 0.993 | 5.52E-95 |
| Ech1 | 8.83E-99 | -0.384735757 | 0.555 | 0.773 | 1.01E-94 |
| Ebf1 | 1.28E-98 | 0.413075985 | 0.974 | 0.978 | 1.47E-94 |
| Ranbp1 | 1.60E-98 | -0.427742635 | 0.755 | 0.952 | 1.83E-94 |
| Scn7a | 1.70E-98 | 0.374255363 | 0.985 | 0.988 | 1.94E-94 |
| Psmd8 | 1.98E-98 | -0.410438196 | 0.594 | 0.816 | 2.26E-94 |
| Rps27l | 2.24E-98 | -0.440581901 | 0.697 | 0.947 | 2.56E-94 |
| Nipsnap3b | 2.82E-98 | -0.407787132 | 0.613 | 0.848 | 3.23E-94 |
| Slirp | 3.28E-98 | -0.415074305 | 0.675 | 0.898 | 3.75E-94 |
| Carkd | 2.68E-97 | -0.394688703 | 0.54 | 0.758 | 3.06E-93 |
| Coro1a | 8.03E-97 | -0.371475827 | 0.563 | 0.763 | 9.18E-93 |
| Map1lc3b | 1.43E-96 | -0.433980928 | 0.83 | 0.965 | 1.63E-92 |
| Frg1 | 2.50E-96 | -0.395212528 | 0.555 | 0.781 | 2.86E-92 |
| Tpgs1 | 2.95E-96 | -0.416630114 | 0.664 | 0.877 | 3.37E-92 |
| Rnh1 | 3.05E-96 | -0.430320293 | 0.594 | 0.827 | 3.49E-92 |
| Rraga | 3.84E-96 | -0.413961716 | 0.663 | 0.884 | 4.39E-92 |
| Ccdc109b | 3.95E-96 | -0.412272158 | 0.663 | 0.871 | 4.51E-92 |
| Rplp0 | 4.51E-96 | 0.443179068 | 0.96 | 0.959 | 5.16E-92 |
| Fkbp1a | 1.30E-95 | -0.30316885 | 0.965 | 0.996 | 1.49E-91 |
| Lrpap1 | 2.28E-95 | -0.324259228 | 0.577 | 0.787 | 2.61E-91 |
| Sra1 | 2.82E-95 | -0.391960061 | 0.614 | 0.83 | 3.22E-91 |
| Prmt1 | 3.07E-95 | -0.428144947 | 0.574 | 0.802 | 3.51E-91 |
| Tspo | 4.29E-95 | -0.685468228 | 0.571 | 0.732 | 4.91E-91 |
| Znhit3 | 4.46E-95 | -0.392583167 | 0.55 | 0.768 | 5.10E-91 |
| Plpp1 | 7.74E-95 | -0.432389895 | 0.536 | 0.754 | 8.85E-91 |
| Timm8b | 1.00E-94 | -0.434813258 | 0.677 | 0.924 | 1.15E-90 |
| Ndufa10 | 1.20E-94 | -0.438009503 | 0.731 | 0.922 | 1.37E-90 |
| Jagn1 | 1.28E-94 | -0.451292299 | 0.575 | 0.8 | 1.47E-90 |
| Ciapin1 | 1.43E-94 | -0.410616384 | 0.58 | 0.811 | 1.63E-90 |
| Pdrg1 | 1.50E-94 | -0.433817366 | 0.635 | 0.867 | 1.72E-90 |
| Akap12 | 1.82E-94 | 0.466563725 | 0.942 | 0.936 | 2.08E-90 |
| D10Jhu81e | 1.85E-94 | -0.388058936 | 0.554 | 0.767 | 2.12E-90 |
| Mrpl33 | 4.42E-94 | -0.415323618 | 0.658 | 0.876 | 5.06E-90 |
| Ict1 | 7.95E-94 | -0.36562032 | 0.615 | 0.844 | 9.09E-90 |
| Bud31 | 8.96E-94 | -0.388710894 | 0.558 | 0.774 | 1.02E-89 |
| Cyb561d2 | 9.54E-94 | -0.371431878 | 0.548 | 0.759 | 1.09E-89 |
| Map4 | 1.62E-93 | 0.406989619 | 0.975 | 0.976 | 1.85E-89 |
| Rnf7 | 1.98E-93 | -0.333617049 | 0.961 | 0.995 | 2.26E-89 |
| Adrm1 | 2.54E-93 | -0.388314395 | 0.565 | 0.801 | 2.91E-89 |
| Spcs2 | 3.12E-93 | -0.410274119 | 0.741 | 0.942 | 3.57E-89 |
| Morf4l1 | 3.62E-93 | 0.329486954 | 0.995 | 0.994 | 4.15E-89 |
| Ociad1 | 4.67E-93 | -0.405262915 | 0.651 | 0.874 | 5.35E-89 |
| Coa6 | 7.64E-93 | -0.360848725 | 0.619 | 0.838 | 8.74E-89 |
| Alkbh7 | 9.11E-93 | -0.401503915 | 0.541 | 0.755 | 1.04E-88 |
| Cebpzos | 1.36E-92 | -0.378523123 | 0.615 | 0.839 | 1.56E-88 |
| Eif4g2 | 1.53E-92 | 0.464712959 | 0.946 | 0.941 | 1.75E-88 |
| Magoh | 1.57E-92 | -0.417915411 | 0.732 | 0.82 | 1.80E-88 |
| Sec11a | 3.22E-92 | -0.441988042 | 0.77 | 0.864 | 3.69E-88 |
| Rpl24 | 4.48E-92 | 0.390515173 | 0.978 | 0.977 | 5.13E-88 |
| Scamp3 | 1.04E-91 | -0.384627413 | 0.574 | 0.789 | 1.19E-87 |
| Cnbp | 1.04E-91 | -0.368611219 | 0.623 | 0.849 | 1.20E-87 |
| Sar1b | 1.46E-91 | -0.41421367 | 0.562 | 0.794 | 1.67E-87 |
| Mrpl12 | 3.98E-91 | -0.37196985 | 0.644 | 0.845 | 4.55E-87 |
| Ddt | 7.55E-91 | -0.376038554 | 0.636 | 0.854 | 8.64E-87 |
| Bcas2 | 7.94E-91 | -0.401707395 | 0.706 | 0.917 | 9.08E-87 |
| Mrpl42 | 8.92E-91 | -0.40016747 | 0.709 | 0.922 | 1.02E-86 |
| Unc50 | 9.63E-91 | -0.396545897 | 0.59 | 0.825 | 1.10E-86 |
| Hexb | 1.15E-90 | -0.37356684 | 0.763 | 0.868 | 1.32E-86 |
| Hspbp1 | 1.26E-90 | -0.410384801 | 0.562 | 0.781 | 1.44E-86 |
| Mrpl23 | 2.32E-90 | -0.385260236 | 0.665 | 0.875 | 2.66E-86 |
| Gnb2 | 2.94E-90 | -0.412619394 | 0.723 | 0.922 | 3.36E-86 |
| Tbcc | 5.61E-90 | -0.404521497 | 0.76 | 0.86 | 6.42E-86 |
| Alcam | 6.86E-90 | 0.340279345 | 0.988 | 0.995 | 7.85E-86 |
| Tmem167 | 9.17E-90 | -0.423895804 | 0.779 | 0.952 | 1.05E-85 |
| Ccdc107 | 1.77E-89 | -0.40456586 | 0.691 | 0.902 | 2.02E-85 |
| Rps26 | 1.84E-89 | 0.448197393 | 0.954 | 0.953 | 2.11E-85 |
| Aimp2 | 2.09E-89 | -0.353405316 | 0.545 | 0.757 | 2.39E-85 |
| Mrpl46 | 2.12E-89 | -0.363899738 | 0.583 | 0.806 | 2.43E-85 |
| Ngfrap1 | 5.60E-89 | -0.348863963 | 0.89 | 0.975 | 6.41E-85 |
| Bub3 | 2.24E-88 | -0.290340004 | 0.594 | 0.809 | 2.56E-84 |
| Vbp1 | 3.93E-88 | -0.395635943 | 0.755 | 0.952 | 4.49E-84 |
| Scp2 | 5.46E-88 | -0.39913236 | 0.658 | 0.876 | 6.25E-84 |
| Mrpl11 | 5.91E-88 | -0.360174097 | 0.576 | 0.789 | 6.76E-84 |
| Cetn2 | 9.30E-88 | -0.366751352 | 0.417 | 0.733 | 1.06E-83 |
| Rgs4 | 4.08E-87 | 0.294161466 | 0.997 | 0.998 | 4.66E-83 |
| Rer1 | 4.89E-87 | -0.354339584 | 0.627 | 0.85 | 5.60E-83 |
| Psmg3 | 5.97E-87 | -0.357715281 | 0.529 | 0.732 | 6.83E-83 |
| Med19 | 7.77E-87 | -0.367977368 | 0.56 | 0.766 | 8.89E-83 |
| Tceal8 | 8.07E-87 | -0.409366553 | 0.763 | 0.943 | 9.23E-83 |
| Snrpn | 1.03E-86 | -0.30735143 | 0.967 | 0.993 | 1.17E-82 |
| Capns1 | 1.45E-86 | -0.362108063 | 0.903 | 0.985 | 1.66E-82 |
| Polr2i | 1.65E-86 | -0.352199108 | 0.649 | 0.861 | 1.88E-82 |
| Wfdc2 | 1.75E-86 | -0.491801981 | 0.747 | 0.918 | 2.00E-82 |
| Ptrhd1 | 1.99E-86 | -0.379193878 | 0.563 | 0.767 | 2.28E-82 |
| Sub1 | 3.48E-86 | -0.36283342 | 0.885 | 0.982 | 3.98E-82 |
| Spink2 | 4.08E-86 | -0.404109299 | 0.615 | 0.826 | 4.67E-82 |
| Tmem147 | 4.54E-86 | -0.394663398 | 0.717 | 0.915 | 5.20E-82 |
| Msmo1 | 7.11E-86 | -0.374377194 | 0.654 | 0.921 | 8.14E-82 |
| Clic1 | 1.07E-85 | -0.40444108 | 0.705 | 0.915 | 1.23E-81 |
| Tmem9 | 1.48E-85 | -0.300121444 | 0.569 | 0.767 | 1.70E-81 |
| Pdia3 | 2.08E-85 | -0.342881303 | 0.651 | 0.881 | 2.38E-81 |
| Ostc | 2.12E-85 | -0.374283953 | 0.659 | 0.872 | 2.42E-81 |
| 2610001J05Rik | 2.24E-85 | -0.388424576 | 0.731 | 0.926 | 2.56E-81 |
| Ssr2 | 2.65E-85 | -0.378524251 | 0.556 | 0.763 | 3.03E-81 |
| Snap47 | 6.03E-85 | -0.346521129 | 0.615 | 0.815 | 6.89E-81 |
| Atraid | 7.73E-85 | -0.331827889 | 0.539 | 0.748 | 8.84E-81 |
| Dpm3 | 9.44E-85 | -0.365375924 | 0.423 | 0.714 | 1.08E-80 |
| Ebp | 1.18E-84 | -0.416595727 | 0.576 | 0.79 | 1.36E-80 |
| Sdf2 | 1.43E-84 | -0.341988775 | 0.602 | 0.827 | 1.64E-80 |
| Malat1 | 1.79E-84 | 0.676149229 | 0.995 | 0.993 | 2.04E-80 |
| Mrpl14 | 2.40E-84 | -0.382821655 | 0.698 | 0.904 | 2.75E-80 |
| Tmco1 | 2.41E-84 | -0.384063575 | 0.61 | 0.823 | 2.76E-80 |
| Isca2 | 9.31E-84 | -0.36260543 | 0.627 | 0.831 | 1.07E-79 |
| Dctn6 | 1.42E-83 | -0.326712827 | 0.615 | 0.82 | 1.62E-79 |
| Chchd7 | 1.69E-83 | -0.26645288 | 0.405 | 0.689 | 1.93E-79 |
| Pfdn1 | 3.95E-83 | -0.373419162 | 0.758 | 0.944 | 4.52E-79 |
| Cd59a | 4.77E-83 | -0.396533478 | 0.767 | 0.949 | 5.46E-79 |
| Jkamp | 9.74E-83 | -0.386302478 | 0.651 | 0.876 | 1.11E-78 |
| Sod2 | 1.33E-82 | -0.364351852 | 0.672 | 0.889 | 1.52E-78 |
| Yif1a | 3.78E-82 | -0.388886915 | 0.557 | 0.773 | 4.32E-78 |
| Pet100 | 4.17E-82 | -0.290464902 | 0.456 | 0.739 | 4.77E-78 |
| Bad | 4.61E-82 | -0.369012654 | 0.676 | 0.877 | 5.28E-78 |
| Rogdi | 5.77E-82 | -0.298286958 | 0.553 | 0.739 | 6.61E-78 |
| Ccdc28a | 8.27E-82 | -0.332241702 | 0.568 | 0.759 | 9.46E-78 |
| Polr2h | 9.86E-82 | -0.276630648 | 0.56 | 0.751 | 1.13E-77 |
| Yif1b | 2.86E-81 | -0.370764879 | 0.557 | 0.773 | 3.27E-77 |
| Slc25a14 | 6.99E-81 | -0.330472819 | 0.553 | 0.754 | 7.99E-77 |
| Sumo1 | 1.18E-80 | -0.371615947 | 0.898 | 0.99 | 1.35E-76 |
| Elavl4 | 1.20E-80 | 0.382489969 | 0.976 | 0.984 | 1.37E-76 |
| Ppp1ca | 1.74E-80 | -0.385886483 | 0.798 | 0.953 | 2.00E-76 |
| Banf1 | 1.99E-80 | -0.303438996 | 0.472 | 0.773 | 2.28E-76 |
| Ywhab | 2.29E-80 | 0.408947098 | 0.954 | 0.975 | 2.62E-76 |
| Mrpl15 | 2.31E-80 | -0.358048829 | 0.541 | 0.737 | 2.65E-76 |
| Sdhaf4 | 3.82E-80 | -0.373214615 | 0.599 | 0.817 | 4.36E-76 |
| Th | 4.92E-80 | -0.479594119 | 0.792 | 0.921 | 5.63E-76 |
| Tsen15 | 5.01E-80 | -0.325172761 | 0.584 | 0.796 | 5.73E-76 |
| Sec61b | 5.25E-80 | -0.377961861 | 0.705 | 0.911 | 6.00E-76 |
| Smim14 | 8.16E-80 | -0.362410023 | 0.873 | 0.984 | 9.34E-76 |
| Sfr1 | 1.21E-79 | -0.325503274 | 0.624 | 0.835 | 1.38E-75 |
| Mrpl4 | 2.26E-79 | -0.303973547 | 0.552 | 0.756 | 2.59E-75 |
| Bnip3 | 2.32E-79 | -0.345751333 | 0.672 | 0.876 | 2.65E-75 |
| Mrps16 | 2.55E-79 | -0.302733979 | 0.612 | 0.806 | 2.92E-75 |
| Mrpl17 | 2.99E-79 | -0.328877959 | 0.641 | 0.842 | 3.42E-75 |
| Uqcc2 | 3.60E-79 | -0.371482341 | 0.803 | 0.97 | 4.11E-75 |
| Degs1 | 4.18E-79 | -0.388078756 | 0.73 | 0.921 | 4.78E-75 |
| Znhit1 | 5.61E-79 | -0.343707479 | 0.728 | 0.925 | 6.42E-75 |
| Commd7 | 6.52E-79 | -0.365831011 | 0.559 | 0.775 | 7.46E-75 |
| Trappc6a | 6.63E-79 | -0.318432199 | 0.6 | 0.799 | 7.58E-75 |
| Rplp1 | 7.10E-79 | 0.260244156 | 0.997 | 0.999 | 8.12E-75 |
| Pdcd6 | 1.04E-78 | -0.336535753 | 0.575 | 0.773 | 1.19E-74 |
| Arf5 | 1.55E-78 | -0.308396933 | 0.943 | 0.993 | 1.77E-74 |
| Pfdn4 | 1.98E-78 | -0.387572412 | 0.576 | 0.858 | 2.27E-74 |
| Smarcb1 | 2.20E-78 | -0.294793307 | 0.55 | 0.748 | 2.52E-74 |
| Gnb1 | 2.27E-78 | 0.61815315 | 0.968 | 0.965 | 2.60E-74 |
| Msrb1 | 2.44E-78 | -0.322955396 | 0.602 | 0.807 | 2.79E-74 |
| Atp6v0e | 2.56E-78 | -0.357125991 | 0.455 | 0.763 | 2.93E-74 |
| Izumo4 | 5.26E-78 | -0.319225934 | 0.547 | 0.74 | 6.02E-74 |
| Ahsa1 | 7.04E-78 | -0.298263974 | 0.62 | 0.835 | 8.05E-74 |
| Ceacam10 | 9.27E-78 | -0.959150263 | 0.754 | 0.833 | 1.06E-73 |
| 2-Mar | 1.02E-77 | -0.332590144 | 0.608 | 0.808 | 1.17E-73 |
| Pmvk | 1.10E-77 | -0.368465652 | 0.797 | 0.959 | 1.25E-73 |
| Mrps15 | 1.26E-77 | -0.320876107 | 0.621 | 0.821 | 1.45E-73 |
| Prune2 | 1.59E-77 | 0.667201327 | 0.993 | 0.99 | 1.82E-73 |
| Paip2 | 2.46E-77 | -0.366743919 | 0.806 | 0.965 | 2.82E-73 |
| Ccdc167 | 2.54E-77 | -0.346786162 | 0.594 | 0.791 | 2.91E-73 |
| Pdia6 | 3.63E-77 | -0.270116409 | 0.747 | 0.833 | 4.15E-73 |
| Uqcrc2 | 6.27E-77 | -0.308090845 | 0.592 | 0.801 | 7.17E-73 |
| Mzt2 | 6.85E-77 | -0.340202282 | 0.566 | 0.767 | 7.84E-73 |
| Rps9 | 1.00E-76 | 0.252063384 | 0.995 | 0.999 | 1.14E-72 |
| H3f3b | 1.37E-76 | -0.365825917 | 0.86 | 0.981 | 1.57E-72 |
| Haghl | 1.39E-76 | -0.251887912 | 0.549 | 0.724 | 1.59E-72 |
| Clu | 1.76E-76 | -0.259132246 | 0.335 | 0.587 | 2.01E-72 |
| Rpl21 | 2.00E-76 | 0.404141247 | 0.959 | 0.968 | 2.29E-72 |
| Cops5 | 2.75E-76 | -0.291881069 | 0.6 | 0.787 | 3.15E-72 |
| C1d | 3.31E-76 | -0.369771605 | 0.848 | 0.982 | 3.79E-72 |
| Rps12 | 6.61E-76 | 0.486969017 | 0.909 | 0.902 | 7.56E-72 |
| Pam16 | 8.15E-76 | -0.339733605 | 0.707 | 0.893 | 9.32E-72 |
| Trappc2 | 1.00E-75 | -0.375912717 | 0.719 | 0.916 | 1.15E-71 |
| Mrps36 | 1.37E-75 | -0.330565413 | 0.685 | 0.892 | 1.57E-71 |
| Amn1 | 1.82E-75 | -0.337818612 | 0.549 | 0.741 | 2.08E-71 |
| Esd | 2.11E-75 | -0.303231303 | 0.579 | 0.778 | 2.42E-71 |
| Polr2l | 5.72E-75 | -0.256011153 | 0.454 | 0.725 | 6.54E-71 |
| Exosc7 | 7.81E-75 | -0.32885325 | 0.538 | 0.729 | 8.93E-71 |
| Mvk | 7.92E-75 | -0.26390333 | 0.542 | 0.713 | 9.07E-71 |
| Mtpn | 1.13E-74 | 0.562695711 | 0.854 | 0.851 | 1.29E-70 |
| Immp1l | 2.16E-74 | -0.298808187 | 0.585 | 0.791 | 2.48E-70 |
| Psmc6 | 3.65E-74 | -0.286683902 | 0.618 | 0.824 | 4.18E-70 |
| Emc10 | 5.22E-74 | -0.350096279 | 0.775 | 0.943 | 5.97E-70 |
| Dctn2 | 1.26E-73 | -0.319447061 | 0.697 | 0.896 | 1.44E-69 |
| 1110008P14Rik | 1.37E-73 | -0.346963324 | 0.847 | 0.974 | 1.57E-69 |
| Cdk5 | 1.42E-73 | -0.275158233 | 0.605 | 0.799 | 1.63E-69 |
| Itpa | 1.81E-73 | -0.263063336 | 0.54 | 0.712 | 2.07E-69 |
| Tomm40 | 1.82E-73 | -0.31670537 | 0.554 | 0.743 | 2.08E-69 |
| Slc35g2 | 2.89E-73 | -0.331094901 | 0.535 | 0.734 | 3.30E-69 |
| Lsm7 | 3.26E-73 | -0.276485841 | 0.413 | 0.688 | 3.73E-69 |
| Mrpl2 | 6.18E-73 | -0.320552787 | 0.542 | 0.731 | 7.08E-69 |
| Tmx2 | 8.17E-73 | -0.319834795 | 0.665 | 0.869 | 9.35E-69 |
| Lsm4 | 1.26E-72 | -0.341651155 | 0.686 | 0.88 | 1.44E-68 |
| Hes6 | 1.51E-72 | -0.340885092 | 0.576 | 0.783 | 1.73E-68 |
| 2210013O21Rik | 1.60E-72 | -0.365095097 | 0.805 | 0.954 | 1.83E-68 |
| Gsk3b | 1.72E-72 | 0.486328375 | 0.939 | 0.943 | 1.97E-68 |
| Cisd3 | 1.75E-72 | -0.346008901 | 0.788 | 0.95 | 2.01E-68 |
| Srp19 | 2.89E-72 | -0.303875241 | 0.69 | 0.891 | 3.31E-68 |
| Hint2 | 3.15E-72 | -0.370576274 | 0.683 | 0.883 | 3.60E-68 |
| Lamtor2 | 4.03E-72 | -0.348804287 | 0.814 | 0.968 | 4.62E-68 |
| 0610007P14Rik | 6.21E-72 | -0.270220719 | 0.757 | 0.828 | 7.10E-68 |
| Cirbp | 6.67E-72 | -0.337878673 | 0.733 | 0.921 | 7.64E-68 |
| Commd6 | 7.13E-72 | -0.353588226 | 0.571 | 0.772 | 8.16E-68 |
| Mrps18c | 8.68E-72 | -0.342953909 | 0.669 | 0.87 | 9.94E-68 |
| Rpl10a | 1.20E-71 | 0.370449813 | 0.964 | 0.958 | 1.37E-67 |
| Rbm42 | 1.46E-71 | -0.308898395 | 0.548 | 0.737 | 1.67E-67 |
| Rps19bp1 | 3.49E-71 | -0.272255293 | 0.4 | 0.668 | 4.00E-67 |
| Mrpl43 | 6.55E-71 | -0.292667137 | 0.633 | 0.828 | 7.50E-67 |
| Ubl7 | 1.04E-70 | -0.305198549 | 0.547 | 0.731 | 1.19E-66 |
| Rpl22l1 | 1.04E-70 | 0.33489919 | 0.991 | 0.996 | 1.19E-66 |
| Commd9 | 1.26E-70 | -0.345649923 | 0.542 | 0.739 | 1.44E-66 |
| Cltc | 1.54E-70 | 0.460376285 | 0.908 | 0.922 | 1.76E-66 |
| Lhfpl5 | 2.04E-70 | -0.438608884 | 0.535 | 0.753 | 2.33E-66 |
| Pfdn5 | 2.55E-70 | 0.375975286 | 0.956 | 0.963 | 2.91E-66 |
| Exosc5 | 2.64E-70 | -0.310145242 | 0.523 | 0.706 | 3.02E-66 |
| Gtf2a2 | 3.54E-70 | -0.325052436 | 0.647 | 0.841 | 4.04E-66 |
| Ndfip1 | 3.91E-70 | -0.378193246 | 0.531 | 0.73 | 4.48E-66 |
| Hopx | 4.30E-70 | -0.382741298 | 0.78 | 0.933 | 4.92E-66 |
| Ercc1 | 4.42E-70 | -0.298639163 | 0.526 | 0.712 | 5.05E-66 |
| Wbp5 | 5.58E-70 | -0.349104407 | 0.858 | 0.955 | 6.38E-66 |
| Tomm5 | 6.13E-70 | -0.349615395 | 0.791 | 0.951 | 7.01E-66 |
| Rps24 | 8.33E-70 | 0.434296269 | 0.921 | 0.924 | 9.54E-66 |
| 1110051M20Rik | 8.34E-70 | -0.250250271 | 0.549 | 0.709 | 9.54E-66 |
| Ywhaq | 1.25E-69 | -0.310589789 | 0.94 | 0.99 | 1.43E-65 |
| Mrps18a | 1.57E-69 | -0.316502443 | 0.698 | 0.884 | 1.80E-65 |
| Creb3 | 2.39E-69 | -0.295199922 | 0.534 | 0.716 | 2.74E-65 |
| Pafah1b3 | 3.91E-69 | -0.339644652 | 0.627 | 0.835 | 4.48E-65 |
| Pigyl | 4.19E-69 | -0.261217018 | 0.609 | 0.796 | 4.79E-65 |
| Snx15 | 4.46E-69 | -0.277776668 | 0.378 | 0.657 | 5.10E-65 |
| Ptov1 | 4.88E-69 | -0.316111875 | 0.716 | 0.898 | 5.58E-65 |
| Ccni | 4.92E-69 | 0.515307863 | 0.87 | 0.873 | 5.63E-65 |
| Nsmce1 | 1.72E-68 | -0.316031597 | 0.543 | 0.721 | 1.97E-64 |
| Ly6h | 3.06E-68 | -0.274379665 | 0.741 | 0.924 | 3.51E-64 |
| Rps3 | 3.28E-68 | 0.340006574 | 0.983 | 0.985 | 3.75E-64 |
| Slc35b1 | 3.63E-68 | -0.291313429 | 0.576 | 0.766 | 4.15E-64 |
| B2m | 3.72E-68 | -0.317713169 | 0.76 | 0.854 | 4.26E-64 |
| Rps28 | 4.61E-68 | 0.394799778 | 0.97 | 0.969 | 5.28E-64 |
| Mrpl22 | 5.34E-68 | -0.418771594 | 0.725 | 0.825 | 6.11E-64 |
| Sbds | 5.75E-68 | -0.295515468 | 0.598 | 0.807 | 6.58E-64 |
| Atp1b3 | 9.31E-68 | 0.297601607 | 0.988 | 0.995 | 1.06E-63 |
| Psmd12 | 1.28E-67 | -0.299301128 | 0.625 | 0.806 | 1.46E-63 |
| Vdac2 | 1.54E-67 | -0.28711757 | 0.679 | 0.883 | 1.77E-63 |
| Mrpl32 | 1.71E-67 | -0.263857992 | 0.557 | 0.746 | 1.96E-63 |
| Fam162a | 2.14E-67 | -0.278732912 | 0.619 | 0.818 | 2.45E-63 |
| Atxn7l3b | 2.15E-67 | -0.332154094 | 0.844 | 0.963 | 2.46E-63 |
| Etfa | 2.40E-67 | -0.32589987 | 0.631 | 0.824 | 2.74E-63 |
| Mrpl57 | 2.66E-67 | -0.325605912 | 0.694 | 0.88 | 3.04E-63 |
| Cct8 | 4.25E-67 | -0.274861566 | 0.663 | 0.868 | 4.86E-63 |
| Commd2 | 7.44E-67 | -0.30935997 | 0.54 | 0.722 | 8.51E-63 |
| Pop7 | 7.66E-67 | -0.281085757 | 0.565 | 0.751 | 8.76E-63 |
| Pja2 | 9.25E-67 | 0.426815653 | 0.917 | 0.933 | 1.06E-62 |
| Stoml1 | 1.28E-66 | -0.303247381 | 0.583 | 0.791 | 1.46E-62 |
| Sptbn1 | 1.88E-66 | 0.423438666 | 0.915 | 0.922 | 2.15E-62 |
| Ddost | 2.85E-66 | -0.276041976 | 0.56 | 0.746 | 3.26E-62 |
| Mkks | 3.86E-66 | -0.256930097 | 0.615 | 0.802 | 4.41E-62 |
| Cct3 | 6.21E-66 | -0.255712393 | 0.612 | 0.815 | 7.11E-62 |
| Them6 | 6.54E-66 | -0.282353534 | 0.67 | 0.863 | 7.48E-62 |
| Emd | 6.68E-66 | -0.357444421 | 0.55 | 0.744 | 7.64E-62 |
| Ost4 | 7.80E-66 | -0.333317432 | 0.769 | 0.944 | 8.92E-62 |
| Sec11c | 7.90E-66 | -0.252191301 | 0.634 | 0.818 | 9.04E-62 |
| Exosc4 | 9.06E-66 | -0.302633509 | 0.54 | 0.722 | 1.04E-61 |
| Wdr18 | 9.67E-66 | -0.33687153 | 0.535 | 0.73 | 1.11E-61 |
| Tmed9 | 1.40E-65 | -0.28134922 | 0.619 | 0.815 | 1.60E-61 |
| 2410015M20Rik | 2.13E-65 | -0.302087143 | 0.708 | 0.898 | 2.44E-61 |
| Cops3 | 2.13E-65 | -0.292238323 | 0.537 | 0.719 | 2.44E-61 |
| Ndn | 2.35E-65 | -0.267477854 | 0.585 | 0.758 | 2.69E-61 |
| Snhg11 | 2.79E-65 | 0.463806712 | 0.929 | 0.932 | 3.20E-61 |
| Rpl23a | 3.23E-65 | 0.358113215 | 0.957 | 0.967 | 3.69E-61 |
| Mrps11 | 3.39E-65 | -0.325700307 | 0.573 | 0.767 | 3.88E-61 |
| Wdr61 | 4.07E-65 | -0.264925732 | 0.54 | 0.721 | 4.66E-61 |
| Psmd13 | 5.64E-65 | -0.269946468 | 0.572 | 0.755 | 6.45E-61 |
| Txndc15 | 5.74E-65 | -0.272870778 | 0.585 | 0.775 | 6.57E-61 |
| Prmt2 | 6.00E-65 | -0.252600902 | 0.646 | 0.836 | 6.86E-61 |
| Tmed10 | 8.86E-65 | -0.293274349 | 0.682 | 0.875 | 1.01E-60 |
| Raly | 8.89E-65 | -0.265798285 | 0.64 | 0.83 | 1.02E-60 |
| Ndufaf3 | 8.94E-65 | -0.254199432 | 0.62 | 0.819 | 1.02E-60 |
| Prkar1a | 1.04E-64 | 0.311317627 | 0.99 | 0.994 | 1.19E-60 |
| Nrip3 | 1.06E-64 | 0.428102682 | 0.889 | 0.9 | 1.22E-60 |
| Pepd | 1.21E-64 | -0.278699493 | 0.535 | 0.71 | 1.38E-60 |
| Vapa | 1.24E-64 | -0.308948447 | 0.783 | 0.957 | 1.42E-60 |
| Mrpl18 | 1.29E-64 | -0.266072303 | 0.65 | 0.845 | 1.47E-60 |
| Rit1 | 1.36E-64 | -0.278114323 | 0.591 | 0.785 | 1.56E-60 |
| Tmem55b | 1.55E-64 | -0.284578002 | 0.709 | 0.903 | 1.78E-60 |
| Hax1 | 1.58E-64 | -0.281684493 | 0.577 | 0.758 | 1.81E-60 |
| Prdx4 | 1.70E-64 | -0.284343346 | 0.541 | 0.72 | 1.94E-60 |
| Cnih4 | 2.43E-64 | -0.273178438 | 0.56 | 0.739 | 2.78E-60 |
| Ccl27a | 2.50E-64 | -0.274885585 | 0.47 | 0.737 | 2.86E-60 |
| Tsn | 2.76E-64 | -0.300209744 | 0.707 | 0.887 | 3.16E-60 |
| Nudt16l1 | 5.66E-64 | -0.277422292 | 0.751 | 0.811 | 6.47E-60 |
| Aasdhppt | 5.70E-64 | -0.264924096 | 0.572 | 0.765 | 6.52E-60 |
| Adprh | 6.56E-64 | -0.271768225 | 0.564 | 0.845 | 7.51E-60 |
| Atp6ap2 | 8.24E-64 | -0.325415357 | 0.764 | 0.939 | 9.42E-60 |
| Ppp1r35 | 1.16E-63 | -0.287190594 | 0.63 | 0.833 | 1.33E-59 |
| Cd151 | 1.17E-63 | -0.314839567 | 0.74 | 0.919 | 1.34E-59 |
| Ssr4 | 1.27E-63 | -0.270749038 | 0.658 | 0.845 | 1.45E-59 |
| Spr | 1.34E-63 | -0.319884335 | 0.545 | 0.739 | 1.53E-59 |
| Eif5a | 1.40E-63 | -0.330193277 | 0.878 | 0.975 | 1.60E-59 |
| Gsto1 | 2.36E-63 | -0.27757859 | 0.549 | 0.729 | 2.70E-59 |
| Slc25a5 | 2.60E-63 | -0.324597587 | 0.856 | 0.969 | 2.98E-59 |
| Wdr83os | 2.73E-63 | -0.304104349 | 0.735 | 0.919 | 3.12E-59 |
| Eri3 | 3.76E-63 | -0.256667646 | 0.613 | 0.808 | 4.30E-59 |
| Tmem45b | 4.81E-63 | -0.448123475 | 0.69 | 0.843 | 5.50E-59 |
| Psmc3 | 7.23E-63 | -0.280324044 | 0.682 | 0.869 | 8.27E-59 |
| Txnl4a | 8.29E-63 | -0.277402558 | 0.669 | 0.847 | 9.49E-59 |
| 1700021F05Rik | 1.29E-62 | -0.268097794 | 0.549 | 0.725 | 1.48E-58 |
| Glo1 | 1.75E-62 | -0.260204515 | 0.632 | 0.814 | 2.00E-58 |
| Serpinf1 | 2.48E-62 | -0.37177246 | 0.535 | 0.749 | 2.84E-58 |
| Cetn3 | 2.63E-62 | -0.284012356 | 0.664 | 0.86 | 3.01E-58 |
| Rbm8a | 2.90E-62 | -0.270596978 | 0.682 | 0.874 | 3.31E-58 |
| Gde1 | 3.27E-62 | -0.303068281 | 0.65 | 0.858 | 3.74E-58 |
| Tpd52 | 4.01E-62 | 0.360646545 | 0.965 | 0.975 | 4.59E-58 |
| Srm | 4.44E-62 | -0.281987704 | 0.53 | 0.702 | 5.08E-58 |
| Uqcrfs1 | 4.44E-62 | -0.312719039 | 0.765 | 0.937 | 5.08E-58 |
| Fam103a1 | 6.58E-62 | -0.319882373 | 0.804 | 0.954 | 7.53E-58 |
| Ctsz | 1.07E-61 | -0.25637262 | 0.563 | 0.744 | 1.23E-57 |
| Trappc5 | 1.12E-61 | -0.287386126 | 0.542 | 0.717 | 1.28E-57 |
| Psme1 | 1.21E-61 | -0.279540423 | 0.568 | 0.74 | 1.38E-57 |
| Mrps18b | 1.23E-61 | -0.262562813 | 0.545 | 0.723 | 1.40E-57 |
| Ssbp1 | 1.27E-61 | -0.306421953 | 0.541 | 0.738 | 1.45E-57 |
| Blvra | 1.29E-61 | -0.297487018 | 0.55 | 0.743 | 1.47E-57 |
| 0610012G03Rik | 1.39E-61 | -0.30459228 | 0.64 | 0.886 | 1.59E-57 |
| Slc50a1 | 2.00E-61 | -0.262494445 | 0.551 | 0.731 | 2.29E-57 |
| Mrpl54 | 2.57E-61 | -0.308555004 | 0.767 | 0.934 | 2.94E-57 |
| Lsm5 | 2.69E-61 | -0.28762109 | 0.495 | 0.784 | 3.07E-57 |
| Ndrg4 | 3.41E-61 | 0.336711379 | 0.966 | 0.976 | 3.90E-57 |
| Pts | 4.56E-61 | -0.276857598 | 0.626 | 0.807 | 5.22E-57 |
| Fcf1 | 5.20E-61 | -0.313675534 | 0.536 | 0.722 | 5.95E-57 |
| Eny2 | 5.38E-61 | -0.284328674 | 0.661 | 0.861 | 6.15E-57 |
| Ndufb10 | 8.55E-61 | -0.303924688 | 0.775 | 0.954 | 9.78E-57 |
| Iah1 | 1.16E-60 | -0.279000465 | 0.535 | 0.703 | 1.33E-56 |
| Ncoa7 | 3.88E-60 | 0.35005949 | 0.954 | 0.976 | 4.44E-56 |
| Eif4a1 | 5.34E-60 | -0.320503664 | 0.78 | 0.934 | 6.11E-56 |
| Tsg101 | 6.94E-60 | -0.266300436 | 0.693 | 0.884 | 7.94E-56 |
| Mrpl40 | 1.00E-59 | -0.27441445 | 0.551 | 0.723 | 1.15E-55 |
| Tmem199 | 1.14E-59 | -0.292524034 | 0.548 | 0.739 | 1.30E-55 |
| Chrac1 | 2.09E-59 | -0.272595021 | 0.56 | 0.731 | 2.39E-55 |
| Aig1 | 3.64E-59 | -0.292037033 | 0.679 | 0.858 | 4.17E-55 |
| Mtx1 | 4.67E-59 | -0.275320584 | 0.524 | 0.695 | 5.35E-55 |
| Stx8 | 5.31E-59 | -0.267225431 | 0.532 | 0.7 | 6.08E-55 |
| Ogfrl1 | 5.73E-59 | 0.448026773 | 0.867 | 0.88 | 6.56E-55 |
| Krtcap2 | 5.90E-59 | -0.298935556 | 0.732 | 0.912 | 6.75E-55 |
| Camk2d | 6.21E-59 | 0.384386579 | 0.919 | 0.95 | 7.10E-55 |
| Nxf1 | 7.51E-59 | -0.289279106 | 0.621 | 0.801 | 8.59E-55 |
| Tmbim6 | 9.94E-59 | -0.256346372 | 0.564 | 0.763 | 1.14E-54 |
| Pigp | 1.07E-58 | -0.26308787 | 0.682 | 0.885 | 1.23E-54 |
| Arpc1a | 1.35E-58 | 0.40211453 | 0.945 | 0.952 | 1.54E-54 |
| Ube2w | 1.73E-58 | -0.254088572 | 0.692 | 0.896 | 1.98E-54 |
| Tsfm | 1.82E-58 | -0.261569332 | 0.549 | 0.726 | 2.08E-54 |
| Marcks | 2.54E-58 | 0.38871341 | 0.919 | 0.936 | 2.91E-54 |
| Mrps28 | 3.25E-58 | -0.313615078 | 0.531 | 0.708 | 3.72E-54 |
| Timm10b | 3.67E-58 | -0.266911278 | 0.661 | 0.861 | 4.20E-54 |
| Tmem261 | 8.59E-58 | -0.302196564 | 0.532 | 0.711 | 9.83E-54 |
| Ndufa3 | 1.17E-57 | -0.312508096 | 0.693 | 0.913 | 1.34E-53 |
| Grina | 1.23E-57 | -0.295399737 | 0.751 | 0.94 | 1.41E-53 |
| Alkbh6 | 1.39E-57 | -0.27212967 | 0.557 | 0.725 | 1.59E-53 |
| Bax | 2.10E-57 | -0.274525032 | 0.692 | 0.883 | 2.40E-53 |
| Dtd1 | 4.20E-57 | -0.259612136 | 0.533 | 0.72 | 4.81E-53 |
| Dnaja1 | 5.85E-57 | -0.294332076 | 0.866 | 0.977 | 6.69E-53 |
| Efcab10 | 6.65E-57 | -0.300987236 | 0.531 | 0.714 | 7.61E-53 |
| Hnrnpa1 | 7.80E-57 | 0.414131322 | 0.893 | 0.921 | 8.93E-53 |
| Tmem158 | 9.70E-57 | -0.302117758 | 0.572 | 0.782 | 1.11E-52 |
| Rpl4 | 1.17E-56 | 0.393730916 | 0.964 | 0.961 | 1.34E-52 |
| Rsrp1 | 1.60E-56 | -0.276422844 | 0.583 | 0.851 | 1.83E-52 |
| Fbxo2 | 2.08E-56 | -0.300975544 | 0.789 | 0.935 | 2.39E-52 |
| Mydgf | 2.39E-56 | -0.282548741 | 0.554 | 0.738 | 2.73E-52 |
| Atp5c1 | 2.42E-56 | -0.285628728 | 0.8 | 0.955 | 2.77E-52 |
| Fdft1 | 2.60E-56 | -0.275584785 | 0.581 | 0.839 | 2.97E-52 |
| Pin4 | 2.68E-56 | -0.259560006 | 0.589 | 0.847 | 3.06E-52 |
| Psmg2 | 3.14E-56 | -0.25912703 | 0.517 | 0.685 | 3.59E-52 |
| Fam96b | 4.24E-56 | -0.28030484 | 0.748 | 0.921 | 4.85E-52 |
| Polr2c | 5.36E-56 | -0.270446064 | 0.546 | 0.733 | 6.13E-52 |
| Glrx3 | 5.81E-56 | -0.289812949 | 0.708 | 0.899 | 6.65E-52 |
| Rasgrp1 | 1.15E-55 | 0.466010333 | 0.902 | 0.885 | 1.31E-51 |
| Snrnp27 | 1.19E-55 | -0.277786962 | 0.63 | 0.881 | 1.37E-51 |
| Mpv17l2 | 1.53E-55 | -0.280393658 | 0.744 | 0.805 | 1.75E-51 |
| Rpsa | 1.79E-55 | 0.405739019 | 0.909 | 0.914 | 2.04E-51 |
| Cox20 | 1.81E-55 | -0.261109394 | 0.509 | 0.78 | 2.07E-51 |
| Eif5 | 2.17E-55 | -0.308503139 | 0.805 | 0.952 | 2.48E-51 |
| Ppp3ca | 2.80E-55 | 0.374914839 | 0.943 | 0.96 | 3.21E-51 |
| Rps21 | 1.23E-54 | 0.331287263 | 0.974 | 0.978 | 1.41E-50 |
| Kif1b | 1.45E-54 | 0.352285189 | 0.93 | 0.943 | 1.66E-50 |
| Rpl19 | 1.64E-54 | 0.272662513 | 0.983 | 0.986 | 1.88E-50 |
| Tusc5 | 1.71E-54 | 0.480279242 | 0.881 | 0.896 | 1.95E-50 |
| Pura | 2.39E-54 | 0.345095393 | 0.945 | 0.962 | 2.74E-50 |
| Ufc1 | 5.35E-54 | -0.267207819 | 0.679 | 0.852 | 6.12E-50 |
| Gm4950 | 1.75E-53 | -0.250300631 | 0.517 | 0.675 | 2.01E-49 |
| Nrp1 | 2.09E-53 | 0.394767439 | 0.891 | 0.913 | 2.39E-49 |
| Klhdc2 | 3.87E-53 | -0.265589046 | 0.677 | 0.863 | 4.42E-49 |
| Actr6 | 3.91E-53 | -0.270710848 | 0.529 | 0.704 | 4.47E-49 |
| 1110004F10Rik | 5.91E-53 | -0.26887617 | 0.611 | 0.862 | 6.76E-49 |
| Vps8 | 1.01E-52 | 0.600626235 | 0.84 | 0.843 | 1.16E-48 |
| Asph | 1.20E-52 | -0.258306254 | 0.655 | 0.816 | 1.38E-48 |
| P2ry1 | 1.53E-52 | 0.296546805 | 0.983 | 0.988 | 1.75E-48 |
| Urod | 1.89E-52 | -0.271238856 | 0.523 | 0.698 | 2.16E-48 |
| Dst | 1.99E-52 | 0.410843519 | 0.864 | 0.88 | 2.28E-48 |
| Eif2s2 | 3.77E-52 | -0.286642377 | 0.665 | 0.898 | 4.31E-48 |
| Pfdn2 | 3.97E-52 | 0.34012697 | 0.945 | 0.961 | 4.54E-48 |
| Iscu | 6.77E-52 | -0.279639541 | 0.841 | 0.962 | 7.74E-48 |
| Ssbp3 | 2.56E-51 | 0.385986906 | 0.911 | 0.938 | 2.93E-47 |
| Rab6a | 4.03E-51 | 0.258256237 | 0.989 | 0.995 | 4.61E-47 |
| Cops6 | 4.31E-51 | -0.261342042 | 0.67 | 0.862 | 4.94E-47 |
| Nrsn1 | 4.59E-51 | 0.432933542 | 0.918 | 0.925 | 5.25E-47 |
| Gm1673 | 7.17E-51 | -0.279941642 | 0.849 | 0.963 | 8.21E-47 |
| Snrpd2 | 7.87E-51 | -0.275605741 | 0.803 | 0.936 | 9.00E-47 |
| Eif2s1 | 1.02E-50 | -0.277287344 | 0.723 | 0.769 | 1.17E-46 |
| Sv2c | 1.29E-50 | 0.422794948 | 0.888 | 0.924 | 1.48E-46 |
| Saysd1 | 3.05E-50 | -0.287057762 | 0.728 | 0.767 | 3.49E-46 |
| Rbms1 | 6.20E-50 | 0.38678868 | 0.911 | 0.939 | 7.10E-46 |
| Prkar2a | 8.87E-50 | 0.473064377 | 0.823 | 0.836 | 1.02E-45 |
| Syt1 | 2.33E-49 | 0.340435375 | 0.925 | 0.949 | 2.66E-45 |
| Atg3 | 2.91E-49 | -0.270868383 | 0.656 | 0.845 | 3.32E-45 |
| Tlx3 | 3.61E-49 | -0.255094867 | 0.746 | 0.915 | 4.13E-45 |
| Ldb2 | 4.76E-49 | 0.387044242 | 0.929 | 0.928 | 5.45E-45 |
| Lyrm2 | 6.07E-49 | -0.257407629 | 0.555 | 0.723 | 6.95E-45 |
| Card19 | 1.19E-48 | -0.270560244 | 0.902 | 0.982 | 1.36E-44 |
| Gm13889 | 1.51E-48 | -0.277924447 | 0.713 | 0.91 | 1.73E-44 |
| Ube2n | 1.96E-48 | -0.263119407 | 0.772 | 0.939 | 2.25E-44 |
| Sec62 | 2.96E-48 | -0.27167587 | 0.827 | 0.964 | 3.39E-44 |
| Eef2 | 5.76E-48 | 0.46331961 | 0.864 | 0.913 | 6.58E-44 |
| Syt11 | 7.91E-48 | 0.328902003 | 0.935 | 0.97 | 9.05E-44 |
| Sumo2 | 1.01E-47 | -0.265157639 | 0.894 | 0.988 | 1.16E-43 |
| Med10 | 2.19E-47 | -0.296145941 | 0.609 | 0.7 | 2.50E-43 |
| BC005537 | 3.67E-47 | 0.35478918 | 0.915 | 0.946 | 4.20E-43 |
| Casz1 | 3.78E-47 | 0.394154223 | 0.85 | 0.878 | 4.32E-43 |
| Ccl1 | 4.38E-47 | -0.266369378 | 0.574 | 0.737 | 5.01E-43 |
| Cd24a | 8.74E-47 | 0.27959768 | 0.968 | 0.976 | 1.00E-42 |
| Fgf12 | 2.14E-46 | -0.251648222 | 0.791 | 0.938 | 2.45E-42 |
| Raph1 | 2.18E-46 | 0.45040365 | 0.83 | 0.841 | 2.49E-42 |
| Slc51a | 4.85E-46 | -0.287865603 | 0.514 | 0.674 | 5.54E-42 |
| Rpl7 | 6.98E-46 | 0.3423863 | 0.955 | 0.96 | 7.99E-42 |
| Rps25 | 8.16E-46 | 0.292012594 | 0.97 | 0.978 | 9.34E-42 |
| Pafah1b1 | 2.24E-45 | 0.357501777 | 0.936 | 0.951 | 2.56E-41 |
| Arpc5l | 3.44E-45 | -0.265019396 | 0.725 | 0.893 | 3.93E-41 |
| Ppp1r2 | 6.58E-45 | 0.264307092 | 0.98 | 0.989 | 7.53E-41 |
| Capzb | 7.12E-45 | -0.255205813 | 0.723 | 0.914 | 8.15E-41 |
| Fos | 9.79E-45 | 0.618058096 | 0.732 | 0.635 | 1.12E-40 |
| Stxbp1 | 1.28E-44 | 0.460062832 | 0.875 | 0.862 | 1.46E-40 |
| Rpl7a | 5.01E-44 | 0.442376095 | 0.892 | 0.894 | 5.73E-40 |
| Pkm | 5.12E-44 | -0.262724732 | 0.827 | 0.954 | 5.86E-40 |
| Rpl35 | 5.90E-44 | 0.315325054 | 0.944 | 0.962 | 6.75E-40 |
| Rpl39 | 1.58E-43 | 0.268931934 | 0.985 | 0.994 | 1.81E-39 |
| Hspe1 | 1.68E-43 | -0.251273758 | 0.79 | 0.948 | 1.93E-39 |
| Atp6v1a | 2.33E-43 | 0.348286592 | 0.92 | 0.955 | 2.67E-39 |
| Gnao1 | 6.36E-43 | 0.448549368 | 0.846 | 0.887 | 7.28E-39 |
| Arpc1b | 1.13E-42 | 0.356974107 | 0.956 | 0.972 | 1.29E-38 |
| Ptms | 2.28E-42 | 0.314134208 | 0.953 | 0.967 | 2.61E-38 |
| Ap1ar | 7.45E-42 | 0.409129643 | 0.832 | 0.845 | 8.52E-38 |
| Atp2b1 | 7.65E-42 | 0.29537869 | 0.936 | 0.967 | 8.75E-38 |
| Txndc17 | 8.50E-42 | -0.258785974 | 0.81 | 0.932 | 9.72E-38 |
| Rpl23 | 1.71E-40 | 0.250321969 | 0.981 | 0.99 | 1.95E-36 |
| Napb | 1.81E-40 | 0.380566738 | 0.855 | 0.877 | 2.07E-36 |
| Lifr | 2.15E-39 | 0.384083697 | 0.846 | 0.889 | 2.46E-35 |
| Zfpl1 | 7.48E-39 | -0.28927352 | 0.527 | 0.704 | 8.55E-35 |
| Dpp10 | 8.21E-39 | 0.351870869 | 0.88 | 0.907 | 9.40E-35 |
| Ddx3x | 1.70E-38 | 0.347812338 | 0.876 | 0.904 | 1.95E-34 |
| Slc17a6 | 1.90E-37 | 0.388724844 | 0.883 | 0.89 | 2.17E-33 |
| Spock2 | 2.68E-37 | 0.358139297 | 0.846 | 0.874 | 3.07E-33 |
| Ret | 5.46E-37 | 0.312046089 | 0.946 | 0.965 | 6.25E-33 |
| Rpl29 | 7.24E-37 | 0.309446039 | 0.955 | 0.96 | 8.28E-33 |
| Ahnak | 1.90E-36 | 0.359060544 | 0.894 | 0.934 | 2.18E-32 |
| Cd63 | 2.62E-36 | -0.252533748 | 0.75 | 0.894 | 3.00E-32 |
| Bag1 | 3.71E-36 | 0.327946077 | 0.938 | 0.962 | 4.25E-32 |
| Canx | 3.94E-36 | 0.370943249 | 0.875 | 0.907 | 4.51E-32 |
| Gfra2 | 9.01E-36 | 0.320429984 | 0.927 | 0.919 | 1.03E-31 |
| Rps15 | 1.48E-35 | 0.25876375 | 0.965 | 0.977 | 1.69E-31 |
| Cox7a2l | 9.29E-35 | 0.260684023 | 0.98 | 0.99 | 1.06E-30 |
| Runx1 | 2.36E-33 | 0.330229725 | 0.917 | 0.945 | 2.70E-29 |
| Pgm2l1 | 1.08E-32 | 0.339643657 | 0.859 | 0.916 | 1.24E-28 |
| Olfm1 | 2.86E-32 | 0.276879758 | 0.913 | 0.95 | 3.27E-28 |
| App | 2.89E-32 | 0.303204715 | 0.912 | 0.948 | 3.30E-28 |
| Prkce | 4.99E-32 | 0.41248948 | 0.794 | 0.824 | 5.71E-28 |
| Gnaq | 6.20E-31 | 0.439064956 | 0.76 | 0.79 | 7.09E-27 |
| Camta1 | 8.10E-31 | 0.419460101 | 0.85 | 0.849 | 9.27E-27 |
| Zfp521 | 1.10E-30 | 0.373338841 | 0.815 | 0.858 | 1.25E-26 |
| Magi1 | 6.78E-30 | 0.373753702 | 0.807 | 0.844 | 7.76E-26 |
| Slc9a6 | 2.39E-29 | 0.354780635 | 0.835 | 0.864 | 2.73E-25 |
| Cplx1 | 1.88E-27 | 0.342781775 | 0.862 | 0.916 | 2.15E-23 |
| Rnf11 | 8.35E-27 | 0.376897558 | 0.8 | 0.837 | 9.55E-23 |
| Plagl1 | 9.50E-27 | 0.404116029 | 0.781 | 0.782 | 1.09E-22 |
| Cab39 | 9.64E-27 | 0.404036006 | 0.742 | 0.766 | 1.10E-22 |
| Csde1 | 1.05E-26 | 0.381802953 | 0.81 | 0.859 | 1.20E-22 |
| Tgoln1 | 2.21E-26 | 0.337026602 | 0.811 | 0.847 | 2.53E-22 |
| Ank3 | 6.95E-26 | 0.405430779 | 0.83 | 0.833 | 7.95E-22 |
| Kcnd3 | 5.39E-25 | 0.30189813 | 0.881 | 0.916 | 6.17E-21 |
| Emp3 | 1.89E-24 | -0.31767357 | 0.74 | 0.755 | 2.16E-20 |
| Gabbr2 | 4.18E-24 | 0.485475506 | 0.686 | 0.703 | 4.78E-20 |
| Pcnp | 7.29E-24 | 0.362700626 | 0.804 | 0.848 | 8.34E-20 |
| Gmfb | 8.05E-24 | 0.296852905 | 0.85 | 0.9 | 9.21E-20 |
| Ids | 9.15E-23 | 0.342915377 | 0.802 | 0.851 | 1.05E-18 |
| Pten | 1.42E-22 | 0.316207102 | 0.868 | 0.902 | 1.62E-18 |
| Cadps | 2.28E-22 | 0.370995223 | 0.79 | 0.844 | 2.61E-18 |
| Arf3 | 2.35E-22 | 0.372751454 | 0.802 | 0.856 | 2.68E-18 |
| Gnai2 | 4.42E-22 | 0.326205952 | 0.873 | 0.91 | 5.05E-18 |
| Epb41l3 | 4.76E-22 | 0.37992027 | 0.75 | 0.79 | 5.45E-18 |
| Ptges3 | 9.72E-22 | 0.30333694 | 0.86 | 0.912 | 1.11E-17 |
| St13 | 2.48E-21 | 0.392786666 | 0.816 | 0.857 | 2.84E-17 |
| Chd3 | 6.76E-21 | 0.378737267 | 0.747 | 0.799 | 7.74E-17 |
| Elavl3 | 2.37E-20 | 0.380303383 | 0.755 | 0.792 | 2.71E-16 |
| Snap91 | 3.12E-20 | 0.349453497 | 0.792 | 0.84 | 3.57E-16 |
| Calm3 | 4.17E-20 | 0.271091781 | 0.86 | 0.914 | 4.77E-16 |
| Asap1 | 5.71E-20 | 0.292618686 | 0.805 | 0.862 | 6.53E-16 |
| Usp9x | 1.29E-19 | 0.354022429 | 0.788 | 0.837 | 1.47E-15 |
| Ncam1 | 1.75E-19 | 0.373075836 | 0.832 | 0.831 | 2.01E-15 |
| Kif5a | 1.75E-19 | 0.379714564 | 0.829 | 0.851 | 2.01E-15 |
| Cbfb | 2.98E-19 | 0.311654326 | 0.822 | 0.871 | 3.41E-15 |
| Rab6b | 3.90E-19 | 0.27752139 | 0.832 | 0.882 | 4.46E-15 |
| Pdpk1 | 7.63E-19 | 0.344756994 | 0.758 | 0.784 | 8.73E-15 |
| Atpaf2 | 1.10E-18 | 0.288133452 | 0.646 | 0.789 | 1.26E-14 |
| Ppp3cb | 1.67E-18 | 0.331846488 | 0.79 | 0.861 | 1.91E-14 |
| Ivns1abp | 1.94E-18 | 0.365377298 | 0.743 | 0.796 | 2.22E-14 |
| Dnm3 | 3.99E-18 | 0.285379595 | 0.826 | 0.857 | 4.57E-14 |
| Fam174a | 5.55E-18 | 0.259846264 | 0.918 | 0.947 | 6.35E-14 |
| Nfe2l1 | 6.31E-18 | 0.377474463 | 0.777 | 0.85 | 7.22E-14 |
| Rcan2 | 8.63E-18 | 0.336076432 | 0.85 | 0.881 | 9.87E-14 |
| Isl1 | 1.07E-17 | 0.2598041 | 0.839 | 0.902 | 1.22E-13 |
| Bhlhe41 | 5.27E-17 | 0.497876208 | 0.565 | 0.474 | 6.03E-13 |
| Pld5 | 1.51E-16 | 0.320954004 | 0.838 | 0.877 | 1.73E-12 |
| Pcp4 | 1.90E-16 | -0.277576049 | 0.549 | 0.704 | 2.18E-12 |
| Csnk1a1 | 2.05E-16 | 0.263717135 | 0.845 | 0.891 | 2.35E-12 |
| Cmip | 2.17E-16 | 0.374023638 | 0.731 | 0.765 | 2.48E-12 |
| Arl8b | 2.45E-16 | 0.365405853 | 0.738 | 0.773 | 2.80E-12 |
| 2-Mar | 3.66E-16 | 0.379570128 | 0.805 | 0.828 | 4.19E-12 |
| Scn9a | 3.92E-16 | 0.253603273 | 0.909 | 0.944 | 4.49E-12 |
| Syt7 | 4.57E-16 | 0.399086763 | 0.851 | 0.895 | 5.23E-12 |
| Syn1 | 6.35E-16 | 0.336315603 | 0.768 | 0.821 | 7.26E-12 |
| Caprin1 | 6.81E-16 | 0.335494408 | 0.743 | 0.804 | 7.79E-12 |
| Zbtb7a | 6.97E-16 | 0.438221647 | 0.746 | 0.825 | 7.97E-12 |
| Mbnl1 | 8.43E-16 | 0.322468485 | 0.835 | 0.853 | 9.64E-12 |
| Ppp2r2c | 1.70E-15 | 0.357745993 | 0.733 | 0.776 | 1.94E-11 |
| Chmp3 | 1.79E-15 | 0.342144581 | 0.764 | 0.81 | 2.05E-11 |
| Hecw1 | 2.37E-15 | 0.356966425 | 0.752 | 0.803 | 2.72E-11 |
| Irf2bpl | 2.72E-15 | 0.442118925 | 0.679 | 0.691 | 3.11E-11 |
| Nrxn2 | 3.60E-15 | 0.345666724 | 0.792 | 0.875 | 4.12E-11 |
| Ppp1cb | 3.80E-15 | 0.308580281 | 0.766 | 0.815 | 4.34E-11 |
| Spop | 9.03E-15 | 0.340589299 | 0.826 | 0.856 | 1.03E-10 |
| Rpl12 | 9.76E-15 | 0.292520126 | 0.875 | 0.911 | 1.12E-10 |
| Eif3h | 1.55E-14 | 0.290243262 | 0.861 | 0.898 | 1.78E-10 |
| Map1a | 1.91E-14 | 0.346236346 | 0.865 | 0.908 | 2.18E-10 |
| H2-T23 | 3.62E-14 | 0.311683506 | 0.516 | 0.683 | 4.14E-10 |
| Taok1 | 1.20E-13 | 0.341443842 | 0.715 | 0.768 | 1.37E-09 |
| Actr2 | 3.55E-13 | 0.355602056 | 0.724 | 0.784 | 4.06E-09 |
| Eif3a | 4.32E-13 | 0.391589856 | 0.711 | 0.769 | 4.94E-09 |
| Ppm1b | 6.19E-13 | 0.313341694 | 0.767 | 0.851 | 7.08E-09 |
| A330069E16Rik | 7.22E-13 | 0.421946584 | 0.504 | 0.66 | 8.26E-09 |
| Matr3 | 1.50E-12 | 0.297355699 | 0.759 | 0.832 | 1.71E-08 |
| Nsf | 2.09E-12 | 0.336992033 | 0.695 | 0.741 | 2.39E-08 |
| Cd44 | 3.84E-12 | 0.308440856 | 0.695 | 0.694 | 4.40E-08 |
| Stau2 | 5.14E-12 | 0.332286819 | 0.727 | 0.791 | 5.88E-08 |
| Spry2 | 6.26E-12 | 0.474109216 | 0.813 | 0.786 | 7.16E-08 |
| Ythdf2 | 6.76E-12 | 0.369273461 | 0.744 | 0.802 | 7.73E-08 |
| Gabbr1 | 9.77E-12 | 0.312632142 | 0.747 | 0.804 | 1.12E-07 |
| Rph3a | 9.81E-12 | 0.361466081 | 0.76 | 0.835 | 1.12E-07 |
| Bmpr2 | 1.17E-11 | 0.297646296 | 0.736 | 0.802 | 1.34E-07 |
| Dgkz | 1.72E-11 | 0.372047043 | 0.743 | 0.835 | 1.96E-07 |
| Paip1 | 2.26E-11 | 0.454429474 | 0.788 | 0.764 | 2.58E-07 |
| Pam | 2.58E-11 | 0.528479032 | 0.837 | 0.917 | 2.95E-07 |
| Cntn1 | 4.88E-11 | 0.365295761 | 0.716 | 0.773 | 5.58E-07 |
| Fbxw7 | 5.26E-11 | 0.426064827 | 0.657 | 0.695 | 6.02E-07 |
| Nav1 | 7.53E-11 | 0.429067597 | 0.64 | 0.664 | 8.61E-07 |
| Purb | 1.63E-10 | 0.334120164 | 0.665 | 0.716 | 1.86E-06 |
| Gabra1 | 1.66E-10 | 0.301372093 | 0.733 | 0.794 | 1.90E-06 |
| Rab10 | 1.80E-10 | 0.281306769 | 0.757 | 0.82 | 2.06E-06 |
| Ano3 | 2.47E-10 | 0.285916803 | 0.825 | 0.874 | 2.83E-06 |
| Flot2 | 3.16E-10 | 0.324464143 | 0.714 | 0.782 | 3.61E-06 |
| Itpkb | 3.68E-10 | 0.333548926 | 0.696 | 0.762 | 4.21E-06 |
| Akap11 | 4.46E-10 | 0.318575146 | 0.718 | 0.773 | 5.11E-06 |
| Rpl22 | 5.94E-10 | 0.300091418 | 0.744 | 0.809 | 6.79E-06 |
| Ntrk3 | 6.18E-10 | 0.269950541 | 0.775 | 0.845 | 7.07E-06 |
| Junb | 7.51E-10 | 0.466607218 | 0.562 | 0.561 | 8.59E-06 |
| Prkacb | 9.13E-10 | 0.351328784 | 0.687 | 0.747 | 1.04E-05 |
| Ptp4a2 | 1.35E-09 | 0.357895166 | 0.644 | 0.689 | 1.55E-05 |
| Vps4b | 1.55E-09 | 0.281025744 | 0.759 | 0.845 | 1.78E-05 |
| Thra | 1.86E-09 | 0.345446497 | 0.743 | 0.83 | 2.13E-05 |
| Xist | 3.56E-09 | 0.473707455 | 0.984 | 0.974 | 4.07E-05 |
| Ncor1 | 3.57E-09 | 0.307248981 | 0.795 | 0.827 | 4.09E-05 |
| Set | 5.20E-09 | 0.265057947 | 0.755 | 0.833 | 5.95E-05 |
| Kdm6b | 7.75E-09 | 0.26063818 | 0.509 | 0.671 | 8.87E-05 |
| Nckap1 | 1.36E-08 | 0.316698732 | 0.707 | 0.775 | 0.000155687 |
| Eif5a2 | 2.33E-08 | 0.299271301 | 0.683 | 0.746 | 0.000267048 |
| Agtpbp1 | 2.58E-08 | 0.330380967 | 0.68 | 0.754 | 0.000295112 |
| Adipor1 | 3.45E-08 | 0.307496339 | 0.808 | 0.841 | 0.000394603 |
| Krit1 | 4.42E-08 | 0.365709563 | 0.63 | 0.689 | 0.000505806 |
| Cd164 | 4.79E-08 | 0.357019974 | 0.671 | 0.72 | 0.000548522 |
| Phactr2 | 2.06E-07 | 0.334274777 | 0.654 | 0.693 | 0.002354962 |
| Stat3 | 2.37E-07 | 0.34360328 | 0.73 | 0.834 | 0.002715962 |
| Larp4b | 2.52E-07 | 0.348344234 | 0.753 | 0.771 | 0.002886985 |
| Mef2a | 2.55E-07 | 0.303596189 | 0.69 | 0.761 | 0.002921088 |
| Adrbk1 | 2.68E-07 | 0.365471375 | 0.709 | 0.805 | 0.00306462 |
| Nedd4l | 2.73E-07 | 0.325495038 | 0.764 | 0.788 | 0.003121745 |
| Fam189b | 2.88E-07 | 0.279702217 | 0.854 | 0.916 | 0.003297763 |
| B4galt6 | 3.03E-07 | 0.373694841 | 0.654 | 0.711 | 0.003463015 |
| Chd5 | 3.96E-07 | 0.389370821 | 0.702 | 0.785 | 0.004529327 |
| Mtmr6 | 4.04E-07 | 0.304377387 | 0.687 | 0.746 | 0.00462373 |
| Dnajc5 | 4.23E-07 | 0.332588912 | 0.655 | 0.702 | 0.004844733 |
| Pou4f2 | 4.90E-07 | 0.256416043 | 0.724 | 0.797 | 0.005607109 |
| Wac | 5.23E-07 | 0.333585347 | 0.621 | 0.674 | 0.005978573 |
| Ppp3r1 | 5.67E-07 | 0.353945163 | 0.656 | 0.712 | 0.006482179 |
| Ctnnb1 | 7.59E-07 | 0.276291725 | 0.67 | 0.732 | 0.0086833 |
| 2900055J20Rik | 8.56E-07 | 0.349732644 | 0.743 | 0.755 | 0.009794354 |
| Msn | 8.84E-07 | 0.277537495 | 0.72 | 0.792 | 0.010118488 |
| Ptbp3 | 9.34E-07 | 0.322013 | 0.629 | 0.685 | 0.010690292 |
| Cpeb2 | 1.34E-06 | 0.314045812 | 0.667 | 0.733 | 0.015310142 |
| Chp1 | 1.59E-06 | 0.298400243 | 0.681 | 0.757 | 0.018138044 |
| Tspyl1 | 1.61E-06 | 0.333715119 | 0.656 | 0.715 | 0.018445324 |
| Tspan2 | 4.09E-06 | 0.281415855 | 0.791 | 0.834 | 0.04678504 |
| Kcnab2 | 5.90E-06 | 0.276991953 | 0.762 | 0.857 | 0.067451285 |
| Lrrc8d | 1.25E-05 | 0.287212752 | 0.66 | 0.746 | 0.143321457 |
| Gls | 1.34E-05 | 0.323948661 | 0.64 | 0.693 | 0.153625543 |
| Usp34 | 1.39E-05 | 0.332082753 | 0.755 | 0.796 | 0.159199511 |
| Nptn | 1.62E-05 | 0.314217794 | 0.651 | 0.727 | 0.185343203 |
| Zswim6 | 1.69E-05 | 0.251692567 | 0.455 | 0.538 | 0.193923822 |
| Vwa5a | 1.90E-05 | 0.265106288 | 0.714 | 0.798 | 0.217202406 |
| Abhd2 | 2.99E-05 | 0.571516428 | 0.695 | 0.796 | 0.342188342 |
| Ube2q2 | 3.04E-05 | 0.345927397 | 0.642 | 0.715 | 0.347703296 |
| Zbtb38 | 3.10E-05 | 0.367998997 | 0.705 | 0.727 | 0.354580385 |
| Marf1 | 3.49E-05 | 0.386248982 | 0.798 | 0.783 | 0.398870737 |
| Pik3ca | 3.99E-05 | 0.328944872 | 0.641 | 0.699 | 0.456072753 |
| Map3k1 | 5.86E-05 | 0.445561139 | 0.548 | 0.578 | 0.670063776 |
| Rtf1 | 5.91E-05 | 0.281983302 | 0.756 | 0.854 | 0.675685617 |
| Mpped2 | 9.91E-05 | 0.275560182 | 0.789 | 0.83 | 1 |
| Rps6ka3 | 9.92E-05 | 0.37275092 | 0.594 | 0.649 | 1 |
| Trim36 | 0.000106152 | 0.288973346 | 0.605 | 0.671 | 1 |
| Trappc10 | 0.000129265 | 0.286900429 | 0.443 | 0.503 | 1 |
| Eno2 | 0.000138014 | 0.287543321 | 0.673 | 0.755 | 1 |
| Robo2 | 0.000141851 | 0.292161465 | 0.757 | 0.795 | 1 |
| Mapt | 0.000146467 | 0.295324796 | 0.681 | 0.748 | 1 |
| Nek7 | 0.000177471 | 0.272365981 | 0.678 | 0.753 | 1 |
| Rbfox3 | 0.000182203 | 0.37052084 | 0.602 | 0.649 | 1 |
| Acbd5 | 0.000241538 | 0.273311472 | 0.673 | 0.756 | 1 |
| Rnf112 | 0.000248732 | 0.2661727 | 0.774 | 0.808 | 1 |
| Ubl3 | 0.000262605 | 0.3245772 | 0.734 | 0.76 | 1 |
| Peli1 | 0.000264426 | 0.389469878 | 0.546 | 0.579 | 1 |
| Pink1 | 0.000353148 | 0.313432428 | 0.76 | 0.792 | 1 |
| Socs2 | 0.000415497 | 0.377682227 | 0.688 | 0.708 | 1 |
| Sptan1 | 0.000460135 | 0.295849444 | 0.654 | 0.716 | 1 |
| Bach2 | 0.000490898 | 0.32385655 | 0.538 | 0.592 | 1 |
| Add1 | 0.000500643 | 0.349448427 | 0.616 | 0.669 | 1 |
| Myef2 | 0.000561047 | 0.393338993 | 0.591 | 0.64 | 1 |
| Clip3 | 0.000596408 | 0.33932663 | 0.617 | 0.681 | 1 |
| Reep1 | 0.000790686 | 0.258309453 | 0.675 | 0.772 | 1 |
| Clasp2 | 0.000883469 | 0.297758191 | 0.572 | 0.729 | 1 |
| Gcnt2 | 0.000954949 | 0.345813974 | 0.594 | 0.65 | 1 |
| Pde1c | 0.001009187 | 0.297646139 | 0.753 | 0.78 | 1 |
| Crk | 0.001175717 | 0.30927043 | 0.61 | 0.699 | 1 |
| Azin1 | 0.001302155 | 0.286001074 | 0.649 | 0.749 | 1 |
| Mapk1 | 0.001344658 | 0.267697438 | 0.665 | 0.753 | 1 |
| Cacnb4 | 0.001354336 | 0.304789158 | 0.64 | 0.724 | 1 |
| Dnm1l | 0.001439999 | 0.256124698 | 0.648 | 0.724 | 1 |
| Brap | 0.00144458 | 0.302774633 | 0.746 | 0.792 | 1 |
| Spata13 | 0.001454453 | 0.293871326 | 0.473 | 0.541 | 1 |
| Ehd3 | 0.00166586 | 0.270830463 | 0.588 | 0.736 | 1 |
| Impad1 | 0.001686701 | 0.291984566 | 0.635 | 0.698 | 1 |
| Dctn4 | 0.001810095 | 0.378865936 | 0.652 | 0.711 | 1 |
| Tusc2 | 0.001846353 | 0.263369625 | 0.799 | 0.849 | 1 |
| Cul3 | 0.001876347 | 0.294869013 | 0.6 | 0.671 | 1 |
| Atp9a | 0.001937496 | 0.309253661 | 0.645 | 0.71 | 1 |
| Cbx3 | 0.002563409 | 0.396380947 | 0.64 | 0.706 | 1 |
| Rtn4rl1 | 0.003580749 | 0.309403323 | 0.716 | 0.744 | 1 |
| Cdk5r2 | 0.004083229 | 0.326778442 | 0.675 | 0.771 | 1 |
| Inpp4a | 0.004241018 | 0.393673167 | 0.659 | 0.777 | 1 |
| C77080 | 0.004467784 | 0.255534155 | 0.583 | 0.723 | 1 |
| Zbtb4 | 0.006111729 | 0.328338614 | 0.696 | 0.711 | 1 |
| Slc25a1 | 0.006741039 | 0.279172934 | 0.721 | 0.827 | 1 |
| Suds3 | 0.00766625 | 0.252683721 | 0.758 | 0.783 | 1 |
| Cbx6 | 0.007742122 | 0.32568952 | 0.585 | 0.639 | 1 |
| Lpin1 | 0.008512583 | 0.32079837 | 0.588 | 0.665 | 1 |
| Lats2 | 0.008740686 | 0.298944591 | 0.578 | 0.657 | 1 |
| Dapk1 | 0.010269423 | 0.337236879 | 0.594 | 0.645 | 1 |
| Atf2 | 0.012487382 | 0.297848751 | 0.704 | 0.73 | 1 |
| Zfos1 | 0.013966359 | 0.300836757 | 0.707 | 0.81 | 1 |
| Sox4 | 0.016215776 | 0.287744597 | 0.426 | 0.483 | 1 |
| Wnk1 | 0.016549172 | 0.283467704 | 0.608 | 0.686 | 1 |
| Tomm70a | 0.018415364 | 0.268941186 | 0.642 | 0.722 | 1 |
| Ttbk2 | 0.01905593 | 0.311536254 | 0.596 | 0.661 | 1 |
| Gatad1 | 0.019441557 | 0.335357547 | 0.685 | 0.792 | 1 |
| Fam63b | 0.01982395 | 0.266279106 | 0.622 | 0.715 | 1 |
| Tiparp | 0.019879972 | 0.271819336 | 0.408 | 0.474 | 1 |
| Capza1 | 0.0213303 | 0.285237051 | 0.619 | 0.703 | 1 |
| Cyfip2 | 0.023592022 | 0.270654469 | 0.641 | 0.716 | 1 |
| Agfg1 | 0.024486452 | 0.302503736 | 0.579 | 0.668 | 1 |
| Id2 | 0.027169844 | 0.258656282 | 0.752 | 0.802 | 1 |
| Camsap2 | 0.027777079 | 0.324486681 | 0.596 | 0.668 | 1 |
| Prkab2 | 0.031364194 | 0.323593711 | 0.721 | 0.751 | 1 |
| Slc24a5 | 0.031830571 | 0.371494095 | 0.695 | 0.674 | 1 |
| Smad1 | 0.032274586 | 0.279779521 | 0.63 | 0.725 | 1 |
| BC030336 | 0.034104857 | 0.284518994 | 0.629 | 0.715 | 1 |
| 3-Sep | 0.034872913 | 0.322446407 | 0.584 | 0.659 | 1 |
| Pygb | 0.037834906 | 0.332726567 | 0.585 | 0.645 | 1 |
| Map4k4 | 0.03866663 | 0.263669821 | 0.51 | 0.571 | 1 |
| Cdk16 | 0.038683547 | 0.323082047 | 0.588 | 0.656 | 1 |
| Brd2 | 0.039944578 | 0.260362212 | 0.515 | 0.596 | 1 |
| Slc24a2 | 0.044828796 | 0.275390675 | 0.708 | 0.814 | 1 |
| Prrxl1 | 0.044926751 | 0.265849715 | 0.836 | 0.919 | 1 |
| Med13l | 0.048899616 | 0.371446373 | 0.513 | 0.562 | 1 |
| Coro2a | 0.050702289 | 0.285160061 | 0.564 | 0.616 | 1 |
| Dync1li2 | 0.053406479 | 0.278290965 | 0.615 | 0.689 | 1 |
| Cpeb4 | 0.054232464 | 0.355604893 | 0.545 | 0.603 | 1 |
| Sobp | 0.063137077 | 0.272675476 | 0.476 | 0.555 | 1 |
| Cacnb3 | 0.067742725 | 0.264646805 | 0.733 | 0.8 | 1 |
| Actb | 0.068207883 | 0.420723551 | 0.999 | 0.999 | 1 |
| Abca1 | 0.0683115 | 0.272705843 | 0.54 | 0.614 | 1 |
| Pak3 | 0.068484405 | 0.26651826 | 0.658 | 0.745 | 1 |
| Myo5a | 0.069089547 | 0.265167196 | 0.606 | 0.686 | 1 |
| Rhoq | 0.072142773 | 0.306188521 | 0.579 | 0.652 | 1 |
| Arid1a | 0.075126844 | 0.261265737 | 0.605 | 0.676 | 1 |
| Kat7 | 0.084075524 | 0.257128537 | 0.531 | 0.608 | 1 |
| Npepps | 0.085332137 | 0.303342243 | 0.554 | 0.616 | 1 |
| Rad21 | 0.085786427 | 0.279655533 | 0.701 | 0.75 | 1 |
| Igsf3 | 0.089477567 | 0.351515213 | 0.647 | 0.67 | 1 |
| Susd6 | 0.090585961 | 0.267487 | 0.523 | 0.591 | 1 |
| Pdzd8 | 0.091441822 | 0.269621122 | 0.51 | 0.613 | 1 |
| Fam168a | 0.093175805 | 0.253605742 | 0.523 | 0.617 | 1 |
| Akap6 | 0.093658681 | 0.263265787 | 0.608 | 0.693 | 1 |
| Plekha6 | 0.095124343 | 0.255738429 | 0.48 | 0.559 | 1 |
| Mfhas1 | 0.096823169 | 0.297060316 | 0.519 | 0.6 | 1 |
| Dock9 | 0.099444484 | 0.287289187 | 0.482 | 0.565 | 1 |
| Asic2 | 0.102606901 | 0.265966336 | 0.628 | 0.708 | 1 |
| Efna5 | 0.107301041 | 0.281229897 | 0.495 | 0.574 | 1 |
| Wtap | 0.123628198 | 0.263667485 | 0.638 | 0.737 | 1 |
| Kcnb2 | 0.124752777 | 0.275713112 | 0.62 | 0.718 | 1 |
| Tnrc6a | 0.132360494 | 0.28502499 | 0.611 | 0.749 | 1 |
| Ctif | 0.135274933 | 0.290300831 | 0.518 | 0.594 | 1 |
| Rgs8 | 0.136946497 | 0.258122561 | 0.659 | 0.698 | 1 |
| Dnajc6 | 0.138494723 | 0.258542155 | 0.53 | 0.613 | 1 |
| Tnrc6c | 0.139595856 | 0.299294587 | 0.541 | 0.622 | 1 |
| Rims1 | 0.139815276 | 0.253576417 | 0.565 | 0.681 | 1 |
| Dnm1 | 0.14089488 | 0.255152854 | 0.572 | 0.66 | 1 |
| Wipf3 | 0.141093403 | 0.281298387 | 0.529 | 0.613 | 1 |
| Plcxd2 | 0.143815178 | 0.266102414 | 0.604 | 0.69 | 1 |
| Mtss1 | 0.144552828 | 0.256869737 | 0.53 | 0.604 | 1 |
| Ugcg | 0.148552808 | 0.26136601 | 0.613 | 0.7 | 1 |
| Ppm1l | 0.148746735 | 0.257402691 | 0.588 | 0.674 | 1 |
| Dusp5 | 0.149644285 | 0.267753529 | 0.555 | 0.63 | 1 |
| Shoc2 | 0.152709144 | 0.309642421 | 0.563 | 0.637 | 1 |
| Usp31 | 0.153137549 | 0.274141056 | 0.626 | 0.72 | 1 |
| Bptf | 0.158587433 | 0.263392339 | 0.5 | 0.587 | 1 |
| Usp22 | 0.159722626 | 0.261361994 | 0.553 | 0.631 | 1 |
| Hdgf | 0.162645972 | 0.263638289 | 0.638 | 0.729 | 1 |
| Mesdc1 | 0.170316052 | 0.255750588 | 0.765 | 0.858 | 1 |
| Uhrf1bp1l | 0.173463975 | 0.2988279 | 0.567 | 0.641 | 1 |
| 2510009E07Rik | 0.18283533 | 0.250455301 | 0.615 | 0.711 | 1 |
| Plekhm3 | 0.202737138 | 0.286107747 | 0.514 | 0.577 | 1 |
| Mycbp2 | 0.234656597 | 0.265489967 | 0.583 | 0.666 | 1 |
| Gnl3l | 0.241340226 | 0.258071662 | 0.601 | 0.67 | 1 |
| Setd7 | 0.250273708 | 0.275310277 | 0.567 | 0.632 | 1 |
| Pim1 | 0.26306423 | 0.372459421 | 0.551 | 0.681 | 1 |
| Dok4 | 0.263382502 | 0.254315739 | 0.808 | 0.9 | 1 |
| Mapk8ip2 | 0.275277673 | 0.250536504 | 0.553 | 0.628 | 1 |
| Onecut2 | 0.291365696 | 0.283649086 | 0.609 | 0.684 | 1 |
| Lrrc8a | 0.299291428 | 0.362564075 | 0.634 | 0.76 | 1 |
| Fat3 | 0.307752279 | 0.267321326 | 0.593 | 0.674 | 1 |
| Strn3 | 0.321063561 | 0.271158906 | 0.565 | 0.665 | 1 |
| Bean1 | 0.32171384 | 0.307803455 | 0.677 | 0.7 | 1 |
| Cul4a | 0.322429413 | 0.302847206 | 0.546 | 0.613 | 1 |
| Rap1gds1 | 0.338262912 | 0.266516978 | 0.595 | 0.691 | 1 |
| Hif1a | 0.343705603 | 0.255721575 | 0.58 | 0.67 | 1 |
| Mapre2 | 0.34765003 | 0.26440257 | 0.58 | 0.67 | 1 |
| Jup | 0.369686845 | 0.272656435 | 0.596 | 0.686 | 1 |
| Zfhx4 | 0.375330369 | 0.295613582 | 0.68 | 0.703 | 1 |
| Amph | 0.387360195 | 0.290153391 | 0.525 | 0.594 | 1 |
| Jph4 | 0.399024042 | 0.252853338 | 0.651 | 0.785 | 1 |
| Foxp1 | 0.39922267 | 0.29813585 | 0.54 | 0.614 | 1 |
| Yme1l1 | 0.401273863 | 0.265069793 | 0.515 | 0.635 | 1 |
| Hcn1 | 0.409283816 | 0.255044405 | 0.511 | 0.593 | 1 |
| Ablim1 | 0.418784609 | 0.265004513 | 0.533 | 0.613 | 1 |
| Birc6 | 0.422342433 | 0.282842361 | 0.49 | 0.567 | 1 |
| Avl9 | 0.427917862 | 0.274172604 | 0.532 | 0.617 | 1 |
| Prkar1b | 0.437657212 | 0.278368482 | 0.602 | 0.696 | 1 |
| Trim37 | 0.457853156 | 0.309435551 | 0.538 | 0.611 | 1 |
| Apba1 | 0.458955556 | 0.273553626 | 0.63 | 0.763 | 1 |
| Tshz3 | 0.465396222 | 0.344946856 | 0.529 | 0.569 | 1 |
| Kmt2c | 0.475673522 | 0.27982774 | 0.485 | 0.563 | 1 |
| Kpna1 | 0.505812725 | 0.264359397 | 0.52 | 0.604 | 1 |
| Rnf44 | 0.517306978 | 0.320881257 | 0.557 | 0.6 | 1 |
| Eif4g3 | 0.539203165 | 0.253661914 | 0.686 | 0.728 | 1 |
| Efr3a | 0.549694801 | 0.268269923 | 0.556 | 0.625 | 1 |
| Ube2o | 0.550048818 | 0.296047229 | 0.599 | 0.73 | 1 |
| Chd4 | 0.564471216 | 0.340968281 | 0.607 | 0.727 | 1 |
| Tmf1 | 0.574197044 | 0.30865623 | 0.702 | 0.701 | 1 |
| Tex2 | 0.578779781 | 0.282492292 | 0.574 | 0.666 | 1 |
| Spin1 | 0.580769855 | 0.257957972 | 0.585 | 0.686 | 1 |
| Mprip | 0.609838071 | 0.27366276 | 0.547 | 0.623 | 1 |
| Nhs | 0.612997098 | 0.292316238 | 0.492 | 0.565 | 1 |
| Socs3 | 0.619813431 | 0.296278975 | 0.609 | 0.612 | 1 |
| Xiap | 0.63722398 | 0.25372242 | 0.591 | 0.682 | 1 |
| Clstn1 | 0.659725139 | 0.255390518 | 0.562 | 0.642 | 1 |
| Sox11 | 0.663230304 | 0.413366912 | 0.475 | 0.502 | 1 |
| Car10 | 0.68662445 | 0.268139633 | 0.671 | 0.713 | 1 |
| Zmiz1 | 0.689061728 | 0.261066136 | 0.543 | 0.623 | 1 |
| Ptprf | 0.692066541 | 0.31665755 | 0.63 | 0.773 | 1 |
| Egr1 | 0.701006738 | 0.263031759 | 0.458 | 0.498 | 1 |
| Lingo1 | 0.704095011 | 0.255908877 | 0.69 | 0.733 | 1 |
| Rab11fip2 | 0.715695406 | 0.25719229 | 0.685 | 0.813 | 1 |
| Wasf3 | 0.727510828 | 0.264193527 | 0.539 | 0.625 | 1 |
| Dmxl2 | 0.731349486 | 0.268497446 | 0.546 | 0.63 | 1 |
| Frat2 | 0.750671934 | 0.322886285 | 0.466 | 0.517 | 1 |
| Ubr3 | 0.792928211 | 0.26625246 | 0.565 | 0.657 | 1 |
| Six1 | 0.80838172 | 0.266414603 | 0.519 | 0.614 | 1 |
| Tmem117 | 0.814583096 | 0.253395712 | 0.551 | 0.646 | 1 |
| Rps27rt | 0.824351636 | 0.298290083 | 0.654 | 0.772 | 1 |
| Mt1 | 0.825186512 | 0.354681918 | 0.515 | 0.552 | 1 |
| Twf1 | 0.826663301 | 0.282025838 | 0.534 | 0.633 | 1 |
| Sorbs2 | 0.828084271 | 0.254646072 | 0.661 | 0.704 | 1 |
| Lrrc75b | 0.858755502 | 0.275697893 | 0.669 | 0.706 | 1 |
| Samd14 | 0.864498957 | 0.287399098 | 0.743 | 0.835 | 1 |
| Synj1 | 0.877086084 | 0.306726623 | 0.515 | 0.584 | 1 |
| Crebrf | 0.879358654 | 0.271032998 | 0.695 | 0.729 | 1 |
| Plekha3 | 0.883491689 | 0.269136598 | 0.517 | 0.613 | 1 |
| Atxn1 | 0.88706628 | 0.299046646 | 0.508 | 0.577 | 1 |
| Apc | 0.897794048 | 0.255684466 | 0.56 | 0.65 | 1 |
| Pum2 | 0.905432307 | 0.252628603 | 0.55 | 0.636 | 1 |
| Picalm | 0.908616541 | 0.253333298 | 0.671 | 0.721 | 1 |
| Sv2a | 0.963711169 | 0.324411526 | 0.527 | 0.614 | 1 |
| Alkbh5 | 0.967967164 | 0.298716621 | 0.676 | 0.671 | 1 |
